# Single-Cell Perturbations Reveal Selective Modulation of Causal Connectivity During Decision-Making

**DOI:** 10.64898/2026.04.07.716761

**Authors:** Mark L. Ioffe, Stephan Thiberge, Carlos Brody, David W. Tank

## Abstract

How does cortical connectivity support decision-making? Behavioral tasks often involve multiple sequential phases that implement different computations. For perceptual decision-making during navigation these can include evidence accumulation, decision commitment, and motor program read out. How are these different phases implemented in circuits with fixed anatomical synaptic connectivity? One potential contribution is that the connectivity of neurons is modulated in the different phases, but this has never been tested. Here we used an all-optical method to probe the causal connectivity of excitatory neurons in layer 2/3 of mouse retrosplenial cortex during different behavioral epochs of a navigation-based decision-making task, as well as in the absence of the task. In-task connectivity was different from no-task connectivity: furthermore, these differences were selective to the cue / decision phase, tapering off in later stages of the task. We propose that fast modulation of connectivity is a prevalent mechanism in neural circuit function.

## INTRODUCTION

Neuronal dynamics are shaped by circuit connectivity. The causal connectivity – defined here as the sign and magnitude of a neuron’s response to the causal optogenetic activation of another neuron – that governs signal propagation in vivo need not be identical to the physical synaptic connectivity, opening up the possibility that the causal connectivity is strongly modulated by ongoing task demands. Whether it is modulated by task demands has not been explicitly probed. Thus, it remains unclear (i) if connectivity is modulated by the task, (ii) if connectivity is modulated dynamically within the task, and (iii) what the magnitude of modulation is. Here we explore task-dependent changes in the causal connectivity of cortical layer 2/3 circuits during a navigation-based decision-making task in a virtual reality (VR) maze, with an all-optical single-cell perturbation protocol.

All-optical simultaneous imaging and stimulation techniques have been in use in awake, behaving animals for over a decade^1–10^, and have only infrequently been used to map the local circuit’s causal connectivity with single-cell activations in vivo^11–13^. Our work builds on a statistical averaging approach termed “influence” (originally defined by Chettih & Harvey^11^) to experimentally characterize the weaker interactions at larger separation distances, which constitute the majority of measured interactions. The causal interaction between a specific pair of neurons can be quantified using the term ΔActivity, which captures the effect of optogenetic activation of a “target” neuron on the magnitude of calcium-dependent fluorescence changes of a “responder” neuron, normalized to the range of responses in the responder neuron. At larger separation distances, these interactions are frequently mediated multi-synaptically, which suggests that the typical magnitude of interactions will be small. Indeed, Chettih & Harvey found that measuring the ΔActivity for a typical pair of neurons reliably would require prohibitively large numbers of repeat optogenetic stimulation. Instead, they searched for trends in the influence, defined as the average of ΔActivity over many (∼10^4) different target-responder pairs. Here we extend their approach with the additional component of examining influence at different phases of the behavior. Our experimental strategy was to constrain target stimulation to given time points during animal traversal of the maze, to estimate how ΔActivity depended on the underlying time-varying computations. Additionally, in a subset of sessions, the same targets were perturbed in a task-independent condition. The novelty in our experimental approach is that the connectivity between a specific pair of neurons can be compared across upcoming decisions (to go right or left), or in the absence/presence of the task, to tease out computation-dependent shifts in the causal connectivity (Fig. 1A, left panel).

**Figure 1.**
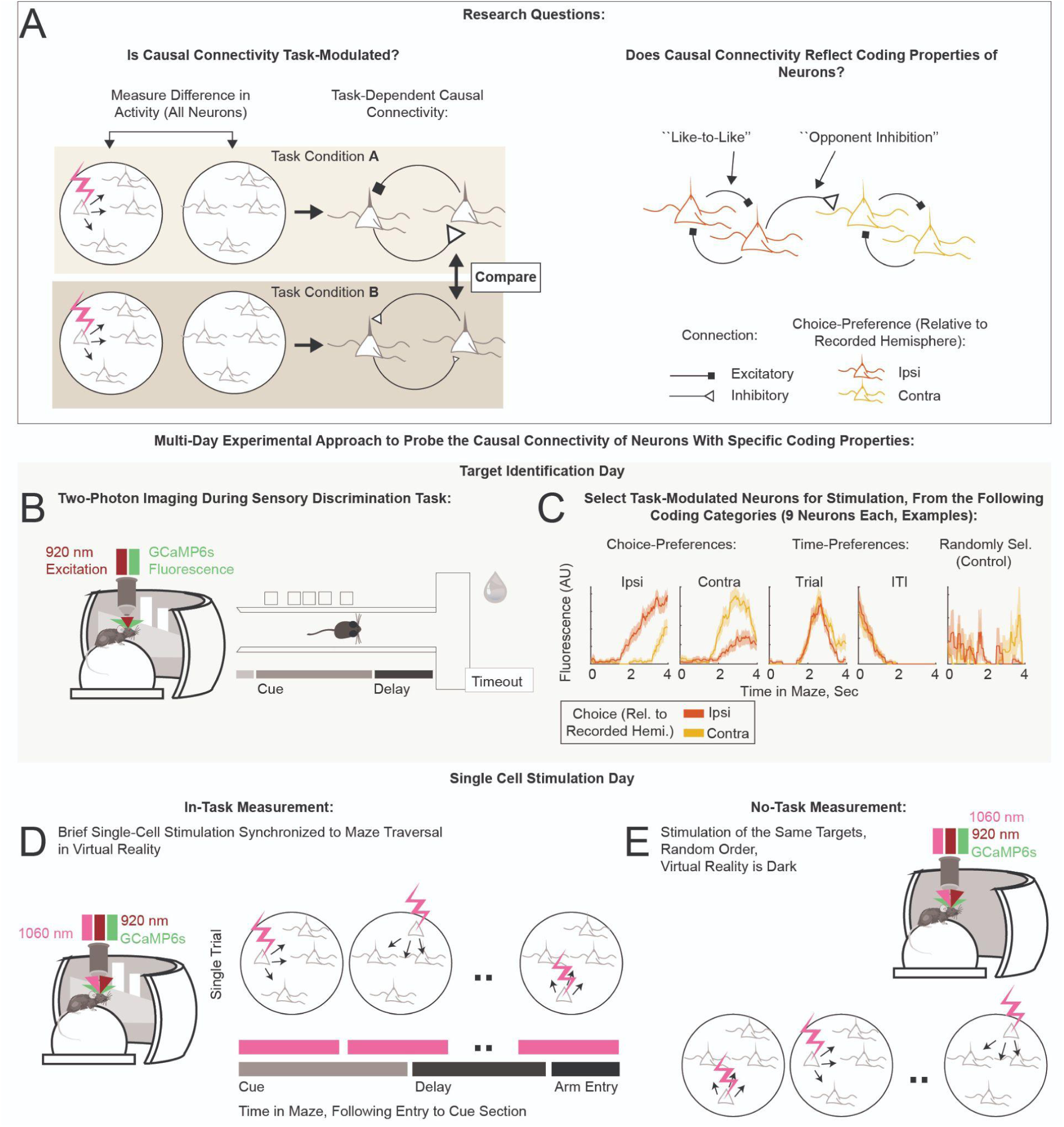
Overview. **A.** Research questions. **B–E**. Schematics of the experimental approach. All-optical simultaneous imaging and single-cell stimulation experiments probed the causal connectivity in cortical circuits, while mice engaged in a virtual reality task. Technical details of the experiment can be found in Fig. S1. **B,C**. Identification of target neurons for subsequent stimulation, based on their task-encoding properties. **B.** Calcium imaging of neural activity during behavior. **C.** Fluorescence activity of example neurons selected for stimulation. Shaded areas indicate standard error (STE). Typical encoding properties of the neurons selected for stimulation can be found in Fig. S2. **D,E**. Single-cell stimulation experiments. **D.** In-task perturbations were synchronized to virtual reality maze traversal (described in Fig. 2). **E.** No-task perturbations were performed immediately following the in-task measurement, in a subset of sessions (11 of 20). Mice remained head-fixed on the spherical treadmill and the virtual reality screen was dark. The same target neurons were stimulated as in the in-task recording (described in Fig. S3).

We performed all-optical experiments on excitatory neurons in layer 2/3 retrosplenial cortex (RSC) in mice trained on the accumulating towers task (ATT)^14^. As mice perform this task, neural activity within the RSC follows choice-specific trajectories^15–17^. We were interested in whether local connectivity motifs (specifically, “like-to-like” excitation between neurons with similar choice-preferences, or “opponent inhibition” between neurons with opposing choice-preferences, Fig. 1A right panel) underlie this neural correlate of decision-making. We chose RSC specifically because of its known involvement in the ATT^18^, and because firing fields of RSC neurons tile ATT maze traversal^15,16^.

We report that task-dependent modulations of causal connectivity peaked near the time of decision formation, fading away to zero by the time of choice. This demonstrates that causal connectivity is modulated selectively by ongoing computation. Separately, in-task connectivity was modulated by the identity of the upcoming choice. This modulation occurred during decision formation, but not subsequently, suggesting that connectivity modulation contributes to changes in pre- versus post-decision commitment neuronal dynamics.

## RESULTS

Our all-optical mapping approach is based on optically activating neurons using two-photon excitation of an opsin and simultaneously measuring neural responses using two-photon imaging of a calcium indicator^1,2^. We used a bicistronic viral expression strategy (similar to Refs. ^5,19^) to co-express the calcium indicator GCaMP6s and the soma-localized opsin ChrimsonR in excitatory neurons of C57BL/6 mice using the CamKiiα promoter (Fig. S1A,B). Following surgery and acclimation to a water-restricted regimen, mice were trained on the ATT maze in VR^14^ (Methods): a T-shaped maze where bright towers appeared on the right- and left-hand sides as mice ran down the stem of the maze (“Cue” section, Fig. 1B). The subsequent “Delay” section of the stem required the mice to maintain their decision in the absence of cues. Following the Delay, a turn into the arm of the “T” with more cues yielded a liquid reward. Incorrect choices resulted in an aversive sound and increased the inter-trial-interval (ITI) before the start of the next trial. Mice were trained until they reached the final maze difficulty, or performance plateaued (Methods). Single-cell perturbation experiments were performed near the end of the training period, in a version of the task lacking distractors (Fig. S1C). The no-distractors task was easy enough for the mice to perform well (correct performance: median, 88%; range, 72–96%; 9 mice, 20 sessions; Fig. S1D), allowing many repeat measurements of the response to perturbation, yet complex enough to test for modulations of the connectivity during decision-making.

### Perturbation Protocols

Our goal was to probe the causal connectivity of neurons with specific task-encoding properties (Fig. 1A). Because many perturbations are required to provide statistical significance, our strategy was to first have an imaging-only target identification session to identify neurons belonging to specific coding classes (Fig. 1B,C). Perturbation experiments were then performed on a subsequent day (Fig. 1D,E). On the target identification day, we volumetrically imaged activity in layer 2/3 (range of depths: 120–350 μm below the coverslip; volume: five 680 μm X 680 μm planes 25 μm apart; Fig. S1H) while the mice performed the ATT. Regions of interest (ROI) identified by Suite2p^20^ (number of ROIs that passed a set of anatomical and statistical criteria on target identification sessions: median, 924; range, 636–1225; Methods) were sorted by the magnitude of their preference for the two upcoming choices (“Ipsi”, “Contra”) and one of two time periods of interest (“Trial”, “ITI”). Up to 9 of the most selective neurons in each of these four target classes were recruited for perturbations. Additionally, 9 or more neurons were recruited into a target class that consisted of randomly chosen neurons, independent of coding properties (“Control”; Fig. 1C; Methods). Consistency of target coding across days, and an analysis of the encoding properties of target classes, can be found in supplementary panels Fig. S1E,F and Fig. S2. This selection process resulted in 45 neurons per session that were targeted for stimulation and included in the subsequent analyses.

On the subsequent stimulation day, we randomly chose 80% of the trials as stimulation trials. On stimulation trials, target neurons were stimulated one-at-a-time following animal entry into the Cue section of the maze (Fig. 2A). Photostimulation consisted of scanning the stimulation laser beam over the target cell soma for a total of 250 ms, using a ∼17 μm diameter scan (∼7 μm FWHM axial depth, Fig. S1G–I, Methods). We chose to randomize the stimulation times of targets with respect to coding classes, firing fields, and time within maze traversal. For each target, we also kept constant the time within trial at which it was stimulated, to sample repeat stimulations of a given target at a fixed time within trial. Before the session started, each of the 45 target neurons was randomly assigned, independent of its coding class or firing field, to one of 18 sequential 266 ms-wide stimulation bins; these assignments were fixed for the duration of the session (Fig. 2A,B). Stimulation time bins spanned 4.8 seconds, covering the typical time mice spent traversing the stem of the maze. Each time bin had three target neurons assigned to it (e.g., targets Q1–Q3 were assigned to the 17th time bin in Fig. 2A). On each stimulation trial, a time bin was selected to have a target stimulated within it with 75% probability; the target to be stimulated was then chosen randomly from the three targets assigned to that time bin (e.g., Q1, Q2, and Q3 are distributed across trials in Fig. 2A). The remaining 25% of stimulation trials, within which no target was stimulated in the stimulation bin, provided the crucial baseline measurement of responder activity. These comparison trials were matched to the target-specific stimulation trials in the statistics of stimulation of other targets in other stimulation bins. The causal connectivity, i.e. the change in responder neuron activity due to stimulation of target Q1, was estimated by comparing responder neuron activity on “Q1-Stimulation” (gray, Fig. 2A,D–F) and “Q1-Comparison” (beige, Fig. 2A,D–F) trials. Thus this in-task measure of causal connectivity captures the effect of stimulating a single neuron at a fixed time in the maze, against a background of randomized stimulation of other targets at other times.

**Figure 2.**
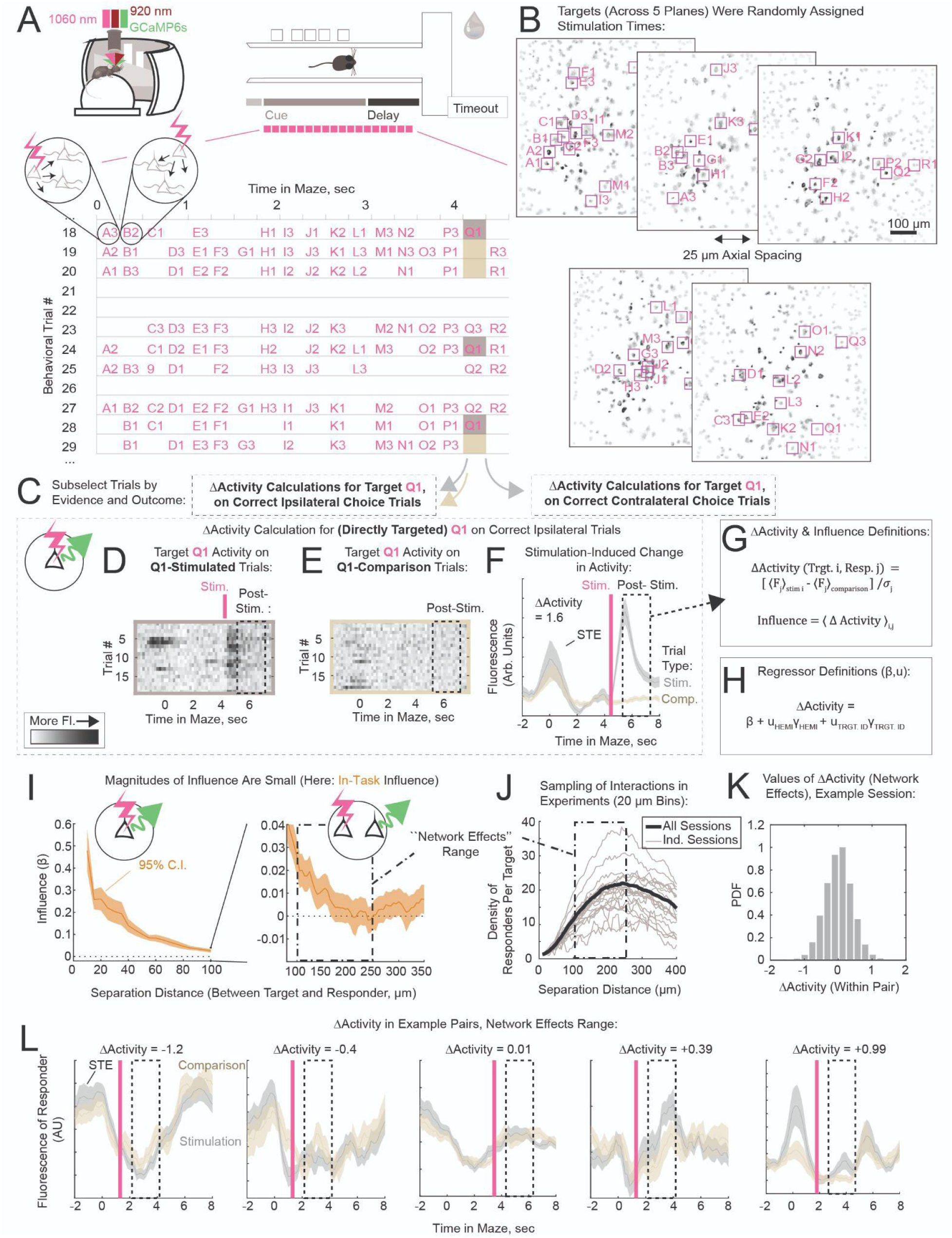
Measuring In-Task Causal Connectivity. **A.** In-task stimulation design. In-task stimulation sequences (examples shown in table) were triggered following animal entry into the Cue section of the maze. For each behavioral trial (row), the target ID stimulated (magenta; e.g., A3) is indicated within each of the 18 sequential stimulation bins (columns, target letter indicates bin). As an example, calculation of influence from target Q1 (bold magenta) is shown. The activity of a responder is evaluated on trials within which Q1 was stimulated (“Stimulation Trials", dark gray) and separately on stimulation trials within which no Q target was stimulated (“Comparison Trials”, beige). Note that there are other targets stimulated in other stimulation time bins (letters A–P, R) during stimulation and comparison trials. Thus the estimate of causal connectivity between a specific target and responding neuron, ΔActivity, was estimated on a background of non-specific stimulation of other targets in other stimulation time bins. Fluorescence activity on non-stimulation trials (here: rows 21, 22 and 26) was withheld to estimate task-encoding, and did not contribute to estimates of ΔActivity. **B.** Anatomical locations of targets selected for stimulation, in an example dataset (shown: Suite2p-extracted ROIs, with magenta boxes and labels indicating targets). Targets were randomly assigned a time of stimulation (letter) for the duration of the session. **C.** Causal connectivity (ΔActivity) was evaluated separately in the two choice conditions. **D–F**. For clarity, the calculation of ΔActivity for target Q1 on ipsilateral trials is illustrated using a strongly activated responder (direct target, within 20 μm of the stimulation point), as the SNR for individual target-responder pairs was typically too low to observe consistent across-trial responses for a particular pair. **D,E.** Fluorescence activity of directly activated neuron on Q1-Stimulation (**D**) and Q1-Comparison (**E**) trials (darker = more fluorescence). Q1 post-stimulation period indicated with dashed line box. **F.** Activity of the neuron shown in panels D,E; trial-averaged within stimulation and comparison trials (shaded area = STE, 1-second rolling bin). **G.** Definitions of in-task ΔActivity, and influence. **H.** Definitions of the regressors estimated by mixed-effects modelling, related to panel I (Model 1.0, Table 1). We used mixed-effects models for most statistical analyses in this paper (Methods). When estimates of the fixed-effects coefficients (*β*) are plotted, shaded areas indicate the 95% confidence interval (C.I.) returned by the model. Throughout this paper, fixed-effects are indicated by *β* and *x*; random-effects are indicated by *u* and *γ*. **I.** The trend in in-task influence vs. separation distance between stimulation point and responder. ΔActivity was pooled across target-responder pairs (grouped by separation distance; 20 μm rolling bin; both choices), and used to estimate the regressor *β* (defined in panel H). In this mixed-effects model, *β* is the fixed-effects coefficient that remains after the variability captured by the random-effects coefficients has been accounted for; it is the corrected population average over ΔActivity, i.e. influence. The “network effects” range of separations is indicated with a dashed-dot box. **J.** Responder density per target, in 20 μm bins. **K.** ΔActivity values in the network effects range, in an example session (54 targets, 6724 pairs). **L.** ΔActivity examples (value indicated), shown as the trial-averaged activity in the responder on stimulation and comparison trials (1-second rolling bin).

**Table 1.** Mixed-Effects Models.

| Model Name | Figure panels | Data Constraints | Formula | Primary Random-Effects Term (n=) | Targets (n=) | Target-Responder pairs (n=) |
| --- | --- | --- | --- | --- | --- | --- |
| 1.0 | 2i, 3d | All in-task measurements of influence (20 sessions); target-responder pairs that were sufficiently sampled in both ipsilateral and contralateral trials. Grouped by distance (20um rolling bin). | $\Delta\text{Activity} = \beta + u_{\text{HEMI}} \gamma_{\text{HEMI}} + u_{\text{TRGT.ID}} \gamma_{\text{TRGT.ID}}$ | Hemisphere (11) | 590 | 160 076 (0–360 um) |
| 1.1 | 3c | All no-task measurements of influence (11 sessions) | $\Delta\text{Activity} = \beta + u_{\text{HEMI}} \gamma_{\text{HEMI}} + u_{\text{TRGT.ID}} \gamma_{\text{TRGT.ID}}$ | Hemisphere (8) | 495 | 116 542 |
| 1.2 | 3f–k | All target-responder pairs that were sufficiently sampled in no-task, ipsilateral and contralateral trials.<br>The three experimental conditions were included together in one fit. | $\Delta\text{Activity} = \beta_{\text{NO-TASK}} + \beta_{\text{IPSI}} X_{\text{IPSI}} + \beta_{\text{CONTRA}} X_{\text{CONTRA}} + u_{\text{HEMI}} \gamma_{\text{HEMI}} + u_{\text{TRGT.ID}} \gamma_{\text{TRGT.ID}}$ | Hemisphere (8) | 389 | 142 034 (Net. Eff.: 53 662) |
| 1.3 | s5c–e | Same data as Model 1.2.<br>The three experimental conditions were fit separately. Grouped by distance (50um bin) | $\Delta\text{Activity} = \beta + u_{\text{HEMI}} \gamma_{\text{HEMI}} + u_{\text{TRGT.ID}} \gamma_{\text{TRGT.ID}}$ | " | " | " |
| 1.4 | s5l–n | Same data as Model 1.2; analysis was done on the binarized $\Delta\text{Activity}$ (+1 / -1).<br>The three experimental conditions were included together in one fit. | $\text{Binarized } \Delta\text{Activity} = \beta_{\text{NO-TASK}} + \beta_{\text{IPSI}} X_{\text{IPSI}} + \beta_{\text{CONTRA}} X_{\text{CONTRA}} + u_{\text{HEMI}} \gamma_{\text{HEMI}} + u_{\text{TRGT.ID}} \gamma_{\text{TRGT.ID}}$ | " | " | " |
| 1.5 | s5o–q | Same data as Model 1.2; analysis was done on the binarized $\Delta\text{Activity}$ (+1 / -1).<br>The three experimental conditions were fit separately. | $\text{Binarized } \Delta\text{Activity} = \beta + u_{\text{HEMI}} \gamma_{\text{HEMI}} + u_{\text{TRGT.ID}} \gamma_{\text{TRGT.ID}}$ | " | " | " |
| 2.0 | s6d–i | This analysis was performed on all task-dependent sessions (20); but not all sessions are sampled at the lower evidence-thresholds.<br>Network effects; Evidence threshold = 1.5; Sampling is shown for Ipsi / Contra comparisons.<br>Evidence-dependent conditions (ambiguous and more-evidence) were fit separately. | $\Delta\text{Activity} = \beta + u_{\text{SESH.ID}} \gamma_{\text{SESH.ID}} + u_{\text{TRGT.ID}} \gamma_{\text{TRGT.ID}}$ | Sessions (17 / 18) | 47/52 | 789/697 |
| 2.1 | s6j–m | Same data as Model 2.0.<br>Evidence-dependent conditions were included together in one fit. | $\Delta\text{Activity} = \beta_{\text{AMB}} + \beta_{\text{EV}} X_{\text{EV}} + u_{\text{SESH.ID}} \gamma_{\text{SESH.ID}} + u_{\text{TRGT.ID}} \gamma_{\text{TRGT.ID}}$ | " | " | " |
| 3.0 | 4c–e | These target-responder pairs were sampled in no-task and both task-dependent conditions (per epoch).<br>Network effects range interactions (100–250um separation). | $\Delta\text{Activity} = \beta_{\text{NO-TASK}} + \beta_{\text{EPOCH / CHOICE}} X_{\text{EPOCH / CHOICE}} + u_{\text{HEMI}} \gamma_{\text{HEMI}} + u_{\text{TRGT.ID}} \gamma_{\text{TRGT.ID}}$ | Hemisphere (8) | 300 | 43 883 |
| 3.1 | 4i–m, s7l–q | Same data as Model 3.0. | $\Delta\text{Activity} = \beta_{\text{NO TASK}} + \beta_{\text{EPOCH / CHOICE}} X_{\text{EPOCH / CHOICE}} + \beta_{\text{TRGT.CLASS}} X_{\text{TRGT.CLASS}} + \beta_{\text{TRGT.CLASS : EPOCH / CHOICE}} X_{\text{TRGT.CLASS : EPOCH / CHOICE}} + u_{\text{HEMI}} \gamma_{\text{HEMI}}$ | " | " | " |
| 4.0 | 5d–f | All in-task sessions (20). Target-responder pairs were sufficiently sampled in both task-dependent conditions. Network effects range. Epochs analyzed separately. | Choice-Dep. Change in $\Delta\text{Activity} = \beta_{\text{CONTROL TRGT}} + \beta_{\text{TRGT. CLASS}} X_{\text{TRGT. CLASS}} + u_{\text{HEMI}} \gamma_{\text{HEMI}}$ | Hemisphere (11) | 590 | 81 734 |
| 4.1 | s11a | Same data as Model 4.0.<br>Choice conditions were analyzed separately. | $\Delta\text{Activity} = \beta + u_{\text{HEMI}} \gamma_{\text{HEMI}}$ | " | " | " |
| 4.2 | 6a, s11b | Same data as Model 4.0.<br>Choice conditions were analyzed separately. | $\Delta\text{Activity} = \beta_{\text{CONTROL TRGT}} + \beta_{\text{TRGT. CLASS}} X_{\text{TRGT. CLASS}} + u_{\text{HEMI}} \gamma_{\text{HEMI}}$ | " | " | " |
| 4.3 | main text (intro to figure 5) | Same data as Model 4.0.<br>This model probed whether coding similarity was relevant as a feature (compared to target class identity) | Choice-Dep. Change in $\Delta\text{Activity} = \beta_{\text{CONTROL TRGT}} + \beta_{\text{TRGT. CLASS}} X_{\text{TRGT. CLASS}} + \beta_{\text{CODING SIMILARITY}} X_{\text{CODING SIMILARITY}} + u_{\text{HEMI}} \gamma_{\text{HEMI}}$ | " | " | " |

In a subset of sessions, our perturbations during the ATT task were followed with a “no-task” single-cell stimulation protocol, within which the same targets were perturbed with identical optical stimulation parameters (e.g., power) in a random order, while the mouse remained head-fixed in the VR rig on the spherical treadmill and the VR screen was dark (Fig. 1E, Fig. S3A).

### Quantifying Causal Connectivity

Because our hypothesis was that the causal connectivity depends on the behavioral state of the animal, we classified trial repeats of the task by behavioral outcome prior to calculating the “in-task” influence. Specifically, we calculated influence separately for ipsilateral and contralateral choices (Fig. 2C; laterality refers to the relationship between choice arm and recorded hemisphere). We also classified trials by whether the choice was correct or not; and we will report correct-outcome findings throughout this paper because error trial sampling was low (Fig. S1D). Finally, we used a latent state model to identify and remove trials within multi-trial periods of task disengagement^21^ (proportion of trials removed within session: median, 8%; range, 0–58%; Fig. S4). In summary, influence was calculated using correct and attentive state trial repeats of the task, separately for the two upcoming choices.

To calculate the causal interaction ΔActivity in-task for a specific target-responder pair of neurons, we averaged the responder fluorescence during the post-stimulation period separately within target-specific stimulation trials (Q1-Stimulation in Fig. 2A,C-F) and within comparison trials (Q1-Comparison), to estimate the stimulation-induced change in the response. This quantity was normalized by the standard deviation over post-stimulation responses, and the result was taken as ΔActivity, a unitless quantity that describes the interaction normalized to the range of observed responses (Fig. 2G, Methods).

Our primary analyses focused on quantifying influence, the average of ΔActivity over many different target-responder pairs. To ensure greater statistical rigor^22^, we employed mixed-effects models to estimate influence throughout this paper (Fig. 2H). The fixed-effects terms in these models (coefficient estimates indicated by *β*) estimate influence or a shift in influence within the population; *β*s have the same scale as the measured influence, and are similarly unitless. The random-effects terms in these models (indicated by u) capture across-individual variability by estimating constant offsets per individual (Methods). We report estimates of the fixed-effects coefficients (*β*) in the main text; model descriptions and statistics can be found in Tables 1, S1.

As shown in Fig. 2I, activations induced in the target neuron ROI generated responses reliably (left panel, *β* = 0.48, \*\*\*\**P* < 0.0001). Fluorescence responses at nearby distances (< ∼100 μm) from the target neuron may be affected by signal from the dendrites of the target neuron^23^, making it difficult to use them as a measure of responder activation. Given the limited spatial spread of AAV expression (Fig. 2J), neuron pairs at separations greater than 250 μm are more likely to include neurons with variable expression. For these reasons, although the magnitude of influence is substantially larger (∼10X) at smaller separation distances, we primarily focused on influence averaged over the 100–250 μm range (referred to as the “network effects” range) in this paper. This range contains a large fraction of measured neuron pairs (86% of pairs within 250 μm were in the 100–250 μm range).

“No-task” influence was estimated using measurements from the no-task stimulation session. In this case, since the animal was not performing a specific task, ΔActivity measures were not calculated by comparing groups of trial repeats with and without stimulation. Instead we compared fluorescence post-stimulation to fluorescence pre-stimulation, normalized to the standard deviation of fluorescence in all post-stimulation time bins (Fig. S3B,C).

### Causal Connectivity is Task-Modulated

Is causal connectivity modulated during task performance? To probe this, we examined differences in connectivity between the no-task and in-task conditions.

The dependence of influence on the separation distance between target and responder has been reported to be excitatory at short distances and suppressive at larger separations^11,9,10,13^. Consistent with these prior studies, we found that no-task influence decreased as the separation between the point of stimulation and the responder increased, crossing over from excitatory to inhibitory effects at ∼150 μm separation (Fig. 3A–C). This is presumably the distance at which suppressive signals from disynaptic inhibitory pathways begin to outweigh the excitatory effects of monosynaptic connections.

**Figure 3.**
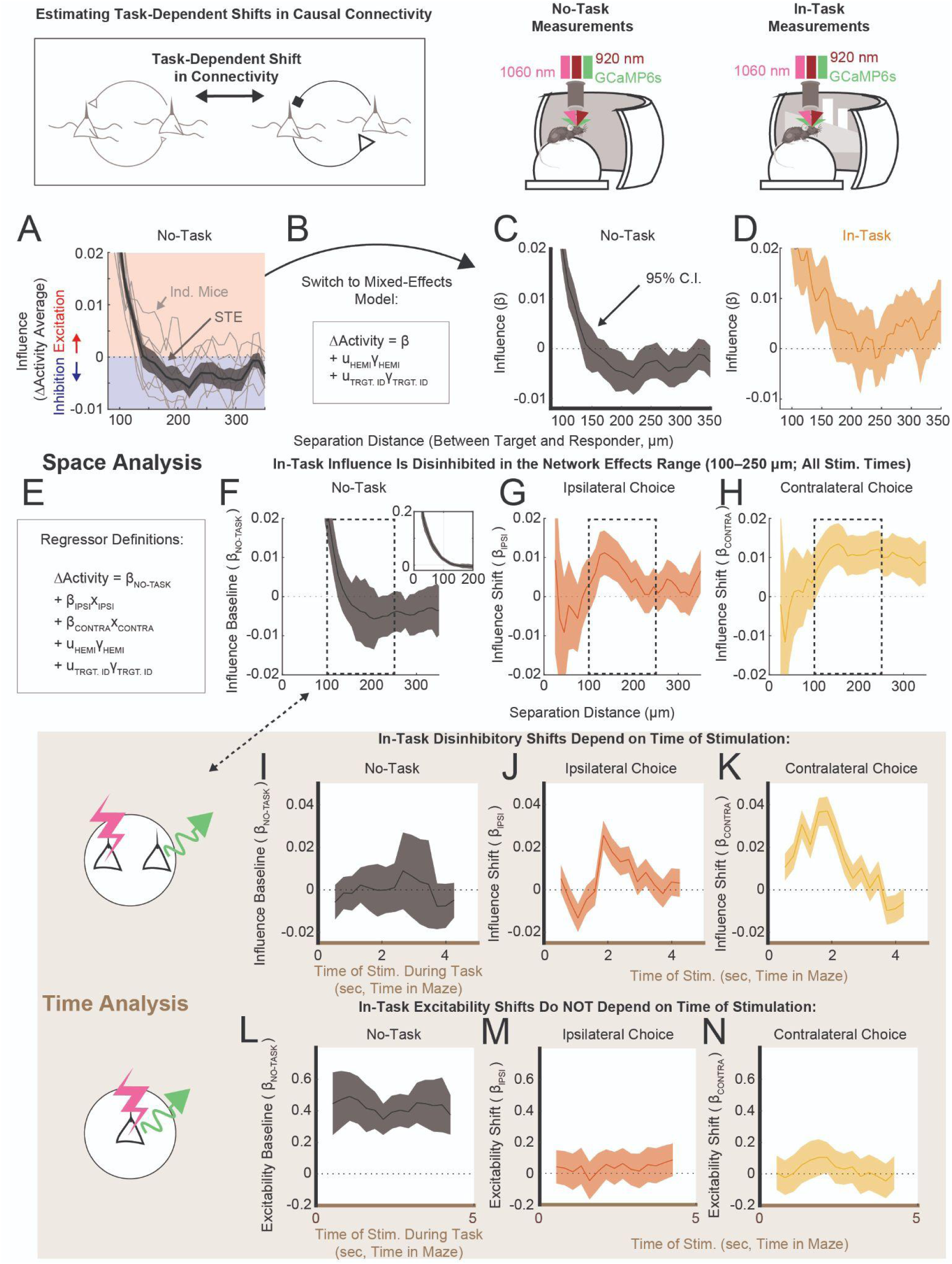
In-Task Causal Connectivity Is Choice- and Time-Dependent. **A.** No-task influence vs. separation distance between point of stimulation (target) and responder (20 μm rolling bin), averaged within mouse (gray traces), and then across mice (black, shaded area = STE; 7 mice, 495 targets, 116 542 target-responder pairs). **B.** Regressor definitions, related to panels C,D. **C.** No-task influence vs. separation distance. Same data and binning as in panel A, (8 hemispheres, see Model 1.1, Table 1). **D.** In-task influence vs. separation distance (same data as in Fig. 2I, note the different y-axis: 20 μm rolling bin; 11 hemispheres, 590 targets, 160 076 target-responder pairs). **E–N**. To compare across experimental conditions directly, target-responder pairs were included for analysis only if they were sufficiently sampled in no-task, ipsilateral and contralateral conditions (Model 1.2, Table 1: 8 hemispheres, 389 targets, 142 034 pairs). **E.** Regressor definitions, related to panels F–N. Here, in-task ΔActivity was treated as a combination of the baseline no-task influence (*β*_NOTASK BASELINE_) and a task-dependent influence shift (*β*_IPSI_, *β*_CONTRA_). Panels F–H use a 50 μm rolling bin. **F.** Influence (baseline, no-task) vs. separation distance. Inset shows the scale of influence for smaller separations. **G,H**. Task-dependent shifts on ipsilateral (**G**) and contralateral (**H**) trials. **I–K.** Influence estimated using target-responder pairs in the network effects range (100–250 μm distance; indicated by dashed line in panels F–H; 8 hemispheres, 389 targets, 53 662 target-responder pairs), and reorganized by the time of target stimulation within the maze (∼1 sec rolling bin). **I**. Influence (baseline, no-task) vs. time of target stimulation in-task. **J,K**. Influence shift vs. time of target stimulation in-task, on ipsilateral (**J**) and contralateral (**K**) trials. **L–N**. An identical analysis to panels I–K, performed on neurons directly excited by stimulation (<25 μm; 236 targets, 359 responders). **L**. Excitability (baseline, no-task) vs. time of target stimulation in-task. **M,N.** Excitability shift vs. time of target stimulation in-task, on ipsilateral (**M**) and contralateral (**N**) trials.

At the same separation distances (∼150 μm), in-task influence was not inhibitory (Fig. 2I, Fig. 3D). To better understand this task-dependent change, we examined target-responder pairs that were measured in both no-task and in-task conditions (Fig. 3E–N). We grouped ΔActivity in pairs with similar separation distances (50 μm bins), and estimated in-task influence as a combination of the no-task influence (*β*_NO-TASK_) and a task-dependent shift (*β*_IPSI_ or *β*_CONTRA_, Fig. 3E; Model 1.2, Table 1). This identified disinhibitory shifts in influence in the ∼100–250 μm range on ipsilateral trials (e.g., at 150 μm, *β*_IPSI_ = +0.010, \*\*\**P* < 0.001, Fig. 3G) and for all separations exceeding ∼100 μm on contralateral trials (at 150 μm, *β*_CONTRA_ = +0.013, \*\*\*\**P* < 0.0001, Fig. 3H). Because task-dependent shifts in causal connectivity were apparent in both choice conditions for target-responder pairs separated by 100–250 μm (the network effects range), we focused on this subpopulation of pairs.

Reorganizing target-responder pairs (in the network effects range) by the time of target stimulation within maze traversal, revealed that the shifts in influence followed time courses that were choice-specific (Fig. 3 J,K). On ipsilateral trials, influence shifts were suppressive early on in maze traversal (Fig. 3J, at 1 sec, *β*_IPSI_ = -0.013, \*\*\**P*_CORR_ < 0.001), becoming excitatory and peaking in excitation ∼2 seconds into maze traversal (Fig. 3J, at 1.8 sec, *β*_IPSI_ = +0.026, \*\*\*\**P*_CORR_ < 0.0001). This peak-excitation time corresponds to the end of the Cue and entry into the Delay sections of the maze (Fig. S5A). On contralateral trials, the shift in influence was excitatory throughout the first half of maze traversal (Fig. 3K: at 1 sec, *β*_CONTRA_ = +0.031, \*\*\*\**P*_CORR_ < 0.0001; at 1.8 sec, *β*_CONTRA_ = +0.037, \*\*\*\**P*_CORR_ < 0.0001). In both choice conditions, shifts in the influence decayed to zero by the time mice turned into one of the arms of the T-maze (∼3–4 seconds, Fig. S5A). We also observed qualitatively similar time-dependent trends in target-responder pairs further away (>250 μm, Fig. S5I–K). We emphasize that all of these modulations were highly statistically significant. In contrast, influence estimated during the no-task experimental condition, in the same target-responder pairs, did not deviate significantly from zero (Fig. 3I, in-task stimulation time of 1 sec, *β*_NO-TASK_ = -0.002, *n.s.*).

We also estimated whether in-task influence statistically deviated from zero by estimating *β* separately within ipsilateral or contralateral conditions (Model 1.3, Table 1). This identified a period of net excitation during contralateral cue presentation (Fig. S5E, at 1.8 sec, *β* = +0.042, \*\*\*\**P*_CORR_ < 0.0001), corresponding to ∼11% more positive than negative signed interactions in the population (Fig. S5Q). The task-dependent change in the distribution of ΔActivity that underlies these excitatory shifts can be found in Fig. S5R-U.

In principle, task-dependent trends in baseline activity could modulate the excitability of target neurons, leading to a modulation of influence without changes in the causal connectivity. We tested whether this was the case by analyzing the excitability of the directly activated neurons (<25 μm from stim). Excitability was not modulated by the ongoing task (Fig. 3M,N). Additionally, there were no time-dependent changes during the task in excitability (two-sample t-test, ipsi excitability, 1 sec vs. 1.8 sec, *n.s.*; contra, 1 sec vs 3.5 sec, *n.s.*). This confirmed that time-dependent changes in in-task influence reflected changes in the connectivity and not in the excitability.

At this point, we had found causal connectivity to be different between the no-task and in-task conditions. Additionally, connectivity changed in a choice-specific, time-dependent manner. The absence of no-task to in-task shifts in connectivity at later stages in the maze suggested that shifts in connectivity were specific to particular computations, an idea that we pursued next.

### Decision-Making Processes Modulate Causal Connectivity

In this task, both navigation and decision-making processes correlated with the time of stimulation, so we next attempted to isolate the contribution of evidence (observed by the time of stimulation) to changes in the causal connectivity (Fig. S6, Methods). Randomized cue presentation times across trials made it possible to identify a small number of targets for which ΔActivity could be measured separately in groups of trials with cumulative evidence above (“more-evidence”) or below (“ambiguous”) a threshold value (Fig. S6A–C). Because the comparison between these two evidence-based conditions was constrained to the same targets, the timing of stimulation in the two evidence-based conditions was statistically matched. Consequently, navigation-related dependencies (e.g., position in maze during stimulation) were matched across comparisons.

We next identified target-responder pairs with significant evidence-modulated ΔActivity (tested against evidence-agnostic bootstrap resamples, Methods), and then probed if the influence estimated over these pairs was systematically modulated by evidence. Using separate models per condition (Model 2.0, Table 1), we found net excitation during contralateral cue presentation, though with low statistical significance, likely due to the small sample sizes in this analysis (2 or more cues vs. 1 or less, network effects, ipsi comparison: Fig. S6D, ambiguous, *β* = -0.013, *n.s.*; Fig. S6E, more evidence, *β* = -0.020, *n.s.*; contra comparison: Fig. S6G, ambiguous, *β* = -0.10, \**P* < 0.05; Fig. S6H, more evidence, *β* = +0.13, \*\**P* < 0.01). Evidence-dependent shifts estimated with comparison models (Model 2.1, Table 1) found disinhibition following 3–4 ipsilateral cues (Fig. S6J,K: evidence threshold of 3.5, *β*_AMB_ = -0.08, \*\*\**P* < 0.001; *β*_EV_ = +0.11, \*\*\*\**P* < 0.0001), or 2 contralateral cues (Fig. S6L,M: evidence threshold of 1.5, *β*_AMB_ = -0.13, \*\*\*\**P* < 0.0001; *β*_EV_ = +0.29, \*\*\*\**P* < 0.0001). Thus increasing evidence had a disinhibitory effect on the causal connectivity. In the case of 1–2 contralateral cues, the causal connectivity was net-excitatory. 1–2 cues may be sufficient for animals to commit to a decision in this no-distractors task. Thus it is possible that these changes in causal connectivity reflect a change in circuit processing following decision commitment.

To probe for a link between decision-making computations and the connectivity explicitly, we categorized each stimulation time on each trial by the ongoing computational epoch, and recalculated influence with this additional constraint (Fig. S7A). We defined six computational epochs. By experimental design, most of the optogenetic stimulation occurred during the Cue and Delay. Because there can be many cues presented on a given trial (cues per trial: median, 8; range, 2–16; standard deviation, 2.6), mice can reach a decision early in the Cue. For this reason, we split the Cue section into half to isolate the epoch underlying decision formation from the subsequent epoch. We refer to these two epochs as “Early Cue” and “Late Cue” (also abbreviated “Cue 1” or “Cue 2”). Indeed, we observed a large increase in the correlation of neural activity (or head direction angle) with choice from the Early Cue to Late Cue, suggesting that the decision is mostly formed by the Late Cue (Fig. S7B,C). For these reasons we link the Early Cue with decision formation, and assume that the time of decision commitment happens in between the Early and Late Cue. We similarly split the Delay section into halves, as the Late Delay contained most of the motor turn indicating choice (Fig. S7D).

How did specific computational epochs modulate the connectivity? To probe this, we used a mixed-effects model to estimate in-task influence as a combination of the no-task influence (*β*_NOTASK_) and an epoch/choice-dependent shift (*β*_EPOCH / CHOICE_, Fig. 4B). This model identified significant modulation of connectivity during the Cue and Early Delay (Fig. 4C–E), but not preceding cue appearance (Pre Cue) or in later epochs (Late Delay, Post Delay). During the Early Cue, the sign of the influence shift was inhibitory during ipsilateral trials and excitatory during contralateral trials (*β*_CUE 1 / IPSI_ = -0.021, \*\*\*\**P*_CORR_ < 0.0001; *β*_CUE 1 / CONTRA_ = +0.018, \*\*\*\**P*_CORR_ < 0.0001). During the Late Cue and Early Delay, influence shifts were disinhibitory for both upcoming decisions (e.g., *β*_CUE 2 / IPSI_ = +0.029, \*\*\*\**P*_CORR_ < 0.0001).

**Figure 4.**
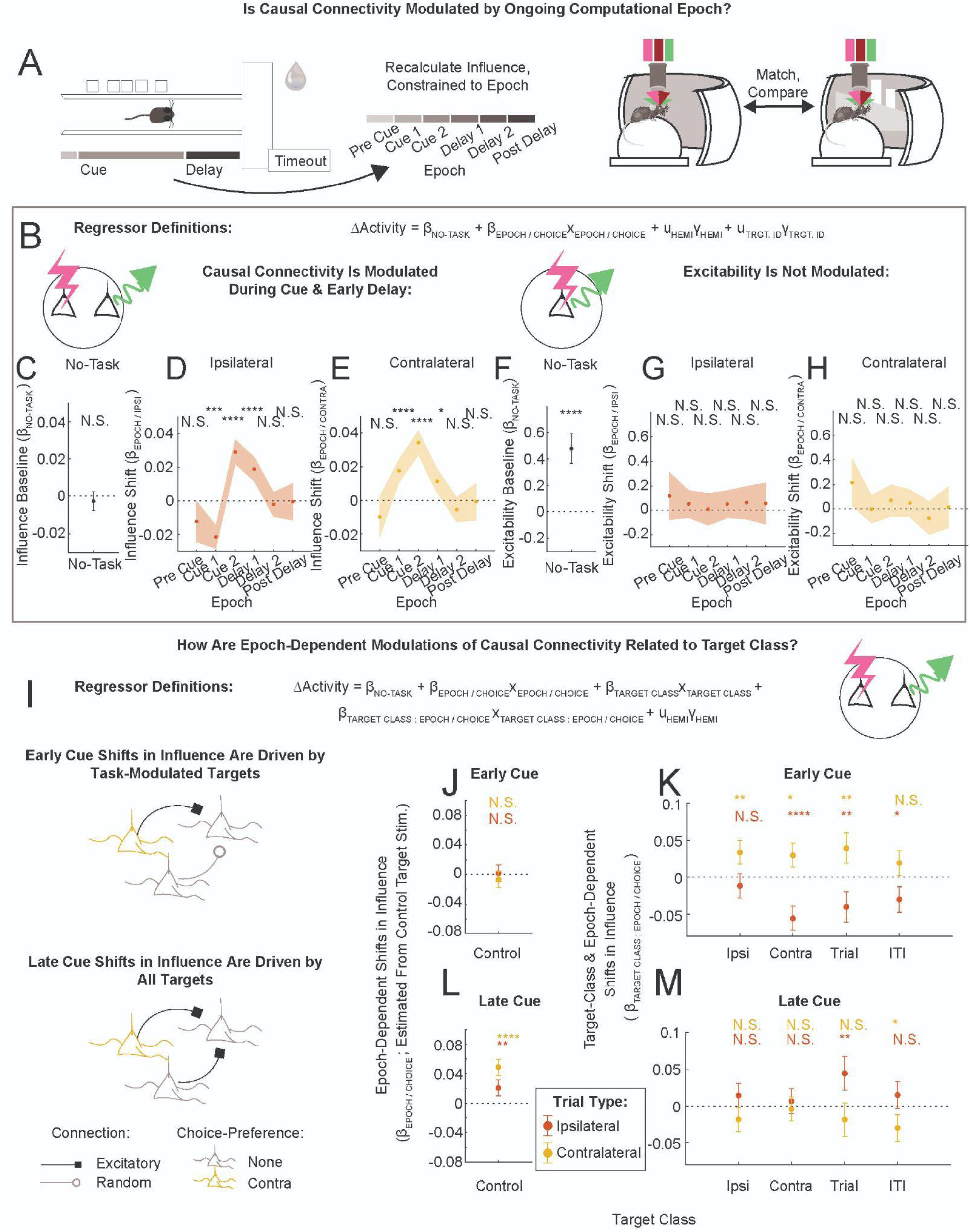
Causal Connectivity Is Modulated by Epoch/Choice and Target Class. **A.** Schematic: In-task influence was recalculated constrained to the ongoing computational epoch, evaluated per-trial per-stimulation time bin. **B.** Definitions of regressors in the epoch/choice-dependent model of influence (panels C–E). In-task influence was estimated as a combination of the no-task baseline coefficient (*β*_NO-TASK_) and an epoch/choice dependent shift (*β*_EPOCH / CHOICE_). Model 3.0 in Table 1: 8 hemispheres, 300 targets, 43 883 pairs. **C–E**. Influence baseline (**C**) and epoch/choice-dependent shifts (**D**,**E**), in the network effects range. Shaded areas indicate uncorrected 95% C.I., asterisks indicate significance values that were corrected for multiple comparisons (Bonferroni, 12X). **F–H**. Excitability baseline (**F**) and epoch/choice-dependent shifts (**G**,**H**), calculated using the directly activated responders (<20 μm, 8 hemispheres, 214 responders). Same model form as in panels C–E. As in Fig. 3L–N, task-dependent shifts in excitability were not significant. **I**. Definitions of regressors in the epoch/choice and target class-dependent model of influence (panels J–M; Model 3.1 in Table 1; full model coefficients can be found in Fig. S7L–Q). In panels J–M, error bars are uncorrected 95% C.I., asterisks indicate 30X corrected significance values. **J,L**. Influence shifts due to ongoing epoch/choice, estimated from stimulation of Control targets, for Early Cue (**K**) and Late Cue (**L**). Ipsilateral choice in red, contralateral choice in gold. **K,M.** Influence shifts due to the combination of target class and epoch/choice, shown for Early Cue (**K**) and Late Cue (**M**).

The excitability did not follow similar trends across epochs (estimated in directly activated neurons, Fig. 4F–H, two sample t-test: ipsi Early Cue vs. Late Cue, *n.s.*; contra Early Cue vs. Late Delay, *n.s.*). We also characterized running speed (Fig. S7E) and global neural activity (Fig. S7F–J) during these computational epochs: there were no clear relationships between these quantities and task-dependent shifts in influence. It is noteworthy that the population-averaged neural activity increases apparently monotonically from a minimum during the Early Cue to peak in the ITI (Fig. S7F-J, see also Discussion). In summary, significant shifts in in-task connectivity occurred selectively during the Cue and Early Delay epochs.

### Task-Modulated Target Neurons Drive Choice-Dependent Shifts in Causal Connectivity During the Early Cue

Do particular subpopulations of excitatory neuron pairs drive task-dependent shifts in influence? Because like-to-like interactions are hypothesized to underlie decision-making circuitry^24^, and have been observed in other tasks and brain areas (e.g., Refs.^25,9,13^) we were interested in models of the influence that could capture a relationship between causal connectivity and similarity in encoding properties between target and responder neurons. To probe this, we defined coding similarity as the cosine distance between target neuron and responder neuron vector representations in the regressor space of the encoding models defined in Fig. S2. We then searched for which features characterized the shift from no-task to in-task influence (Fig. S8). These statistical tests identified ongoing epoch/choice and target class, but not coding similarity, as the relevant features.

Separately, we used the encoding properties of responder neurons to classify responders into the most appropriate target class, and probed whether particular target class : responder class pairings modulated causal connectivity (Fig. S9). This approach also did not find evidence for like-to-like, or opponent inhibition motifs in the causal connectivity.

For these reasons, characterizations of influence in the analyses presented below will refer to the encoding properties of the target neurons, not the responders. We emphasize here that even though ΔActivity was calculated using the fluorescence response of the responder neurons, our most successful categorization of influence was based on the encoding properties of the stimulated target neurons.

We incorporated target class and epoch/choice features into a comprehensive model of in-task influence (Fig. 4I). In this model, the no-task baseline term (*β*_NO-TASK_) was estimated from stimulation of Control targets. All other terms captured shifts in influence due to ongoing epoch/choice (relative to no-task) and / or due to target class stimulated (relative to Control). Full model results can be found in Fig. S7K-Q.

This approach revealed that influence shifts during the Early Cue were driven by task-modulated target neurons (Fig. 4J,K). On ipsilateral trials, Contra, Trial and ITI target classes increased inhibition (Fig. 4K, red, *β*_CONTRA TRGT : CUE1 / IPSI_ = -0.055, \*\*\*\**P*_CORR_ < 0.0001; *β*_TRIAL TRGT : CUE1 / IPSI_ = -0.040, \*\**P*_CORR_ < 0.01; *β*_ITI TRGT : CUE1 / IPSI_ = -0.030, \**P*_CORR_ < 0.05). On contralateral trials, Ipsi, Contra and Trial target classes provided additional excitation (yellow, *β*_IPSI TRGT : CUE1 / CONTRA_ = +0.034, \*\**P*_CORR_ < 0.01; *β*_CONTRA TRGT : CUE1 / CONTRA_ = +0.030, \**P*_CORR_ < 0.05; *β*_TRIAL TRGT : CUE1 / CONTRA_ = +0.040, \*\**P*_CORR_ < 0.01). Notably, Control target stimulation did not yield an effect during the Early Cue (Fig. 4J, *β*_CUE1 / IPSI_ = +0.001, *n.s.*; *β*_CUE1 / CONTRA_= -0.007, *n.s.*). This revealed that the differences in ipsilateral and contralateral influence during the Early Cue were driven specifically by targets with task-modulated coding properties.

In contrast, influence during the Late Cue was disinhibited for all pairs, independently of target encoding properties. Epoch/choice-dependent shifts measured from Control target stimulation were positive relative to the no-task baseline (Fig. 4L, *β*_CUE2 / IPSI_ = +0.021, \*\**P*_CORR_ < 0.01; *β*_CUE2 / CONTRA_ = +0.049, \*\*\*\**P*_CORR_ < 0.0001). Additionally, most target class-dependent offsets did not statistically deviate from zero (Fig. 4M, e.g., *β*_IPSI TRGT : CUE2 / IPSI_ = +0.014, *n.s.*; for ANOVA on term, see Fig. S8D). Thus, in contrast to the Early Cue results, influence during the Late Cue was generally disinhibitory, for all target-responder pairs.

How different is the connectivity underlying ipsilateral and contralateral choices, per epoch? So far we had approached this question indirectly, estimating shifts in in-task influence relative to a no-task baseline. Here, we estimated the choice-dependent change in ΔActivity by subtracting ipsilateral-choice ΔActivity from contralateral-choice ΔActivity, per target-responder pair (Fig. 5A,B). As previously, we tested which features capture variability in the choice-dependent change in influence. We again found a dependence of causal connectivity on target class, not on coding similarity (Model 4.3, Table 1: ANOVA tests in the Early Cue: coding similarity, *F*-statistic = 0.0, *n.s.*; target class, *F*-statistic = 14.0, \*\*\*\**P* < 0.0001). As before, the change in influence during the Early Cue was driven by task-encoding, not Control, targets (Model 4.0, Table 1. Cue 1: Fig. 5D, *β*_CONTROL TRGT_ = -0.002, *n.s.*; Fig. 5E,F, *β*_IPSI_ = +0.052, \*\*\*\**P*_CORR_ < 0.0001, *β*_CONTRA_ = +0.054, \*\*\*\**P*_CORR_ < 0.0001, *β*_TRIAL_ = +0.041, \*\**P*_CORR_ < 0.01, *β*_ITI_ = +0.036, \**P*_CORR_ <0.05). Notably, there was very little target class-dependent modulation in other epochs (Fig. 5E, white is *n.s.*). Analyses of the causal connectivity using different criteria to select the attentive state yielded qualitatively similar results (Fig. S10). In-task influence evaluated separately on ipsilateral and contralateral choice trials can be found in Fig. S11A,B. This approach revealed that choice-dependent modulations of connectivity by target class occurred primarily in the Early Cue, and were largely absent in other epochs.

**Figure 5.**
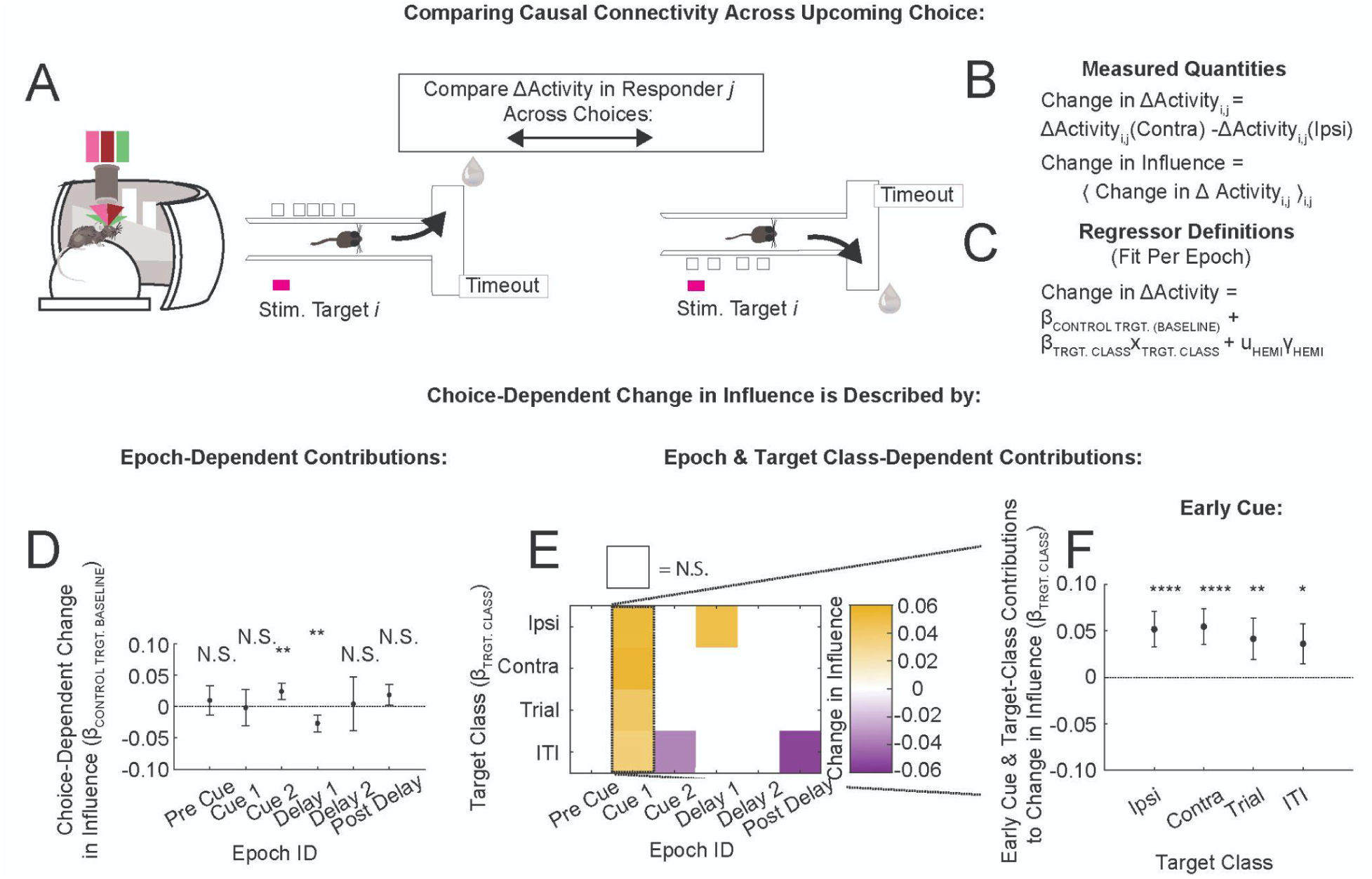
Choice-Dependent Changes in Causal Connectivity Occur Primarily During the Early Cue. **A.** Measurement Schematic. Our goal was to characterize choice-dependent changes in causal connectivity, per epoch (Model 4.0 in Table 1). **B.** Definitions of choice-dependent change in ΔActivity and influence. **C.** Definitions of regressors (models fit per epoch). **D.** Baseline choice-dependent change in influence, estimated per epoch, by Control target stimulation. **E.** Shifts in the choice-dependent change in influence due to target class-specific stimulation, estimated per epoch, shown as a heat map (row: target class; column: epoch). Coefficients which were not significant (*P*_CORR_>0.05) are shown in white. **F.** Target class-dependent coefficients, during the Early Cue (second column in panel E). In all panels, error bars indicate the uncorrected 95% C.I. All significance indicators were multiple comparisons corrected (30x, Bonferroni).

### Conceptual Model

We hypothesized that a combination of (a) target class-specificity in the synaptic connectivity to inhibitory interneurons, and (b) nonlinearities in the interneuron response, could generate choice-dependent shifts in the causal connectivity. We aimed to reproduce the following observations during the Early Cue (summarized in Fig. 6A): 1. The effect of Control target stimulation was not choice-dependent. 2. The effect of perturbation was different when stimulating choice-preferring target or Control target neurons. 3. The effect of choice-preferring target stimulation was choice-dependent (disinhibitory on contralateral and inhibitory on ipsilateral trials). 4. The choice-dependent shift in connectivity did not depend on which choice the targets themselves preferred.

**Figure 6.**
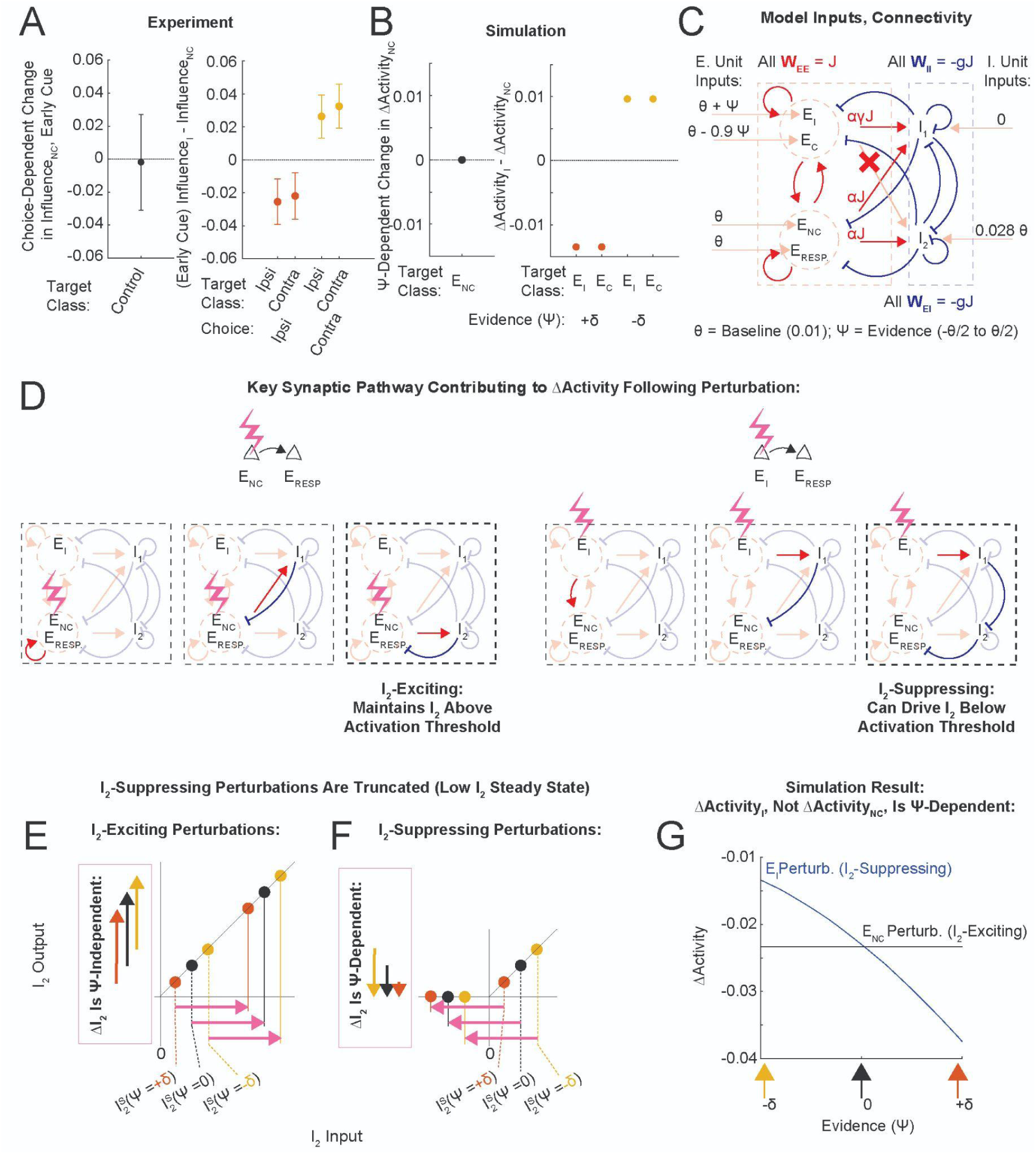
Conceptual Model: Evidence-Dependent Modulation of ΔActivity. We simulated a recurrent network model with four excitatory and two inhibitory units to probe how a nonlinearity in the interneuron units, coupled with asymmetric E to I synaptic connectivity, could account for evidence-dependent shifts in the connectivity. **A.** Experimental properties of causal connectivity (Early Cue, Model 4.2 in Table 1): Left panel: Baseline choice-dependent change in influence, during the Early Cue (coefficients from Fig. 5D, replotted), estimated from Control target stimulation. Right panel: Influence shifts due to target class, during the Early Cue (coefficients from Fig. S11B, replotted; error bars are 95% C.I.). **B.** Simulation results reproducing the experimental measurements in panel A. **C.** Schematic of the network inputs and synaptic connectivity (designed following Ref. ^26^). This network was used to probe how the activity of the responder neuron E_RESP_ changes following E_I_, E_C_ or E_NC_ activity perturbations. Note that differences in ΔActivity between E_I_ and E_NC_ perturbations are due to differences in the E→I synapses (W_IE_). Effects of E_I_ and E_C_ perturbations were identical by construction (Fig. S12C); we follow E_I_ perturbations below. Simulation details can be found in Methods and Fig. S12. **D.** Key synaptic pathways that have large contributions to ΔActivity. This network operated in the underdamped regime, within which the effect of a perturbation decreased with subsequent steps through the pathway (see also Fig. S12B). Notice that because E_I_ did not directly drive I_2_, E_I_-perturbations suppressed I_2_. E_NC_-perturbations excited both interneurons (Fig. S12D). **E-G**. Schematics (E,F) and simulation result (G) illustrating how ΔActivity following E_I_, but not E_NC_, perturbations was *Ѱ*-dependent. **E,F**. I_2_ output vs. input. The steady state of I_2_ (pre-perturbation) is indicated for different values of *Ѱ*. **E.** In the case of I_2_-exciting perturbations, the magnitude of the perturbation-induced change in I_2_ (ΔI_2_, indicated by vertical arrows) did not depend on the steady state of I_2_. **F.** In the case of I_2_-suppressing perturbations, ΔI_2_ was truncated by the low level of steady state activity in I_2_. Because *Ѱ* modulated the steady state of I_2_, and so modulated the contribution of ΔI_2_ to ΔActivity, ΔActivity here was *Ѱ*-dependent. For example, on an ipsilateral evidence trial (*Ѱ*>0), I_1_ received increased excitatory drive, which consequently reduced the steady state activity of I_2_. This reduced the possible contribution of the trisynaptic disinhibitory pathway E_I_ →I_1_ →I_2_ →E_RESP_ (rightmost pathway in panel D) in response to E_I_ perturbations, leading to more inhibitory values in ΔActivity on ipsilateral trials (see also Fig. S12D). **G.** Simulation result: ΔActivity following E_I_, but not E_NC_, perturbations was *Ѱ*-dependent. *Ѱ*-dependent changes in panel B were evaluated at *Ѱ* = +/- δ.

We simulated a recurrent network with four excitatory and two inhibitory units with threshold-linear activation functions (Fig. 6, closely following the methods of Ref. ^26^). We slightly tweaked the simple connectivity within which all excitatory units excite everybody and all inhibitory units inhibit everybody, to have target class-specific E to I connectivity (Fig. 6C). Specifically, ‘choice-preferring’ (E_I_ and E_C_) excitatory units did not connect directly to one of the inhibitory units (I_2_), whereas ‘non-coding’ (E_NC_) excitatory units did. Additionally, E_I_ and E_C_ had the same connectivity, which trivially satisfied observation (4); we will describe the effect of perturbing E_I_. Furthermore, evidence (*Ѱ*) modulated the relative steady state activity levels of the two interneurons. As an example, on ipsilateral evidence trials (*Ѱ*>0), E_I_ received a positive input slightly larger in magnitude than the negative input that E_C_ received. This provided a net increase in excitatory drive to I_1_, and reduced the steady state activity of I_2_. We simulated this network with different levels of evidence (*Ѱ*), and estimated ΔActivity as the stimulation-induced change in E_RESP_ activity (simulation details can be found in the Methods and Fig. S12).

This model reproduced both the evidence (*Ѱ*)-dependent shifts in ΔActivity following E_I_ perturbations and the lack of such shifts following E_NC_ perturbations (Fig. 6A,B,G). This occurred because E_I_ and E_NC_ perturbations had opposite effects on the interneuron element I_2_ (Fig. 6D), which was above but close to threshold. In this regime, low activity levels in I_2_ had no effect on the magnitude of I_2_-exciting perturbations (Fig. 6E), but truncated the magnitude of I_2_-suppressing perturbations (Fig. 6F). Thus E_I_ perturbations, which were I_2_-suppressing, contributed less to ΔActivity when the I_2_ steady state was lower (as would happen in the ipsilateral evidence case, *Ѱ*>0). Because this pathway contributed a disinhibitory term to ΔActivity, ΔActivity following E_I_ perturbations was more inhibitory on ipsilateral evidence trials (Fig. 6G). Notice that because E_NC_ perturbations excited both I_1_ and I_2_, ΔActivity following these perturbations remained constant as *Ѱ* varied. In summary, we reproduced choice- and target class-dependent changes in the connectivity by combining target class-specific E→I synaptic connectivity with the nonlinear response of an interneuron, the operating point of which was evidence-modulated.

## DISCUSSION

We combined all-optical single-cell perturbations with animal engagement in a decision-making task to find that the causal connectivity is selectively modulated by ongoing computation. The ongoing computational epoch was the most significant feature for characterizing causal connectivity, followed by the coding class of the stimulated target neurons; responder coding properties were irrelevant. This unexpected hierarchy of relevant features shows that the causal connectivity is first and foremost dynamic; and suggests that causal connectivity is tailored to fulfill the immediate computational requirements of the circuit.

Task-dependent shifts in influence were most robust during Cue presentation, and early in the Delay, fading away by the time subjects implemented their decision (Fig. 3J,K, Fig. 4D,E). In the same neurons, we observed that activity was minimized at the start of maze traversal and increased monotonically to peak in the inter-trial-interval (Fig. S2I, Fig. S7F–J). If changes in causal connectivity were simply due to increased activity levels driving more circuitry, one would expect task-dependent shifts in connectivity to increase as animals progress through the maze, which was not the case. Relatedly, the global brain state ought to increasingly diverge between ipsilateral and contralateral choice trials as the animals progress through the maze, reflecting the increasing differences in visual flow field and motor pattern output. If choice-dependent changes in connectivity were simply due to divergent global brain states, we would expect to see increasingly large choice-dependent changes in connectivity at later stages of the maze. Yet this was also not the case (Fig. 5). Instead, the largest changes in connectivity occurred when activity levels were low, suggesting a possible connection between reduced excitatory drive of (unobserved) interneuron circuit elements and changes in the connectivity.

As illustrated in our conceptual model, choice-dependent changes in causal connectivity during the Early Cue could be driven by an interneuron circuit element whose operating point is positioned near threshold by ongoing computation. This is a familiar concept from the field of neural networks – in which neural networks change their functional properties in response to different computational demands by moving populations of units above or below threshold^27,28^. Experimentally, it is plausible that somatostatin-expressing (SST) interneurons, which were shown to densely and non-specifically innervate pyramidal cells^29^, and have also been recently implicated in a central role responding to ensemble pyramidal neuron stimulation^30^, underlie the observed shifts in causal connectivity. Modulation of the operating point of SST neurons could be implemented through dedicated local interneuron circuitry (e.g., VIP→SST modulation^31^), or by external inputs from other brain areas, a mechanism suggested by two studies of visual sensory responses across locomotory states^32,33^. Additionally, these studies found that the operating regime of SST interneurons is sensitive to locomotory state or levels of ambient luminance. It is possible that an appropriate combination of behavioral and neural circuit conditions (e.g., running speed, ambient luminance due to cue onset, excitatory population activity levels) sets the operating point of interneuron circuitry during the Early Cue, as suggested by our conceptual model. One way to test this would be to combine manipulation of SST interneuron activity with all-optical measurements of in-task connectivity; however such experiments would require further technical development and are beyond the scope of this current work.

Our measurements did not provide evidence for connectivity motifs such as preferential like-to-like^24,34^ or opponent inhibition^35^ (Fig. S8, Fig. S9). Direct monosynaptic interactions that are expected to contribute to like-to-like connectivity are known to be strong but sparse^25,36^; a property that may be difficult to detect with the population averaging approach that we employed. Perhaps a different experimental modality, such as pairing cellular-resolution optogenetic excitation with electrophysiological recordings and a dedicated sensing algorithm^37,38^, would be better suited to uncovering sparse and strong connectivity motifs. In contrast, our technique is better suited for detecting small population-wide shifts in connectivity, putatively mediated by interneuron circuitry.

Technically, we used a “bread-and-butter” systems neuroscience approach of comparing measurements across behavioral conditions, to identify changes in the average connectivity. Despite the far higher measurement accuracy of ΔActivity in the no-task condition (approaching ∼10X higher effective sampling than in the in-task condition, Fig. S5R), independent analysis of the no-task influence did not identify any notable properties (e.g., Fig. S3I). In contrast, consider the change in detection resolution obtained by incorporating no-task measurements of ΔActivity into the analysis of in-task influence (Fig. S11C–G). Combining the two behavioral conditions reduced uncertainty of task-dependent shifts 3.3x, compared to the 1.2x gains obtained from nearly doubling the size of the dataset (Fig. S11G). Thus our results hinged on contrasting measurements across behavioral conditions. We hope others will find this approach useful when designing all-optical assays of causal connectivity.

These results show that causal connectivity can change quickly with respect to underlying computation, and indicate that care must be taken in extrapolating connectivity across experimental conditions, and even across phases within behavioral trials.

Our observations suggest that decision-making may involve choice-specific alterations in signal propagation through the circuit during decision formation. Signals from task-encoding neurons spread in a choice-dependent manner during decision formation, but not following decision commitment (Fig. 4I–M, Fig. 5). Such dynamic shifts in the connectivity pre- and post-decision could underlie brain-wide, rapid transitions in dynamical regimes recently observed during decision-making^39,40^.

### Limitations of the Study

This study has several limitations. As already mentioned, the statistical significance of an estimate of ΔActivity for a typical pair is low for experimentally-practical sample sizes, which necessitates the use of statistical averages over the population (influence). Second, our statistical averaging approach favors the detection of effects that are common across the population of interactions. Strong effects in a small group of target-responder pairs can be washed out by these population-averaging analyses. Third, we didn’t have sufficient error trial sampling to measure the causal connectivity on incorrect trials. Thus our analyses could not disambiguate cleanly between cue- and choice-dependent shifts in the causal connectivity. Fourth, we analyzed imaging frames that were recorded simultaneously with stimulation, which provided additional sources of noise to the estimates of causal connectivity. In hindsight, a better approach would have been to image and stimulate on alternate frames. Still, analyses excluding imaging frames with stimulation yielded results consistent with those reported throughout this paper (Fig. S13).

## METHODS

### Surgeries

All procedures performed in this study were approved by the Institutional Animal Care and Use Committee at Princeton University and were performed in accordance with the Guide for the Care and Use of Laboratory Animals (National Research Council, 2011).

14 C57BL/6J mice of either gender (Jackson Laboratory), aged 8–20 weeks at the time of initial surgery, were used in this study. A sterile stereotaxic surgery was performed to inject a vector for expression, implant a cranial window and attach a titanium headplate, as previously described^41,42^. Briefly, mice were anesthetized with Isoflurane (Induction: 2–2.5%; Maintenance: 0.5–2%) and given a preoperative intraperitoneal dose of Meloxicam (following changes to our approved protocol, pre-2021: 1 mg/kg; 2021 and later: 5–10 mg/kg), as well as a postoperative subcutaneous dose 24 hours later. During the surgery, mouse body temperature was maintained with a heating pad and rectal probe (Harvard Apparatus). Following hair removal and skin sterilization (Beta-Iodine), the skull was exposed and leveled. A 5 mm (or in a few cases, 3 mm) diameter craniotomy was performed using a handheld dental drill and carbide burs (FG1/4, Midwest), centered over the midline, -2.0 mm AP.

After the craniotomy, the bicistronic construct AAV 2/8 CamKIIa-GCaMP6s-ChrimsonR-KV2.1-HA-WPRE (7 * 10^12 GC/mol titer, created by the PNI vector core, inspired by the GCaMP6M-ChrMINE construct from Ref. ^5^) was backfilled into beveled (20–30 μm tip, Sutter BV-10) glass pipettes (Borosilicate, 1mm O.D. x 0.5mm ID, Fisher NC9617263). Pipettes were positioned just above the dura and then advanced at a 60 degree angle to the normal to final target positions of ∼500 μm lateral offset relative to the midline and ∼250 μm depth (Sutter MP285). After ∼2 minutes of the pipette resting, ∼250nl of the vector was injected in ∼10 minutes with a custom pressure-controlling injection system ^42^. Following a ∼10 minute wait to let the virus settle, pipettes were retracted over a period of 2–3 minutes. We performed 1–3 injections per animal. In two animals (m87 and m88), we also tested the construct AAVrg-CaMKiia-mCherry (1 injection, ∼400 nl of 10^12–10^13 GC/mol), injected into the opposing hemisphere to label across-hemisphere projecting neurons, but did not observe sufficient expression.

Following injections, a cranial window was implanted and secured to the skull with cyanoacrylate (Vetbond). Cranial windows consisted of a stainless steel cannula (Ziggy’s Tubes and Wires, 0.2’’ OD or 0.12’’ OD) cured with UV optical adhesive (Norland’s Optical Adhesive 81) to a coverslip (3 or 5mm round #1 coverslip). Dental cement (C&B Metabond) was used to cover the cyanoacrylate and to attach a titanium headplate to the skull.

### Behavioral Training

After the mice recovered from surgery, mice were acclimated to a restricted water regimen and training began on the accumulation of evidence task in a virtual reality (VR) environment^14,41^.

The virtual reality was projected (Optoma) onto a single spherical mirror (Thorlabs LA1740-P01-SP) that reflected the environment onto the inside of a hollow styrofoam sphere (18’’ OD, 16’’ ID) that had been cut to provide azimuthal coverage of the central ∼270° of the VR. Mice ran on an air-supported spherical treadmill (8’’ diameter) while head fixed to a custom headplate holder that constrained the angular, AP and ML positions of the imaging field of view relative to the VR environment. Body size variations across mice were handled by adjusting the height of the headplate holder relative to the supporting posts, with washers (0.5mm minimal increment).

All mice were trained on the full ATT maze until performance plateaued^14^. All single-cell stimulation sessions were performed in mice (m72, m73, m74, m75, m77, m87, m88, m89, m90) that were trained exclusively in the VR environment of the microscope. The imaging and behavioral sessions include several additional mice (m51, m65, m66, m68, m69) that were trained in a separate training facility and transferred down to the microscope at late stages in the training. These mice were downgraded to the maze level without distractors (T7) on the first day in the new physical location, and then gradually advanced through subsequent training levels.

There were minor modifications to the training protocol from what was previously described^14^. In the original shaping protocol, mice were automatically promoted through the 11 stages of the towers task, denoted as T1 through T11, when they met performance criteria. We modified this in the following ways. Training stages T1 through T4 have a permanently visible turn guide and vary only in length. Mice were started daily on T1 until they were able to autopromote through to T4 (within one day) and run T4 stably. Following this, the visual guide disappeared on training state T5, and the Delay period was gradually lengthened (from T5 to T7). Since the transition to T5 was difficult for the mice, the original training algorithm downgraded the maze difficulty back to T4 for a handful of trials during sessions when animals underperformed for a stretch of time. However, this can backfire: mice can learn to wait for the easier stretches of trials. For this reason, automatic downgrades were turned off if mouse performance stalled for more than 5–10 days at the T4–T5 transition.

The T7 maze was full-length with the full-length Delay, lacking distractors. From this point maze level difficulty was increased manually, when mice ran with long bouts of good performance (typically > 200 consecutive trials before a downgrade to an easier block). Additionally, at T7, the light shield set up was introduced to adapt mice to the typical experimental procedure. The light shield, a black piece of rubber glued to a washer (McMaster 91877A119), was attached to the headplate with a fast setting adhesive (Fast Acting Body Double, with black silicone pigment mixed in), to block VR light from reaching the objective during imaging. At T10 or T11 we would start imaging on occasional sessions, and if both behavioral performance and the quality of the imaged field of view were promising, then we proceeded to the primary stimulation experiments. The experiments reported here were conducted after or instead of the primary stimulation experiments, in the no-distractors T7 maze, when animal performance had plateaued (session numbers, range: 23–39; days after start of training, range: 30–57).

### Histology

Mice were anesthetized with Isofluorane (5%) and injected intraperitoneally with Ketamine (200 mg/kg) and Xylazine (20 mg/kg), before they were transcardially perfused with phosphate buffered saline (PBS), followed by 4% paraformaldehyde (PFA) in PBS. Extracted brains were maintained in 4% PFA overnight at room temperature with agitation, and washed in PBS (3 10-minute rinses) the following day. Coronal sections (50 μm) were obtained on a vibratome (Leica VT1200S). For immunohistochemistry, slices were kept for one hour in a blocking solution of 5% normal goat serum and 0.3% Triton X-100 in PBS. Slices were then exposed to primary antibodies to tag GFP and/or HA (Rabbit Anti-GFP A11122, Mouse Anti-HA PI26183 ThermoFisher) at 500:1 dilutions in blocking solution, in 4° C overnight with agitation. Following primary antibody incubation, slices were washed (3 10-minute PBS rinses) and exposed to secondary antibodies (Donkey Anti-Mouse Alexa 647 A31571 and Donkey Anti-Rabbit Alexa 488 A21206 ThermoFisher), at 500:1 concentrations in blocking solution, in 4° C overnight with agitation. Slices were then rinsed in PBS (2 10-minute washes) and mounted on slides (Fisher 1255015) with DAPI mounting solution (DAPI Fluoromount-G, Southern Biotech). Slices were imaged with a confocal microscope (Leica TCS SP8 X) and processed in ImageJ and MatLab.

### Microscope Overview

Our microscope performed simultaneous two-photon imaging of a calcium indicator and two-photon stimulation of an opsin^1^. Conceptually our custom-built system is similar to the “hybrid holographic’’ approach of Ref. ^4^, where a spatial light modulator (SLM) generates a point cloud of foci in the sample that are co-scanned in a spiral pattern over the scale of a cell soma. We opted for a tighter focus on the stimulation path (axial FWHM of 6-7 μm at center) and introduced an additional (MEMS) scanner into the stimulation path, dedicated to generating the spiral pattern at high speeds (3 kHz), to compensate for the reduced excitation volume of the tighter focus.

Both imaging and stimulation paths were volumetric: the imaging side used a remote-focusing configuration with a secondary objective focusing light onto a mirror mounted on a voice coil, a method that provides high quality optical performance over a wide axial range^43,44^; whereas volumetric positioning of cellular-resolution stimulation was performed with phase mask manipulations by the SLM^45^.

Here we report a brief description of the optical parameters relevant to single-cell perturbations.

### Imaging

On the imaging side, the light source was a 80 Mhz repetition rate Ti:Sapphire laser, Coherent Chameleon Vision II, set to 916 nm. This path utilized remote focusing with a paired secondary objective as in Ref.^43^, generating consistent high quality optical performance across different axial offsets. We measured distortions ∼8% in area between two focal planes 100 μm apart, which significantly outperformed the ∼50% area distortions reported with an ETL over a comparable range^8,46^. PSFs were consistent (FWHM within 50% of central FWHM) over a ∼600 μm axial range.

Imaging frames were collected at a 30 hz frame rate at 5 different axial depths 25 μm apart, ∼680 μm a side, at an effective volumetric rate of 6 hz. Depths of planes ranged from 120 to 350 μm below the coverslip. Imaging power in the top and bottom planes was adjusted manually so that the planes appeared similar in brightness, and the imaging powers for intermediate planes were estimated with linear interpolation. Typical imaging powers were 10–15 mW for the most superficial plane and 15–25 mW for the deepest. ScanImage output was processed with Suite2p^20^, all further analyses were done in MATLAB.

### Stimulation

The stimulation light source (Light Conversion) consisted of a pump (Carbide 40 W, 1030 nm) feeding into a OPA (Orpheus HP, 1.8 W out at 1060 nm, 1 Mhz repetition rate). Of the 1.8 W OPA output, we typically had ∼250 mW out of the objective to work with. At 1 Mhz repetition rate, and with our expression strategy, this is typically sufficient to excite ∼100 neurons at 1 mW target power (with typical power reduction due to SLM efficiency losses of ∼2.5x). Power modulation was controlled with a Pockels Cell (Conoptics), so to optimize power control for single neuron experiments, we introduced a neutral density filter (12.5x reduction) into the stimulation path, before the scanning relays, on the day of the experiment. This still left plenty of power for stimulation with 20 mW available out of the objective. In our experiments we delivered 2–3.5 mW to a target neuron (1 Mhz repetition rate). Preliminary experiments determined the max pre-damage threshold as ∼4 nJ (4 mW at 1 Mhz) at a 100 μm depth below the coverslip.

This beam was scanned with a MEMS scanner (Mirrorcle) at 3 kHz, generating an annular scan over a ∼17 μm outer diameter shape to optimize delivery of excitation with an axially-confined beam (6–7 μm axial FWHM). The spatial light modulator (Meadowlark 1920) deflected the beam towards a single neuron in X, Y, and Z.

### Registration

Registration of the imaging to stimulation paths closely followed the approach of Ref. ^4^: we bleached patterns in a fluorescent slide with the stimulation path and imaged them with the imaging path, to estimate a mapping between the two volumetric systems. On the initial registration we estimated affine transformations per plane for the 5 planes typically used experimentally. These transformations map between SLM-based deflections on the stimulation path and the imaging path. In principle, points at axial offsets in between these planes can be accessed by interpolating the affine transformation parameters (following Ref. ^4^), but in practice we simply calibrated and recorded in the same five planes across all recordings.

After an initial registration, we checked the location of bleach spots at the start of each experimental day, and implemented ∼1–2 μm lateral or axial volume-wide translation adjustments to the registration map. Corrections that would require per-plane recalibrations were required rarely; a single initial registration usually lasted through an experimental run of 4–6 months. These initial checks also included a check of the power out of the objective, a check of the registration, and sometimes a check of the beam profile at the MEMS (Gentech / Dataray). Because the MEMS is the limiting aperture in the stimulation optical path, and the start of the scanning relays, it is a good location to check for possible drift.

### Synchronization

At high stimulation powers, 1060nm excitation will excite GCaMP6s, leading to an artifact in the imaging data, which can potentially saturate the PMT responses and damage them. In our system, gated PMTs (H11706P-41) were included to blank out the imaging signal during high power stimulation. The blanking signal (a TTL pulse going high 1 ms before Pockels Cell modulation) was generated by ScanImage, accessible as the ‘Beam active output’ in scanimage.components.Photostim. Since this blanking signal was recorded, it was a natural choice for post-experimental synchronization of stimulation times to imaging acquisition. In these low-power single-cell stimulation experiments, we did not blank out the acquired signal on the green channel, but recorded the blanking signal through the red PMT acquisition channel for the same synchronization purposes.

### ROI selection

Regions of interest returned by Suite2p^20^ were selected using several criteria on the anatomical shape and statistics of fluorescence. After experiments were completed, we settled on a more stringent set of criteria to analyze the results of perturbation. For this reason we report both thresholds when they differ. On the target identification day sessions, 924 ROIs (median; range, 636–1225) passed criteria out of a total 3093 ROIs (median; range, 2605–3533) returned by the Suite2p algorithm. On the stimulation day, 329 ROIs (median; range, 159–549) passed criteria out of a total 2645 ROIs (median; range, 2087–2985) returned by the Suite2p algorithm. We used two criteria calculated within Suite2p. First, the anatomical “compact” score which measures deviations in the shape of the ROI from a disk had to be lower than threshold (<1.03 on target identification sessions, <1.05 for perturbation sessions). Second, the statistical property, “skew” (third-order moment of the time-collapsed distribution of fluorescence), had to exceed 1 (not used on target identification sessions, but was used on perturbation sessions). Additionally, rois with aberrantly sharp autocorrelation functions in time, and those whose median and modal values of fluorescence were far apart (a check for prolonged elevated baseline activity) were also removed. The later criterion was evaluated over a histogrammed smoothed fluorescence trace (one second moving mean, normalized to unity). If the difference between median and modal (location of peak in histogram) exceeded the ad hoc threshold of 0.08, the ROI was excluded. Analyses were based on the raw fluorescence signal, as neuropil and raw fluorescence were not correlated in the neurons we examined. The raw fluorescence signal was corrected for slow fluctuations, estimated as the moving max over the moving min of a 10 minute rolling window. In a subset of analyses we used a deconvolved fluorescence signal estimated by the OASIS algorithm^47^.

### Selection of Targets for Stimulation

To select targets for stimulation, we characterized the choice- and time-preferences of individual ROIs. We used several ad-hoc criteria to identify and remove periods of inattentive behavior on the target identification day sessions (Fig. 1B,C). Note that this is different to the analyses employed with the stimulation day data, for which inattentive behavior was identified and removed using a latent state model (GLM-HMM, described below, and relevant for Figs. 3–5). Prior to identifying targets for stimulation, trials of excess duration (threshold length of 600 or 2X median number of frames to enter the arm) were automatically removed, and periods of behavioral bias were manually identified (viewed as a 40-trial rolling average performance per trial type) and excluded. Neural activity within trials passing behavioral criteria was processed to identify target neurons.

We selected target neurons for stimulation based on the following six categories (9 targets each for 54 targets total): two choice-preferring (“Ipsi”, “Contra”), two time-preferring (“Trial”, “ITI”), “Active”, and “Control” classes. To select choice-preferring neurons, neurons were sorted by the z-scored difference of fluorescence across correct-went-right and correct-went-left trials, estimated over the [0.5 to 3.5] sec interval following the start of the trial. The top nine neurons within the session were recruited into the right- and left-choice-preferring target classes (Right, Left). These were then converted into Ipsi and Contra labels, relative to the recorded hemisphere (5 out of 20 sessions were recorded in the left hemisphere; of 3 out of 9 mice). Time-preferring targets were selected by their z-scored fluorescence, averaged in the 1st and 3rd second within the maze (collapsed across choice trial types). Trial-preferring neurons were selected if the z-scored activity exceeded 3 in the 3rd second and was below 1 in the 1st second, and ITI-preferring neurons were selected following the opposite criteria. Because GCaMP6s is slowly decaying, the peak of activity for neurons following this selection criteria precedes the trial start by several seconds. The fifth group, Active, contained neurons with elevated activity throughout the maze. Initially these were included to provide a control class of targets that is generally but non-specifically active during the maze. However, this category of neurons was less reliably activated on the day of stimulation than the other categories, so it was removed from analysis throughout this paper (no major qualitative changes occurred as a consequence of this removal). Finally, the Control target class was randomly selected from all remaining neurons, to provide a non-coding-specific target class comparison. When fewer than nine neurons passed criteria for a particular coding target class in a session, remaining targets (to fill up 54 target slots) were recruited into the Control target class for stimulation.

### In-Task Stimulation

Prior to the start of the experiment, trials were randomly selected to be stimulation trials with an 80% probability, starting ten trials into the session. The remaining 20% of trials were withheld to estimate properties unrelated to the connectivity (for example, choice preference). With this sampling rate of stimulation trials, the stimulation design described in the main text (Fig. 2) provided ∼60 repeat stimulations of a target at a fixed time point in a 300-trial session. Subselecting these trials further by decision and correct outcome leaves ∼20–25 stimulation trials per target per correct trial type (and a similar number of comparison trials).

On stimulation trials, the computer controlling the virtual reality environment sent out a trigger signal immediately after the animal entered the Cue section of the VR maze. Note this is the region within which cues could appear, but when and where exactly the cues did start to appear was probabilistically determined on each trial. Following this trigger, a sequence of 144 (8 X 18) stimulation frames was executed, triggered on the start of each imaging frame (at ∼30 hz). On those stimulation bins where targets had been selected for stimulation, 8 consecutive stimulation frames delivered power for 25 ms to the selected target. Each of these frames consisted of three parts (a ScanImage group): (i) a 2 ms pause to allow the stimulation galvos to travel to the center of the field of view from the out-of-optical-path parked location, (ii) a 25 ms exposure with the pockels cell going high, and (iii) another 2 ms pause at the end. Thus for light to go through to the sample, both the stimulation galvos and the pockels cell had to be in the correct configuration. The SLM phase mask (which controlled beam steering in these experiments) would flip to the next (buffered) phase mask at the end of each group. We reasoned the 6.3 ms gap between the end of a stimulation group and the star of the next exposure was sufficiently large, because the switching time of this SLM (Meadowlark 1920 SLM, Overdrive) is 3–4 ms. Non-stimulation bins triggered a ‘park’ pattern in ScanImage which did not allow light through to the sample and did not send a blanking signal for synchronization.

### No-Task Stimulation

On a subset of experimental days we followed in-task stimulation sessions with additional single-cell stimulation sessions in the absence of a task. During these sessions, the VR environment was dark, and mice remained head-fixed. We continued imaging the same fields of view with the same parameters, and stimulated target neurons with identical power and duration as in the in-task stimulation protocol. We generated a new pseudo-random stimulation sequence. This sequence was organized so that each of the 54 targets was stimulated once per 36 seconds, to obtain 100 stimulations per target in a one hour recording. Specifically, each 36 second period was organized as 135 8-frame bins. We assigned each target to one of these bins, and randomized this selection for each 36 second period. We organized the randomization this way to reduce the rate of two-in-a-row stimulations of the same target neuron. Notice that the rate of stimulation here is constant, which is different to the ‘pulsed’ rate of stimulation during the task: during the task all the stimulation per trial is packed into the first ∼4.5 seconds, and there is no stimulation during the ITI. On longer time scales the rate of stimulation in-task is comparable to the 54 targets / 36 seconds rate of stimulation in no-task: in-task, 13.5 targets were stimulated on average per trial, and trial durations were typically ∼8–9 seconds.

### Analyses Using Non-Stimulation Imaging Frames

In the analyses reported in this paper, we opted to analyze imaging frames acquired during stimulation. An alternative option would have been to alternate stimulation and imaging by frame, and analyze non-stimulation imaging frames only. We opted not to, because the stimulation was brief (8 X 25 ms), and we prioritized the efficiency of target activation. As a result, most imaging frames on stimulation trials were recorded with simultaneous stimulation. There are two sources of signal contamination in such frames. First, the photostimulation-induced GCaMP6s excitation created additional fluorescence. Second, an electronic noise bled into the green PMT channel acquiring the fluorescence signal, from the red PMT channel used to acquire the synchronization signal. This signal was visible as a darkening of the fluorescence during stimulation. Because of these sources of signal contamination, we always analyzed the response to stimulation by comparing sets of trials (stimulation and comparison trials) within which the statistics of which imaging frames included stimulation were matched, at times following the conclusion of target-specific stimulation (influence was calculated using fluorescence responses 1–3 seconds following stimulation, substantially later than the 266 ms target stimulation period).

We also explicitly checked if analyses based on (only) non-stimulation imaging frames would give qualitatively similar results to those reported throughout the paper (Fig. S13). To do this, we excluded all imaging frames with stimulation from analysis. We then pooled all the usable imaging frames to calculate post-stimulation and comparison responses. Unlike the main text, where we first averaged over the appropriate two-second period and then calculated the responses and normalization coefficient, here the calculations were performed per imaging frame (without averaging first). This changes the scale of the normalization coefficient. We also had to adjust the sufficient sampling criterion, which had been set to 10 trials of a particular type, to 40 imaging frames of a particular type. Otherwise, the analyses performed on non-stimulation imaging frames only were identical to those reported in the main text. The results were consistent with the key panels in Figs. 3–5 that they aimed to reproduce. This consistency was not particularly surprising as there was no clear way for the stimulation signal contamination to systematically bias the measurements of stimulation-induced changes in activity (as long as the changes in activity were evaluated >266 ms following stimulation).

### Behavioral Trial Selection

Aberrant and warm up trials were removed from analysis (median, 6%; range, 4–16% across sessions) in the following cases: (i) if animals turned around before the Delay period (view angle exceeding π radians in the first 200cm), (ii) if animals entered the Delay period late (time of entry exceeding the median time of entry for that session by twice the width of the distribution; the width was defined as the largest of (a) the median absolute deviation or (b) quarter of the median), (iii) if trials were incompletely recorded in both behavior and imaging, and (iv) the first 10 trials of the recorded block (the warm up period within the block). Performance in recorded sessions (Fig. S1D) was reported over the remaining trials.

### Choice- and Time-Preferences of Neurons

To estimate the choice-preferences of individual neurons in Fig. S1E, we first averaged the fluorescence response of each neuron in the first four seconds of maze traversal on each trial. We then estimated the z-scored preference for correct-went-right over correct-went-left trials per neuron. The z-scored choice-preferences per neuron were converted to a normalized, ranked [-1 to +1] representation, within which -1 indicated the most contralateral-choice–preferring neuron in the session, and +1 indicated the most ipsilateral-choice–preferring neuron. This ranked representation maintained the sign of choice-preference (i.e. all contralateral-choice–preferring neurons were mapped to the [-1 0] range; similarly ipsilateral-choice–preferring neurons were mapped to the [0 +1] range).

Time-preferences (reported in Fig. S1F) were evaluated over the 1st and 3rd second of activity on all correct choice trials. The logarithm of the ratio of these activities (1st over 3rd second) was positive for neurons that prefer the inter-trial-interval (because GCaMP6s fluorescence is slowly decaying, the logarithm will be large when activity in the 3rd second is much lower than in the 1st second), and negative for Trial-preferring neurons. As with the choice-preferences, this was converted to a ranked [-1 1] representation per session, preserving the sign of time-preferences per neuron.

### Latent State Model of Task Engagement

Animals may employ behavioral strategies other than the idealized attend-to-both-sides of the maze strategy. We used a latent state model (GLM-HMM^21^) of behavioral performance to identify and remove behavioral periods within which animals followed a lower-performance behavioral strategy (Fig. S4). Attentive state subselection of behavioral trials was used to preprocess the data for almost all in-task influence calculations (Figs. 3–5 and associated supplements), with the exception of the supplemental analysis of evidence-dependent shifts in connectivity (Fig. S6). We also implemented two alternative attentive state definitions and reproduced key analyses of Fig. 5 (described in Fig. S10), to check that our results were not overly sensitive to the method of attentive state definition.

To fit the latent state model, all no-distractor (T7) sessions recorded in mice (∼37K trials, 15 mice) were grouped together for a meta mouse fit. The GLMs estimated the choice on a current trial based on a constant term (the bias), evidence on the right-hand and left-hand sides of the maze (separate regressors; normalized to the 0-1 range), the prior choice on the trials preceding the current one by 1 to 5 trials (-1 for left, +1 for right; 5 separate regressors) as well as the prior reward (-1 for prior left reward, 0 for no reward, +1 for prior right reward). The GLM-HMM model estimates the best fitting latent GLM states as well as the transition matrix between them. Our procedure followed Ref.^21^: data was split for ∼5 fold cross validation with roughly equal sampling per mouse, to estimate the improvement a multi-latent-state GLM-HMM yielded over a baseline single-state GLM model. This was done varying the number of latent GLM states (1 to 5). There was a clear jump in improvement from 1 to 2 states, and incremental performance improvement after that (Fig. S4B). The 3- and 4-state models didn’t identify simply interpretable states, such as right- vs. left-biased states, as we initially hoped (data not shown). For simplicity, we opted to use the 2-state model to identify poor-performance periods in a standardized way across mice (Fig. S4C,D; see also Fig. S10). At this point we fit the 2-state model to the full dataset, and all behavioral trials with a probability greater than 50% of belonging to the higher performance state (state 1) were classified as attentive. For most sessions, the majority of behavioral trials were classified as attentive (Fig. S4E–G; median, 92%; range, 42–100%). Two (out of 22) sessions were classified as primarily inattentive, and for consistency were removed from all analyses in this paper (not shown).

### Statistics

Statistical significance was primarily evaluated in the context of mixed-effects models^22^, which can incorporate random-effects terms to capture variability in measurements across individuals. We used the random-effects coefficients to allow for constant offsets in influence per recorded hemisphere, and in some cases, target identity. The hemisphere identifier was used to group together the sessions that were recorded in the same injection site, as some of the recorded neurons in these sessions may be the same; though we have not observed any qualitative changes in results from switching between mouse identity, hemisphere, and session identity as the random-effects label. Models are indicated within each figure and listed in Table 1. Detailed statistics are reported in Table S1. Model parameters were estimated with Matlab’s **fitglme**; all statistical comparisons are against a zero baseline. In Table S1, we report p-values uncorrected for multiple comparisons (*P*). Throughout this paper, p-values corrected for multiple comparisons (Bonferroni correction) are indicated as *P*_CORR_, and the multiplicative factor is provided nearby or can be found in Table S1. In figures, asterisks indicate significance values following multiple comparisons correction. Error bars and shaded areas on fixed-effects coefficients are the 95% C.I. returned by the model, and were not corrected for multiple comparisons.

### Calculations of ΔActivity

Throughout the text, we calculated ΔActivity for target-responder pairs in an experimental condition provided that they were sufficiently sampled (defined as at least ten stimulation and ten comparison trials). The stimulation response was always evaluated 1–3 seconds following stimulation, which translates to 11 frames at the volumetric imaging rate (per trial). To calculate the causal interaction ΔActivity in-task for a specific target-responder pair of neurons, we first averaged the responder fluorescence in the post-stimulation period on each trial to estimate the per-trial response. This was separately averaged within target-specific stimulation and comparison trials to estimate the stimulation-induced change in the response, and then normalized by the range of responses (*σ*) to estimate ΔActivity. The normalization coefficient *σ* was estimated as the standard deviation of post-stimulation activity over all behavioral trials with stimulation (which included the target-specific stimulation and comparison trials, as well as the trials stimulating the other two targets assigned to the same stimulation bin, and all behavioral outcomes). Thus *σ* was constrained to be the same for ΔActivity calculated on ipsilateral and contralateral choice trials, for a particular pair of neurons.

We constrained most analyses to work with the same target-responder pairs (e.g., in Fig. 3E–N, ΔActivity had to be sufficiently sampled in no-task, ipsilateral and contralateral conditions in order to be included in the analysis of influence). Details are listed in Table 1.

### Single Neuron Encoding Models

We used single neuron generalized linear models (GLMS) to predict the deconvolved fluorescence activity of individual neurons across trials and time bins using behavioral variables (Fig. S2). Task variables such as sensory evidence vary across time bins within a trial; other variables such as prior choice were represented with a constant value per trial (see below, and Fig. S2B). Responder coding was estimated on the day of stimulation using all correct nonstimulation trials in the attentive state defined by the latent state model (Fig. S4). Target coding properties were estimated using the imaging day data, on all correct non-aberrant trials following the warm up period of ten trials. We estimated target neuron models on the imaging day because there were significantly more (∼5X) nonstimulation trials to use for model fitting, target identity was unambiguous, and all targets were represented.

GLMs were estimated using lasso regularization (Matlab’s **lassoglm**), which aims to sparsify representations. We used 4-fold crossvalidation to train on 3 quarters (**lassoglm** with 3-fold crossvalidation) and test on the withheld quarter. Even with perfect matching between training and testing data, and a model that captures the behavioral dependence of a neuron exactly, the across-trial variability of neural responses sets a noise ceiling on the performance of a GLM model. For this reason we estimated the variance explained normalized by the ceiling, and reported the normalized R^2^_CORR_. Specifically, for each unique behavioral state we estimated the average response in the testing data, which would have been the best-performing prediction. We normalized the true model variance R^2^ by the variance explained by this best-performing prediction, to obtain R^2^_CORR_. The R^2^_CORR_ reported per neuron (in Fig. S2D,E) is the median over the four 4-fold estimates of R^2^_CORR_.

Final model parameters per neuron were estimated using all the data available (**lassoglm** with 4-fold crossvalidation). Including the offset, these models allowed for 22 parameters per neuron. Models were kept if the variance explained on withheld trials was greater than zero (4862 out of a total of 6695 neurons). The first four regressors were the cumulative contralateral cues presented up to the time bin, conditioned on the epoch (other time epochs, 0; this time epoch, scaled so that the max across trials = +1). Conditioning sensory evidence by epoch was done to avoid underweighing the sensitivity of neurons that are sensitive to evidence early, when the total number of presented cues is low (see also Ref. ^16^). Similarly, the next four regressors represented the cumulative ipsilateral cues. The next five regressors were constant across the time bins within a trial: the upcoming choice (contralateral, -0.5; ipsilateral, +0.5), and four prior trial regressors (representing the correct / incorrect and ipsilateral / contralateral combinations; didn’t happen, 0; did happen, 1). Finally, we included an indicator (0, 1) of the ongoing time epoch to identify time-preferences in the population of neurons. We included three additional epochs to those defined before: “Pre-ITI” (including 3 seconds before the trial start), “Inter” (a brief technical pause between arm entry and reward delivery, during which the behavioral computer records data), and “Post-ITI”.

To estimate similarity between target classes based on their coding properties, we used a split-half linear discriminant analysis (Matlab’s **fitcdiscr**). Half of the targets were used to estimate a linear classifier, which was used to predict labels in the withheld half of targets. The confusion matrix from one run is shown in (Fig. S2K). To better visualize the coding properties of the separate classes, we used a two-dimensional embedding of the 21-dimensional target coding profiles (Matlab’s **tsne**). This embedding aims to preserve distances between points, and the implementation was agnostic to the target class label. This embedding suggests that a good portion (∼65%) of Control target neurons do not strongly encode the task features we identified, while the rest could be classified as belonging to the encoding categories used throughout our paper (Ipsi, Contra, Trial and ITI; Fig. S2L).

### Analysis of Evidence-Dependent Shifts in Causal Connectivity

The goal here was to probe evidence-dependent shifts in causal connectivity within the same target-responder pairs (Fig. S6). Because a decision can be reached quickly in the no-distractors task, we aimed to separate out trials within which the animals could have reached a decision, from those within which the animals would still be making up their mind, using the evidence available at the time of stimulation. To do this, we defined an evidence threshold to separate behavioral trials by the evidence at time of stimulation, performed analyses on these separate groups of trials, and then repeated analyses while varying the evidence threshold. “Ambiguous” trials were classified as such if the absolute cumulative evidence (including both sides) was less than or equal to the evidence threshold. “More evidence” trials were classified as such if the instantaneous evidence exceeded threshold; ipsi and contra evidence comparison analyses were performed separately. As in other analyses, ΔActivity was included for analysis from target-responder pairs which were sampled at least ten times in stimulation trials, and ten times in comparison trials, per evidence condition (Fig. S6A–C).

We next identified those target-responder pairs with significant evidence-dependent modulation of ΔActivity. This was done by resampling ΔActivity measurements for each target-responder pair. For each resample, we resampled stimulation trials from the true stimulation trials (matching the number of stimulation trials, resampling with replacement), ignoring the evidence on each trial. We similarly resampled comparison trials from the true comparison trials for the target-responder pair, ignoring the true evidence amount on each trial. We generated this agnostic-to-true-evidence resampled ΔActivity 1000X per pair. If the real change in ΔActivity between evidence conditions was greater than, or less than, 95% of the resampled changes in ΔActivity, that pair was classified as evidence-modulated, and the ΔActivity values were included in the subsequent analysis of influence.

We then asked if evidence modulated influence systematically across the population of target-responder pairs. We grouped evidence-modulated target-responder pairs by distance into 100 μm rolling bins, and used a mixed-effects model with 2-level random-effects to estimate the influence per distance per evidence condition. These analyses were repeated for different levels of the evidence threshold (Fig. S6D–M).

### Feature Selection in Mixed-Effects Models of Influence

We used statistical tests to determine which features (epoch/choice, target class identity and coding similarity) categorize influence, in order to choose the appropriate model (for Fig. 4 and Fig. 5). We report this analysis in the no-task to in-task comparison (Fig. S8). A similar approach to the in-task distributions yielded qualitatively similar results (data not shown). We first fit a comprehensive mixed-effects model that incorporated all terms up to the three-way interaction between the epoch/choice, target class, and coding similarity (Fig. S8A). Marginal tests on the fixed-effects terms identified a very large dependence of influence on the ongoing epoch/choice (Fig. S8B). The far higher significance on terms associated with epoch/choice made it difficult to interpret the significance of other terms. For this reason, we subdivided the data by epoch/choice and fit separate mixed-effects models per epoch/choice condition, to identify the features that were most consistently important conditions (Fig. S8C,D).

### Conceptual Model Simulation

We simulated a recurrent network (Fig. 6), closely following the work of Ref.^26^. Dynamics were simulated following the equation 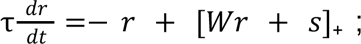 with r the vector of firing rates (including both E and I units), W the synaptic connectivity matrix, s the inputs to each unit, and []_+_ indicating the threshold-linear nonlinearity. We used a 0.1 ms time step, and set τ = 1 ms for both E and I units. Inputs (shown in Fig. 6C) consisted of a constant baseline input per unit, as well as an evidence signal (*Ѱ*, from -Θ/2 to +Θ/2), which provided additional drive to the two choice-preferring E units (+1*Ѱ* to E_I_ and -0.9*Ѱ* to E_C_). The I_2_ baseline input (0.028 Θ) adjusts the relative balance between I_1_ and I_2_, setting the crossover point in ΔActivity_I_ relative to ΔActivity_NC_ in Fig. 6G. In an alternative implementation, we provided equal magnitude *Ѱ* inputs to E_I_ and E_C_, and modulated the activity levels of the interneurons with evidence by providing *Ѱ* as a direct input to I_1_. This approach yielded similar results (data not shown).

The synaptic connectivity matrix was designed to generate small and suppressive ΔActivity, again following Ref.^26^. E to E connectivity was set to J = 1/ N_E_. Inhibitory output weights were set to -gJ (with g = 2). E to I weights for the non-coding neurons (E_NC_ and E_RESP_) were set to αJ (with α = 2.2) to generate small suppressive interactions. Because E_I_ and E_C_ did not connect to I_2_ directly, we compensated for the reduced connectivity by scaling up their drive of I_1_ by an additional factor, γ = 1.9. A consequence of maintaining similar small and negative suppressive effects from excitatory units with different connectivity is that I_1_ receives substantially more drive than I_2_, putting I_2_ near threshold.

To estimate ΔActivity, we ran two simulations of the same network. In one simulation, at the perturbation time point (200 time steps after the start of simulation, Fig. S12B), one excitatory unit received an additional input (amplitude = 0.05). In the other simulation there was no perturbation. ΔActivity was defined as the total difference in the activity of the responder E_RESP_ between the two simulations (starting immediately following the perturbation), normalized to perturbation magnitude^26^.

## ACKNOWLEDGEMENTS

This work was supported by the US National Institutes of Health (NIH) grant 5U19NS132720, and the Simons Collaboration on the Global Brain (D.W.T and C.B.).

We thank Joshua Julian and P. Dylan Rich for guidance on statistical analysis; Scott Bolkan for assistance with GLM-HMM implementation; and Efthymia Diamanti, Jesse Kaminsky, Wynne Stagnaro and Joshua Julian for their thoughtful comments on the manuscript. We thank K. Deisseroth for the GCaMP6m-ChRmine construct that was used in pilot studies that influenced our development of the GCaMP6s-ChrimsonR construct, which was the basis for the work in this paper. We thank Esteban Engel for help with the development of the GCaMP6s-ChrimsonR construct. We thank Juan Lopez and Alvaro Luna for technical support with the VR environment. We are grateful to members of the Tank and Brody labs, as well as the Princeton U19 Brain Grant consortium for helpful feedback throughout this project. For technical instruction, Mark Ioffe thanks Alex Song and Matthias Koch (optical engineering); Steve Lowe (machining); and Yi Gu, Scott Bolkan, Lucas Pinto, Edward Nieh, Alex Riordan and P. Dylan Rich (animal work).

## AUTHOR CONTRIBUTIONS

C.B. and D.W.T. obtained funding. M.L.I., C.B. and D.W.T. conceived the project, M.L.I, S.T. and D.W.T. designed the optical instrumentation, M.L.I. built the optical instrumentation under the supervision of S.T. and D.W.T., M.L.I. designed and performed experiments, analyzed the data, created the visualizations, and wrote the original draft. M.L.I, C.B. and D.W.T. wrote and edited the draft. C.B. and D.W.T. supervised the project.

## Declaration of generative AI and AI-assisted technologies in the manuscript preparation process

During the preparation of this work the author used Gemini in order to improve readability of the article: to suggest alternate phrasings and proofread. After using this tool/service, the author reviewed and edited the content as needed, and takes full responsibility for the content of the published article.

## SUPPLEMENTAL INFORMATION

**Figure S1.**
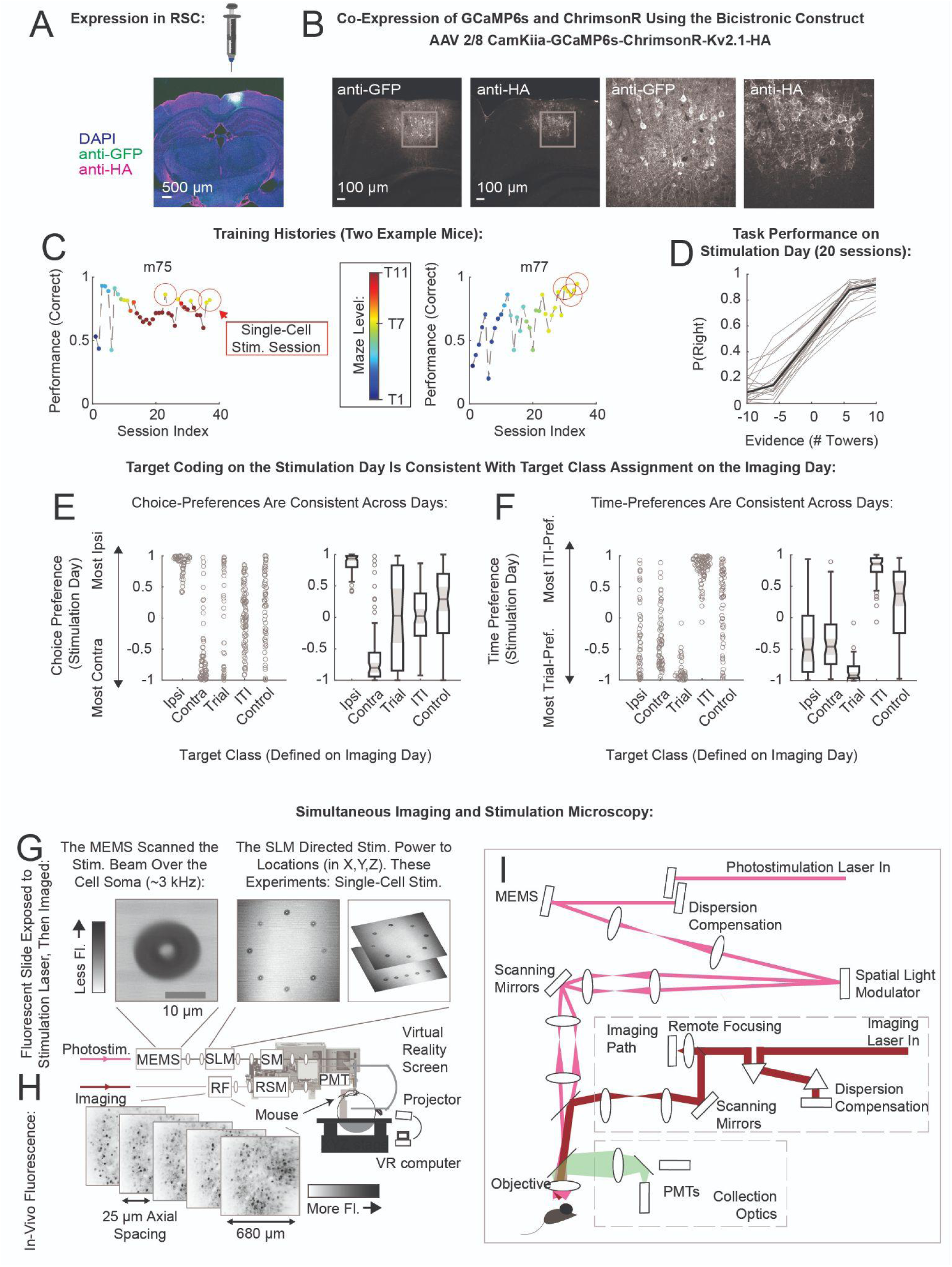
Technical Details, Related to Figure 1. **A,B**. We used a bicistronic construct to co-express the calcium indicator GCaMP6s and the soma-localized opsin ChrimsonR-Kv2.1 using the CamKiiα promoter^1–5^. **A.** Image of viral expression targeting RSC. **B.** Higher magnification of the image in panel A; anti-GFP and anti-HA expression shown separately. Right panels: Expanded view of the boxed region in left panels. Neurons co-express opsin and indicator; ChrimsonR-Kv2.1 expression (anti-HA) appears less prominent in the background. **C.** Performance correct (on all trials per session, including warm up and aberrant trials) vs. session index, for two representative mice. Single-cell stimulation sessions are indicated with red circles. Color indicates maze difficulty. All mice used in the single-cell stimulation experiments were able to perform the no-distractors version (T7) of the ATT well (see panel D). Mice were trained on the full ATT as far in maze difficulty as they could progress (Methods : Behavioral Training). Training histories for all nine mice are shown in Fig. S14. **D.** Performance per session, in trials passing behavioral criteria, but not selected for attentive state (see Fig. S4 for attentive state selection; individual 20 sessions in 9 mice in gray, median in black). Correct performance: median, 88%; range, 72–96%. **E,F**. Choice (**E**) and time (**F**) preferences of neurons close to the stimulation point (directly activated by stimulation), evaluated on nonstimulation trials on the stimulation day (Methods : Choice- and Time-Preferences of Neurons), organized by the target class definition on the imaging day. Left panel: swarm chart; Right panel: statistics (Matlab’s **boxchart**: shaded region indicates 95% C.I. on median; box indicates lower and upper quartiles). For a more detailed analysis of the encoding properties of target classes using generalized linear models, see Fig. S2. **G–I.** Simultaneous imaging and stimulation microscope. **G.** Stimulation path deflections were calibrated by bleaching a fluorescent slide, which was then imaged by the imaging path (top panels). The photostimulation path used a MEMS scanner to create a ∼3kHz spiral scan over a ∼17 μm wide annulus (top left), which was directed towards a target neuron with the spatial light modulator (SLM; top middle and top right). Scanning mirrors (SM) were centered in these experiments. **H.** Averaged fluorescence (mean image from Suite2p^6^) from a recorded session in-vivo (bottom panels, inverted grayscale indicates brighter cells as darker). We recorded from five planes with 25 μm axial spacing in these experiments. The imaging path incorporated a remote focusing (RF) unit consisting of a paired secondary objective and voice coil^7^ to provide high quality optical performance over a range of axial deflections. Resonant scanning mirrors (RSM) and photomultiplier tubes (PMT) completed the imaging path. **I.** Schematic of the optical pathways, described in greater detail in Methods.

**Figure S2.**
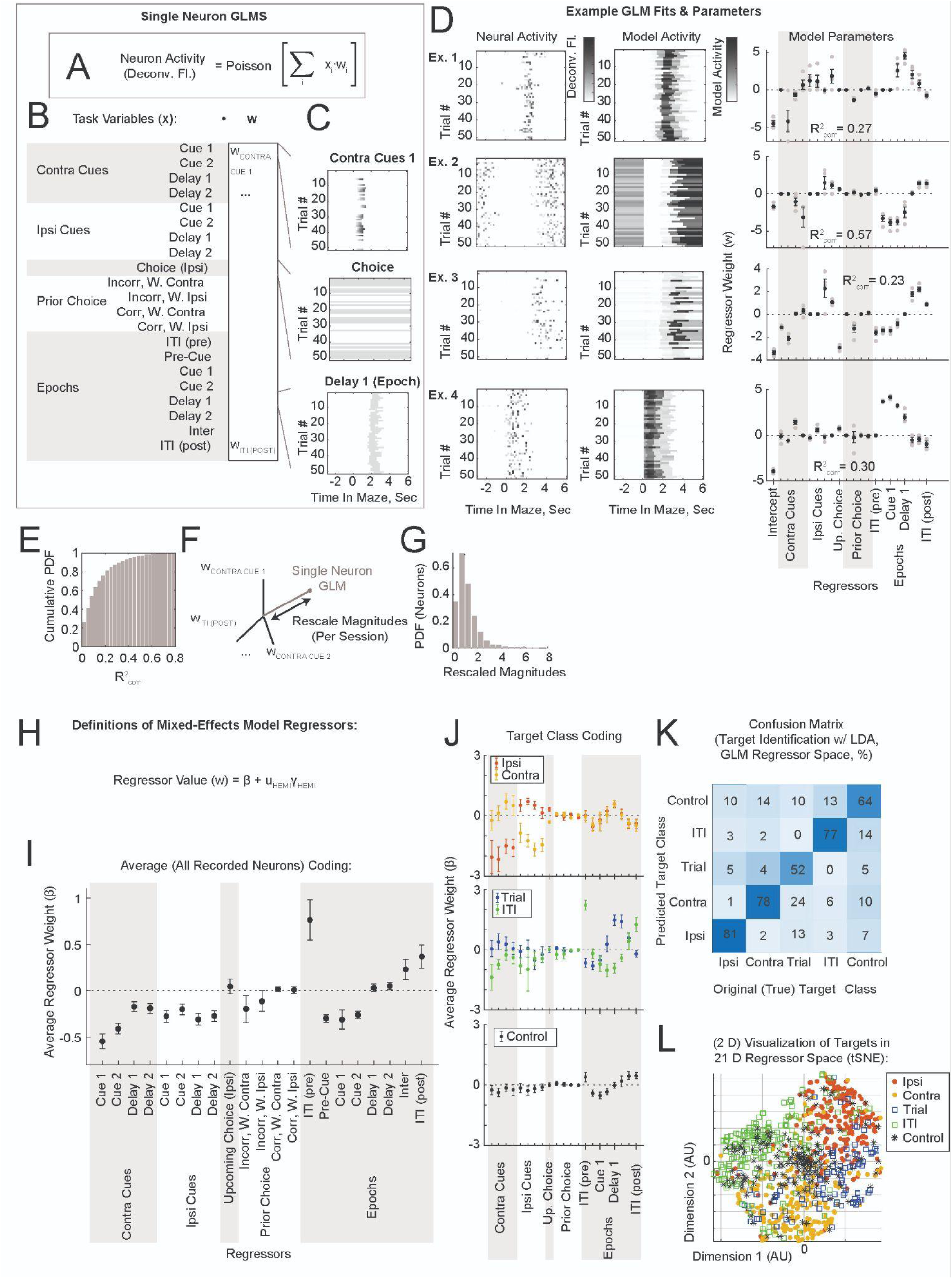
Encoding Models for Single Neurons, Related to Figure 1. We regressed the activity of a single neuron against task variables with generalized linear models (Methods: Single Neuron Encoding Models). **A.** Formula describing the encoding model. Individual GLMs were fit to the activity (deconvolved fluorescence estimated using the OASIS algorithm^8^) of a single neuron on non-stimulation trials. **B.** The task variables used as regressors in the model. **C.** Examples of regression inputs (heat maps of input values, trial (row) by time (column)). All task variables were normalized across trials to a range of 1. Top panel: Sensory evidence input varied within-epoch and across-trials. Sensory evidence was included in the model in 8 terms (top 8 rows in panel B), corresponding to which side the cues were presented on and conditioned to the four maze-traversing epochs (Early Cue, Late Cue, Early Delay, Late Delay). These values were set to 0 in other epochs, and scaled to the 0–1 range within-epoch. Middle panel: Choice on the current trial and the four prior choice conditions (binary values for correct/incorrect and went-ipsi/went-contra) was represented with constant terms per trial. Bottom panel: The ongoing epoch was represented with a binary indicator (other epoch, 0; this epoch, 1). **D.** Four example neurons (rows Ex. 1: Ex. 4). These were randomly selected from all neurons with R^2^ > 0.08. Left column: Heat maps of the deconvolved neural activity. Center column: Predicted Model Activity. Right column: Estimates of the regressor coefficients (w) per neuron. Gray: values from individual 4-fold estimates, black and error bars are mean and standard error across 4-folds. The intercept was not used in subsequent analyses. **E.** The distribution of variance explained across all model fits with sufficient samples and positive variance explained (4862 out of a total of 6695 neurons, 20 sessions). **F.** In order to compare regressors across sessions, we rescaled all regressor vectors so that the median magnitude within-session would be unity. **G.** Distribution of rescaled magnitudes across sessions. **H.** Definitions of regressors (related to panels I,J). The fixed-effects coefficients (□) provide estimates of the average values of the encoding-model regressors w, allowing for variability across hemispheres. **I.** Average encoding of regressors across all neurons (4862 neurons, 11 hemispheres). Black dots and error bars indicate the estimate and 95% C.I. **J.** Average encoding of regressors within target class neurons recruited for stimulation, organized by target class (data from imaging day recording). Top panel: Ipsi (red) and Contra (gold) target classes. Middle panel: Trial (blue) and ITI (green) target classes. Bottom panel: Control target class. The coding of Control neurons seems fairly similar to the average coding profile in panel I. **K,L.** We used the encoding models of each target, and the assigned target class, to assess the separability of target classes by their encoding. **K.** Classifier performance, shown as a confusion matrix, averaged over 1000x 2-fold cross-validated fits. Linear classifiers (Matlab’s **fitcdiscr**) were trained to separate target classes in the regressor space of the encoding models, and tested on a withheld half of targets. The test targets which did not have a classification probability >50% for any target class (16% of all targets) were excluded from this analysis. **L.** Two-dimensional embedding of the 21-dimensional target coding profiles (tSNE, Euclidean distance). This embedding was agnostic to the target class label. The classes with opposing preferences separated well from each other (panel K; Ipsi and Contra; Trial and ITI). Trial targets were frequently misidentified as Ipsi or Contra targets (panel K). Trial targets tend to have choice-preferences (Fig. S1E), and time-preferences of Trial and Ipsi / Contra neurons behave similarly (panel J; Trial targets tend to peak a little later than Ipsi / Contra targets). Control targets were identified as Control in ∼65% of cases (panel K). Most of these are a core non-coding group of neurons (dense core of black asterisks in the center of panel L). The remaining ∼35% fall into the remaining categories, suggesting that the excitatory layer 2/3 RSC population could be categorized as ∼10% Ipsi, ∼10% Contra, ∼5% Trial, ∼15% ITI. For these reasons, throughout this paper we focused on three target class-based comparisons: Ipsi to Contra, Trial to ITI, and all four task-encoding target classes to Control.

**Figure S3.**
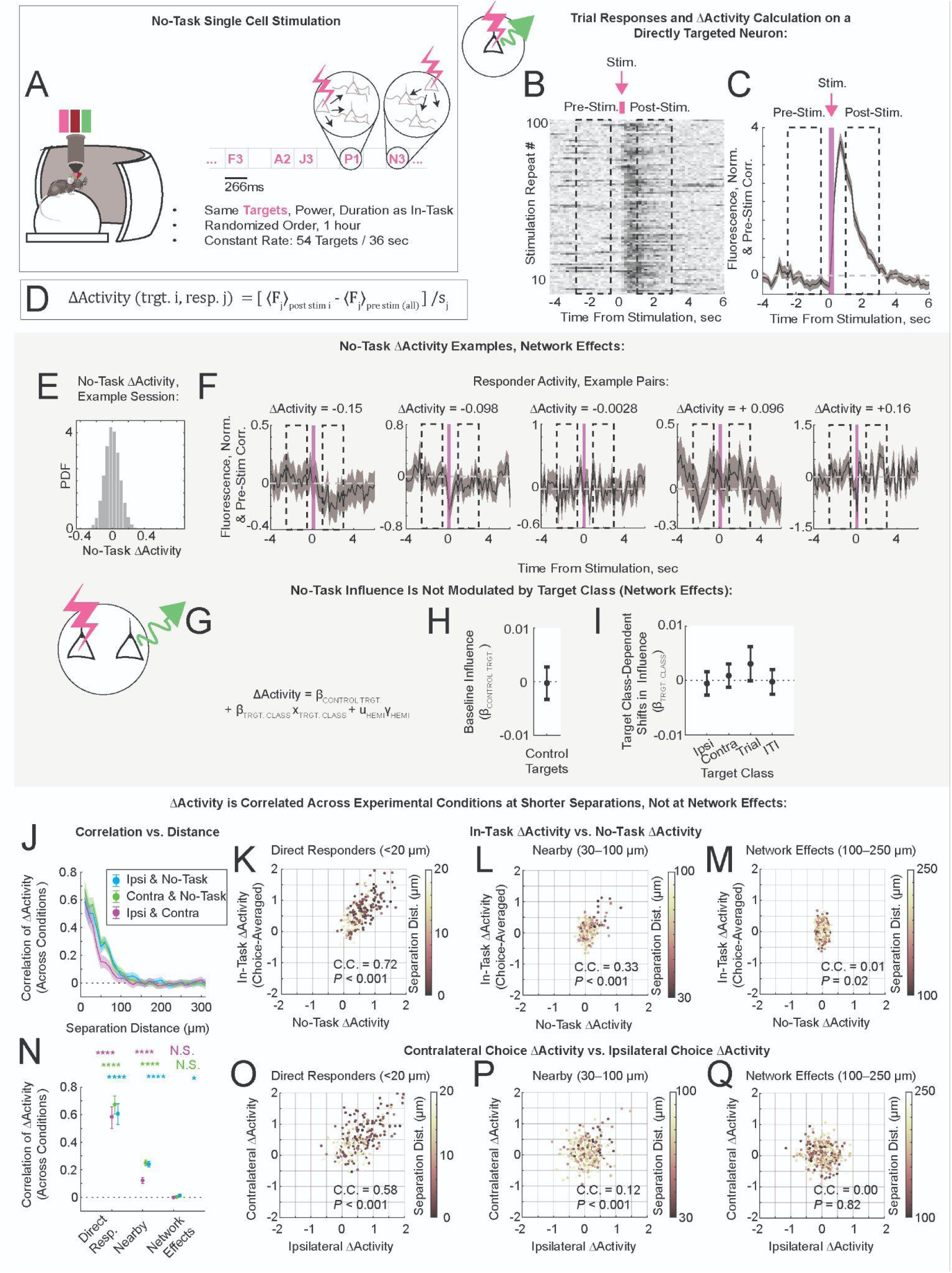
Measurement and Characterizations of No-Task Causal Connectivity, Related to Figures 1–4. In-task measurements of causal connectivity (Fig. 2) were followed with a no-task stimulation protocol in a subset of sessions (11 out of 20). **A.** Following the in-task stimulation session, mice remained head-fixed on the spherical treadmill, while the VR screen was dark. Optical stimulation parameters were identical to those used in the in-task stimulation protocol (target location, duration of exposure, power). Stimulation order was randomized in the following way: every 36 seconds, each target was stimulated once. The order of stimulation within each 36 second block was randomized within each block (Methods : No-Task Stimulation). **B,C**. As in Fig. 2, calculation of ΔActivity is illustrated using a directly stimulated neuron for clarity. **B.** Fluorescence of a responding neuron (heat map, inverted grayscale colorbar such that darker = more activity) in response to 100 stimulation repeats. The effects of brief 266ms long stimulation (magenta bar) were evaluated comparing the averaged activity in post-stimulation (1–3 seconds following stim, dashed box) and pre-stimulation (-2.5 to -0.5 seconds preceding stim, dashed box) periods. Note that the pre-stimulation periods were not target-specific and included all pre-stimulation time bins in the session. **C.** Averaged fluorescence in response to stimulation, with pre-stimulation, stimulation and post-stimulation periods indicated. Shaded area indicates STE across 100 stimulation repeats. **D.** Definition of no-task ΔActivity. Stimulations of targets closer than 25 μm to the responder were excluded from the calculation of the normalization coefficient. Since most stimulation targets are >> 25 μm away, the normalization coefficient was estimated over ∼5000 numbers per target-responder pair (54 targets, 100 stimulations each per session). **E.** No-task distribution of ΔActivity (network effects range). **F.** Examples of ΔActivity (shown: stimulation-averaged responder activity vs. time following stimulation). **G–I.** To check if there were target class-dependent shifts in influence (similar to the epoch-conditioned, target class-dependent shifts in Fig. 5E), we used a mixed-effects model (**G**) to estimate the dependence of influence on target class, using ΔActivity measured in the no-task condition. Neither the baseline influence (Control target class stimulation, **H**), nor the target class-dependent shifts in influence (**I**) significantly deviated from zero. **J–O**. Correlation coefficients of ΔActivity across experimental conditions. These are measures of: how consistent is the causal connectivity when measured across conditions? Perhaps surprisingly, ΔActivity measured during ipsilateral choices was less correlated with ΔActivity measured during contralateral choices, than with ΔActivity measured in the no-task condition. **J.** Correlation of ΔActivity vs. separation distance (20 μm bins). **K–M.** Choice-averaged in-task ΔActivity vs. no-task ΔActivity (individual pair = point), grouped by distance (**K**, <20 μm; **L**, 30–100 μm; **M**, 100–250 μm). For target-responder pairs in the nearby and network effects separation ranges, we subselected a random 500 pairs to plot. Correlation coefficient and corresponding significance value for all pairs indicated (# of pairs in panel K, 271; L, 8 555; M, 53 662 pairs). **N**. Correlation of ΔActivity vs. separation distance (same as panel J), grouped into the separation categories Direct Resp., Nearby, and Network Effects. **O–Q.** Comparison of ipsilateral to contralateral in-task ΔActivity. Same pairs, same organization as in panels K–M. Significance values in panels K–Q were multiple comparisons corrected (Bonferroni, 6X).

**Figure S4.**
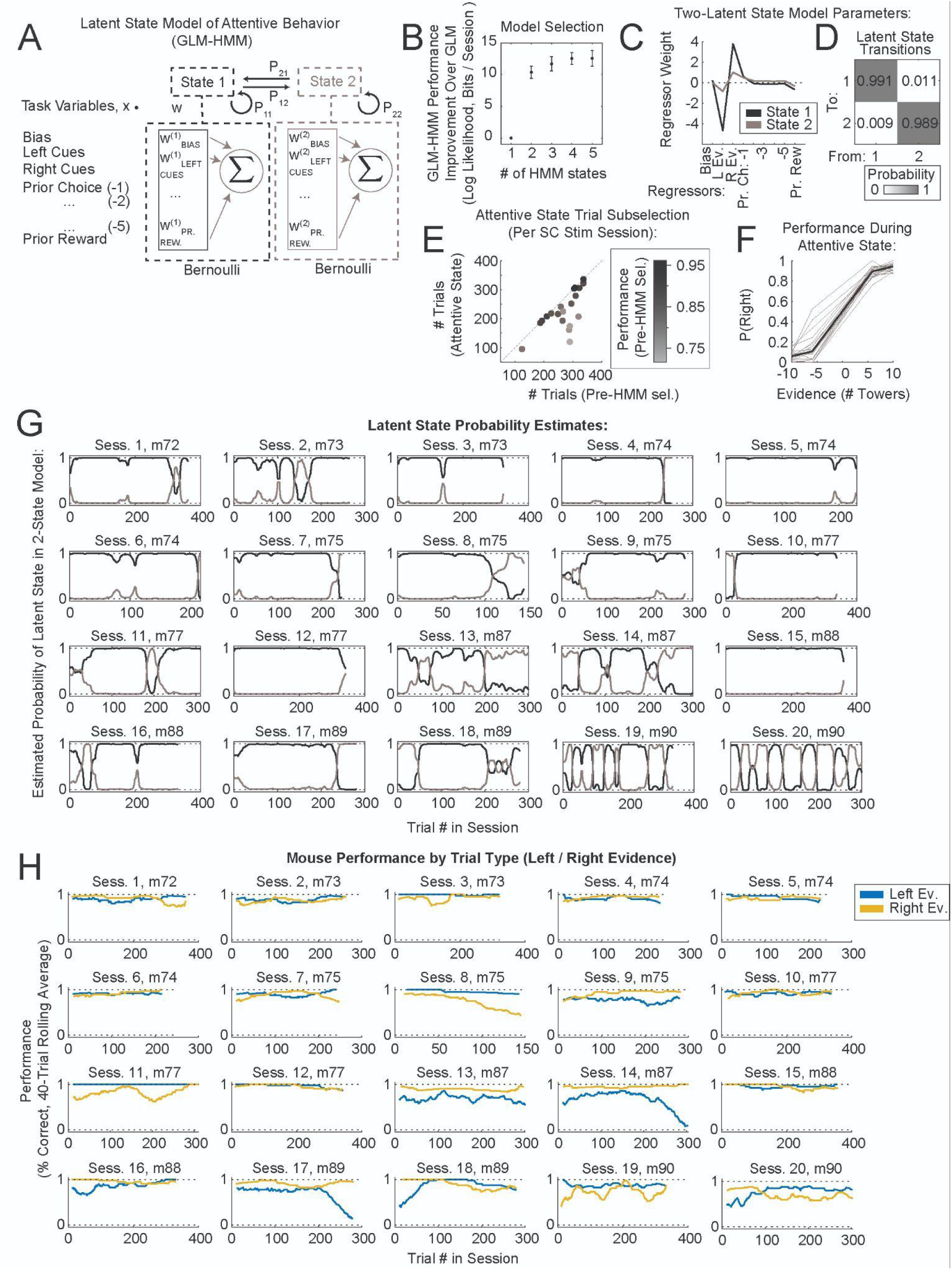
Attentive State Characterization Using a Two-Latent State GLM-HMM Model of Behavior, Related to Figures 3–5. **A.** Schematic of the latent state model of behavior implemented (see also Ref. ^9^). This model aimed to capture the dependence of animal choice on task variables (sensory evidence, prior choice) using multiple latent states (in this example, the regressor weights w within the black and gray squares have superscripts indicating the state), and a markov transition matrix between them. The model fitting procedure is described in Methods : Latent State Model of Task Engagement, and closely followed Ref. ^9^. **B.** Cross-validated performance of models vs. number of latent states. **C.** Regressor weights per state, for the two-latent state model. Prior choice (went-right / went-left) had five regressors corresponding to the number of trials back (i.e. -1 is the trial immediately preceding the current one). The two-latent state model is simple to interpret: state 1 is more attentive, state 2 is less attentive with more weight in the prior choice terms. **D.** Probabilities of transitions between the latent states, as estimated by the model. Low probabilities of a transition between different states reflect the slower time scales captured by the model. **E.** # of trials classified as attentive vs. total # of trials (individual points are sessions), colored by performance estimated pre-selection. **F.** Performance in the attentive state on stimulation sessions. Gray lines are individual sessions, black is median over sessions. **G.** Model-estimated probabilities of latent states, per session, shown for the twenty single-cell stimulation sessions analyzed in this paper. **H.** Mouse performance on all trials (prior to attentive state selection, averaged in a 40-trial rolling bin), per trial type, is shown for comparison to panel D.

**Figure S5.**
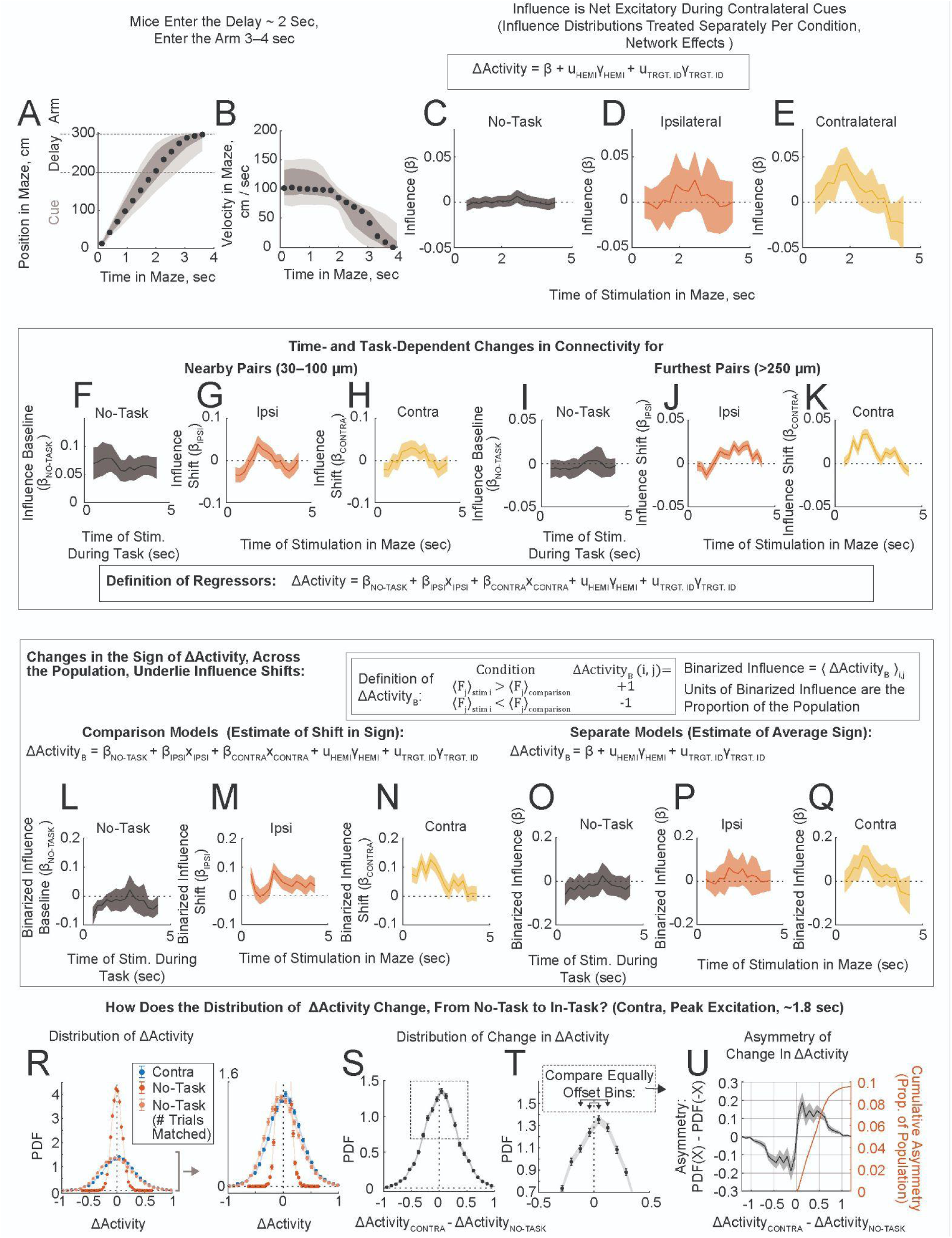
Supplemental Information Related to Figure 3. **A,B.** Median position (**A**) and velocity (along the stem of the maze, **B**), across sessions, in the maze. Position and velocity were evaluated at the stimulation onset times of targets (median across sessions, black dot, 2nd and 3rd quartiles dark shaded area, 95% range, light shaded area). Entry into the Delay was typically around 2 seconds from the start of the maze, the time point at which animals also started to slow down to navigate the upcoming turn. **C–E.** Influence (□), vs. time of stimulation in the maze, estimated using target-responder pairs in the network effects range (Model 1.3, Table 1). These regressors were estimated separately per experimental condition; data and conventions (shaded areas indicate 95% C.I.) are the same as in Fig. 3I–K. There were no significant trends in the no-task or ipsilateral influence. Peak deviation in contralateral influence (∼1.8 seconds in panel E) was □ = 0.042, \*\*\*\**P*_CORR_ = 6.1e-05 (Bonferroni 9X). **F–H**. Influence baseline (**F**) and task-dependent shift (**G,H**) vs. time of stimulation in the maze, analogous to the main text (Fig. 3I–K), for target-responder pairs nearby (30–100 μm, 8 hemispheres, 378 targets, 8 555 pairs). It is unclear from these panels: is this a noisy version of the same trends we observed in the network effects range? Or is this a common, choice-invariant trend? Because our sampling was greatly reduced at closer separations (and for the other reasons mentioned in the main text), we focused on the target-responder pairs at network effects separations throughout this paper. **I–K.** Influence baseline (**I**) and task-dependent shift (**J,K**) vs. time of stimulation in the maze, analogous to the main text (Fig. 3I–K), for target-responder pairs at greater separation distances (>250 μm, 8 hemispheres, 389 targets, 79 723 pairs). The trends in panels I–K look qualitatively similar to those reported in the main text, for network effects separations. **L–Q**. Analyses of the binarized ΔActivity and binarized influence (definitions in inset). We binarized ΔActivity to a +1/-1 notation (ΔActivity**_B_**). This quantity is advantageous because the units of the binarized influence are the proportion of the population and hence easy to interpret. The disadvantage is that the normalization of ΔActivity has been removed, and so noisy pairs contribute to estimates of the binarized influence on the same scale as less noisy pairs. **L–N.** Analyses of the binarized influence (network effects range; Model 1.4, Table 1, compare to Fig. 3I–K). **L.** Binarized influence baseline (estimated in the no-task condition) vs. time of stimulation in the maze. **M.** Shifts in binarized influence vs. time of stimulation in the maze, ipsilateral choice. At ∼1 sec, there is no significant change in the proportion of signed interactions (□ = 0.00, *n.s.*). At ∼1.8 sec, ∼9% of the proportion of pairs shifts excitatory (□ = 0.09, \*\*\*\**P*_CORR_ = 1.8e-10, Bonferroni 9X). **N.** Shifts in binarized influence vs. time of stimulation in the maze, contralateral choice. Both ∼1 and ∼1.8 sec have >10% shifts in the proportion of pairs that are excitatory (1 sec: □ = 0.12, \*\*\*\**P*_CORR_ = 6.3e-19; 1.8 sec: □ = 0.11, \*\*\*\**P*_CORR_ = 4.5e-15, Bonferroni 9X). **O–Q**. Binarized influence vs. time of stimulation in the maze; regressors estimated separately for the no-task (**O**), ipsilateral (**P**) and contralateral (**Q**) conditions. Data and approach (estimating regressors per condition) is the same as in panels C–E (Model 1.5, Table 1). The peak excitation during contralateral cue presentation (at ∼1.8 seconds, panel Q) corresponded to ∼11% more excitatory than inhibitory pairs in the population (□ = 0.11, \*\*\**P*_CORR_ = 5.4e-04, Bonferroni 9X). **R–U**. Changes in the distribution of ΔActivity, shown for peak contralateral offsets (∼1.8 seconds, stimulation bins 6 through 9, network effects, 11 200 pairs). **R.** Distributions of ΔActivity for in-task contralateral (blue), and no-task (red). The right panel provides an expanded view. Error bars are standard deviations over resampled pairs (with replacement). No-task ΔActivity was estimated by comparing 100 post-stimulation responses to 3–5 x 10^3^ pre-stimulation responses. In-task ΔActivity compared ∼20 stim to ∼20 comparison trials. We suspected that differences in sampling between no-task and in-task conditions were the leading cause of the different widths in the distributions of ΔActivity (compare red to blue). To check for this, we resampled no-task ΔActivity, downsampled to match the number of samples in the in-task measurement (N_STIM_, N_COMPARISON_). Specifically, we selected (1/N_STIM_ + 1/N_COMPARISON_)^-1^ of the available no-task post-stimulation responses (numbers matched per target-responder pair), and did not downsample the pre-stimulation responses (light red). Error bars for the downsampled estimates were standard deviations over 100X different post-stimulation resamples. Notice that this corresponds to a ∼10X downsampling of the no-task stimulation data. Following this correction for sampling, the widths of the distributions of ΔActivity became comparable between contra and no-task conditions (compare light red to blue), while the means remained offset. **S.** Distribution of the change in ΔActivity (contra - no-task), per pair. **T.** Expanded view of the distribution in panel S. **U.** Asymmetry in PDF (left y-axis) shows increased weight in positive values. The cumulative asymmetry (right y-axis) shows the excess proportion of the population (reaching 9.8%, right axis). This is similar to the coefficient of ∼11% estimated in panels N,Q.

**Figure S6.**
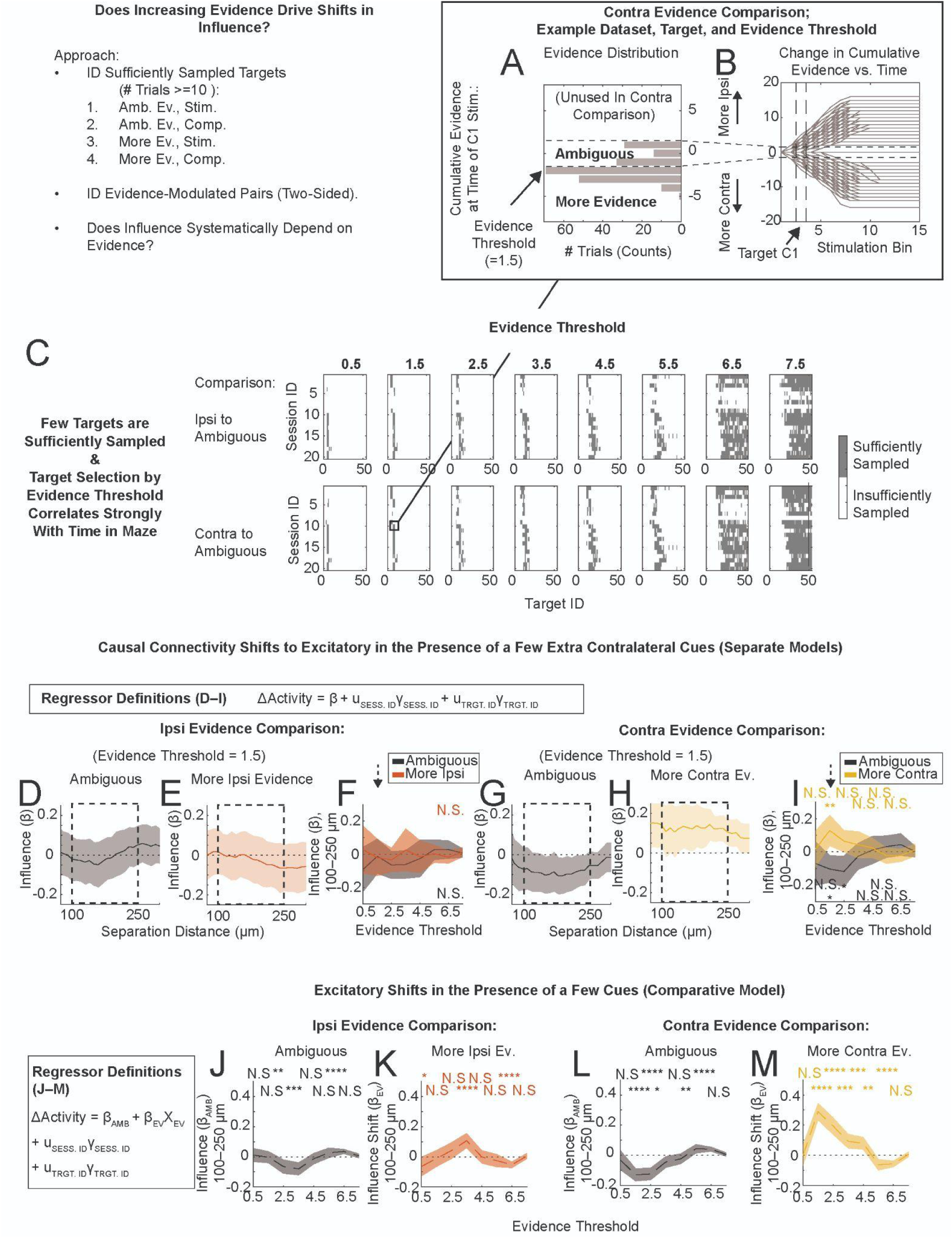
Contralateral Cues Drive Net Excitation, Related to Figure 4. We probed whether influence depends on evidence. This analysis was constrained to comparing ΔActivity across evidence conditions within target-responder pairs. This constraint constrains parameters related to navigation, such as the timing of stimulation relative to maze traversal. Because we were very limited in sampling with this analysis, we included all trials that passed the behavioral trial selection criteria (and did not condition trials by attentive state or correct outcome, as in all other analyses in this paper). **A,B**. We first identified the targets that were sufficiently sampled for comparison. At an evidence threshold of 1.5, target C1 in the 12th session was sufficiently sampled in both ambiguous and contralateral more-evidence conditions. **A.** Histogram of the instantaneous evidence at the time of C1 stimulation. Evidence threshold indicated with dashed lines. Trials with a cumulative two or more contralateral cues at the time of stimulation were classified into the “more evidence” trials; trials with a cumulative 0 or 1 cue (on either side) were classified as “ambiguous” trials. Trials with 2 or more ipsilateral cues were not used in this (contra) comparison. **B.** Instantaneous evidence vs. stimulation time bin for all trials in this session; stimulation bin of target C1 and evidence threshold are indicated with dashed lines. **C.** Targets that were sufficiently sampled per evidence threshold, shown per session, across comparisons. **D–M**. From the adequately sampled targets, we then identified target-responder pairs with significant evidence-dependent modulation of ΔActivity (by comparing to a resampled null distribution, Methods : Analysis of Evidence-Dependent Shifts in Causal Connectivity). The analysis of influence estimated using significantly modulated pairs is shown in panels D–M. **D–I.** Data was matched within comparisons (i.e. panels D and E are calculated using the same target-responder pairs), regression models were fit separately per condition (Model 2.0, Table 1). **D,E.** Influence vs. distance, 100 μm rolling bin, for ambiguous (**D**) and more ipsilateral evidence (**E**) conditions, evidence threshold = 1.5; 17 sessions, 47 targets, 789 significantly modulated pairs. **F.** Influence (network effects) vs. evidence threshold, ipsilateral comparisons. **G,H.** Influence vs. distance, for ambiguous (**G**) and more contralateral evidence (**H**) conditions, evidence threshold = 1.5; 18 sessions, 52 targets, 697 pairs. **I.** Influence (network effects) vs. evidence threshold, contralateral comparisons. **J–M**. Estimates of evidence-dependent shifts in influence (network effects) vs. evidence threshold. Same data as in panels D–I, regression models (inset) were fit to both conditions to capture evidence-dependent offsets (Model 2.1, Table 1). **J,K**. Influence baseline (*β*_AMB_, **J**) and evidence-dependent shift (*β*_EV_, **K**) in the ipsi evidence comparison, vs. evidence threshold. **L,M**. Influence baseline (*β*_AMB_, **L**) and evidence-dependent shift (*β*_EV_, **M**) in the contra evidence comparison, vs. evidence threshold.

**Figure S7.**
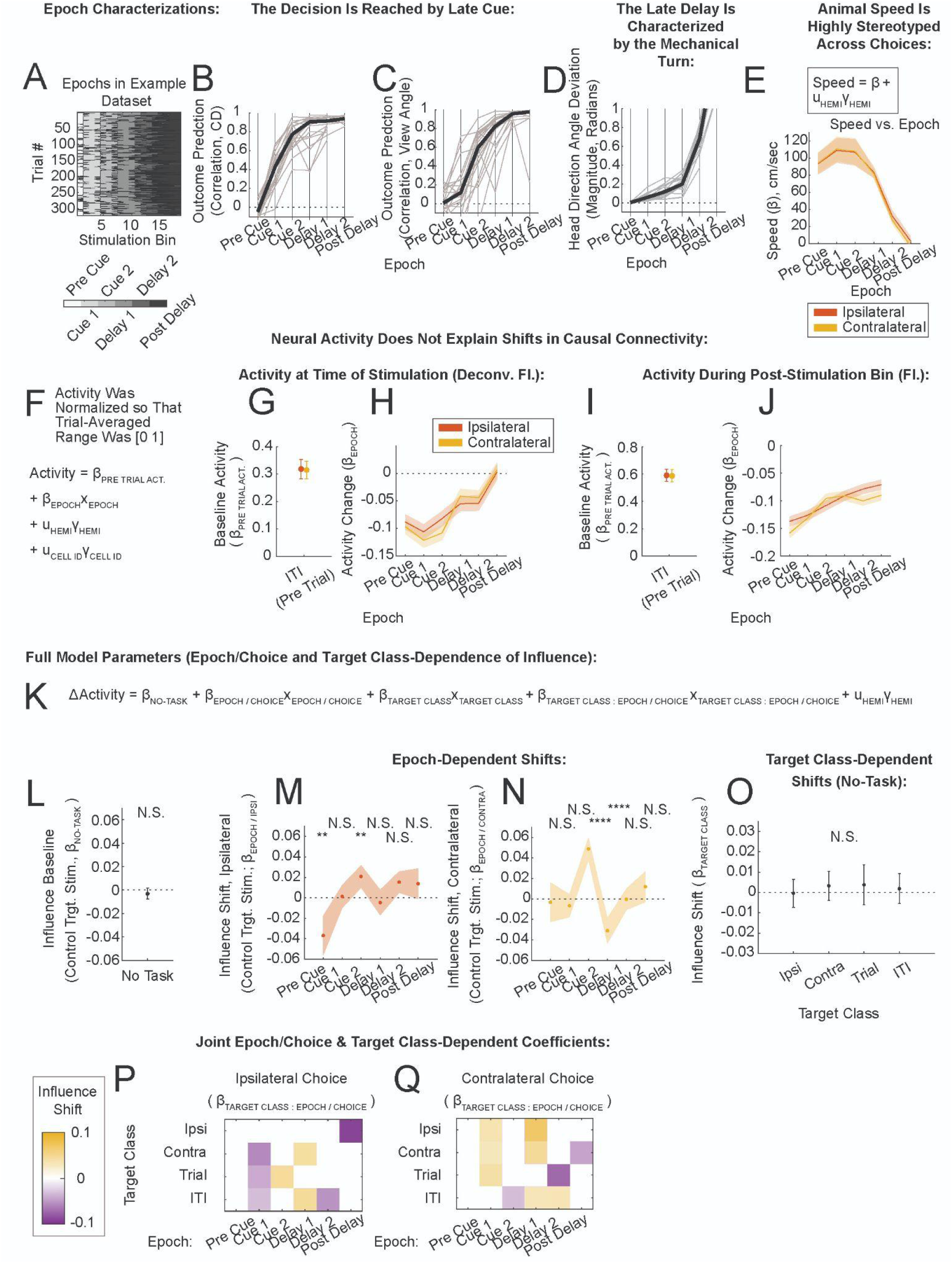
Epoch Characterizations and the Full Task-Dependent Model of Influence, Related to Figure 4. **A.** Epoch definitions, evaluated per stimulation bin per trial (shown is an example session). The Cue period was identified on each trial, using the first and last stimulation bins within which the number of cues on the right or left hand side was changing. This time period was split in half to define the Early Cue and Late Cue epochs. The Delay period was similarly identified immediately following the Cue period until arm entry, and similarly split in half by time to define the Early and Late Delay epochs. **B.** Correlation of the choice-coding projection with outcome, per epoch (gray lines are individual sessions, black is median). To calculate this, we randomly selected half of the correct went-right and correct went-left trials to calculate the z-scored choice preference per neuron per time bin (a matrix N_CELLS_ X N_TIME_ in size). We then projected the withheld-trial fluorescence onto the z-scored choice preference to estimate the choice-coding projection, per trial per time bin. To convert time-dependence to epoch-dependence, we averaged the projection over time bins belonging to the same epoch (within each trial), to obtain the choice-coding projection, per trial per epoch. This was then correlated with outcome. **C.** Correlation of head view angle with choice across epochs. In this case we averaged head direction angle within epoch first, to obtain head direction angle per epoch per trial. Similarly to panel B, we used a randomly selected half of the correct went-right and correct went-left trials to calculate the average trend in head direction angle vs. epoch; projected the withheld-trial head direction angle onto this trend, and correlated the projection with outcome. **D.** Absolute value of the head direction angle (radians) vs. epoch. **E.** Velocity (down the stem of the T) across animals, ipsilateral and contralateral choices in red and gold, respectively, estimated as the fixed-effects coefficient in a mixed-effects model (inset). Shaded areas are 95% C.I. Running speeds were indistinguishable across choices in early epochs, until the Post-Delay epoch (e.g. two sample t-test, ipsi vs. contra, Early Cue, n.s.; Post-Delay, *P* = 0.02. Full statistics in Table S1). This comparison indicates that running speed could not explain choice-dependent differences in causal connectivity (which were found primarily during the Early Cue). The time courses of running speed and task-dependent shifts in connectivity also did not correlate cleanly: Running speed was consistently high from Pre Cue through Late Cue; while shifts in connectivity started during Cue onset, peaked in the Late Cue and were significant in the Early Delay. **F–J.** Trends in neural activity vs. epoch, evaluated on attentive, nonstimulation trials. **F.** Definitions of regressors in the epoch-dependent models of neural activity. Activity within-epoch was treated as the sum of a baseline term (measured within the ITI preceding trial start), and an epoch-dependent shift. Prior to this analysis, neural activity was normalized within each neuron across trials so that the trial-averaged range across time bins would be in the 0 to 1 range per neuron. As elsewhere, shaded areas are 95% C.I. **G,H**. Neural activity baseline (**G)** and epoch-dependent shifts (**H**). We used deconvolved fluorescence here to improve the time resolution of the estimate. **I,J**. Neural activity baseline (**I)** and epoch-dependent shifts (**J**), evaluated in the post-stimulation time bin (the same time bin used to calculate influence, i.e. 1–3 seconds following the epoch). Here we used the raw fluorescence for consistency with the calculation of influence. In all relevant trends of neural activity (panels G–J), neural activity was minimized in the Pre Cue / Early Cue, and rose gradually as the animals progressed through the maze. These results are similar to those obtained from population averages over the time-dependent encoding terms in single neuron GLMs (Fig. S2I). These trends in population-level activity do not clearly correlate with task- or choice- dependent changes in connectivity (Figs. 3–5). Speculatively, epoch-dependent changes in the global activity could play a role in setting the operating points of inhibitory interneuron circuitry, which could lead to changes in causal connectivity (see Fig. 6, Discussion). **K–Q**. The features and estimates of a comprehensive mixed-effects model of influence measured in both no-task and in-task conditions (Model 3.1, Table 1). Dots indicate estimates; asterisks indicate significance values following multiple comparisons corrections (30X, Bonferroni); and shaded areas indicate uncorrected 95% C.I. returned by the model. **K.** Definitions of regressors. **L**. Influence baseline (estimated from Control target stimulation in the no-task condition). **M,N**. Influence shifts due to ongoing epoch/choice (estimated from Control target stimulation). The coefficients describing Early Cue and Late Cue are those reported in the main text panels Fig. 4J,L. **O**. Influence shifts due to target class (estimated in the no-task condition). These results are similar to the results of no-task only influence analyses in Fig. S3I. **P,Q**. Influence shifts due to the combination of target class (row) and epoch/choice (column), shown as a heat map. Non-significant coefficients (*P*_CORR_ > 0.05, Bonferroni 30X) are shown as white. The columns of coefficients corresponding to Early Cue and Late Cue are shown in Fig. 4K,M.

**Figure S8.**
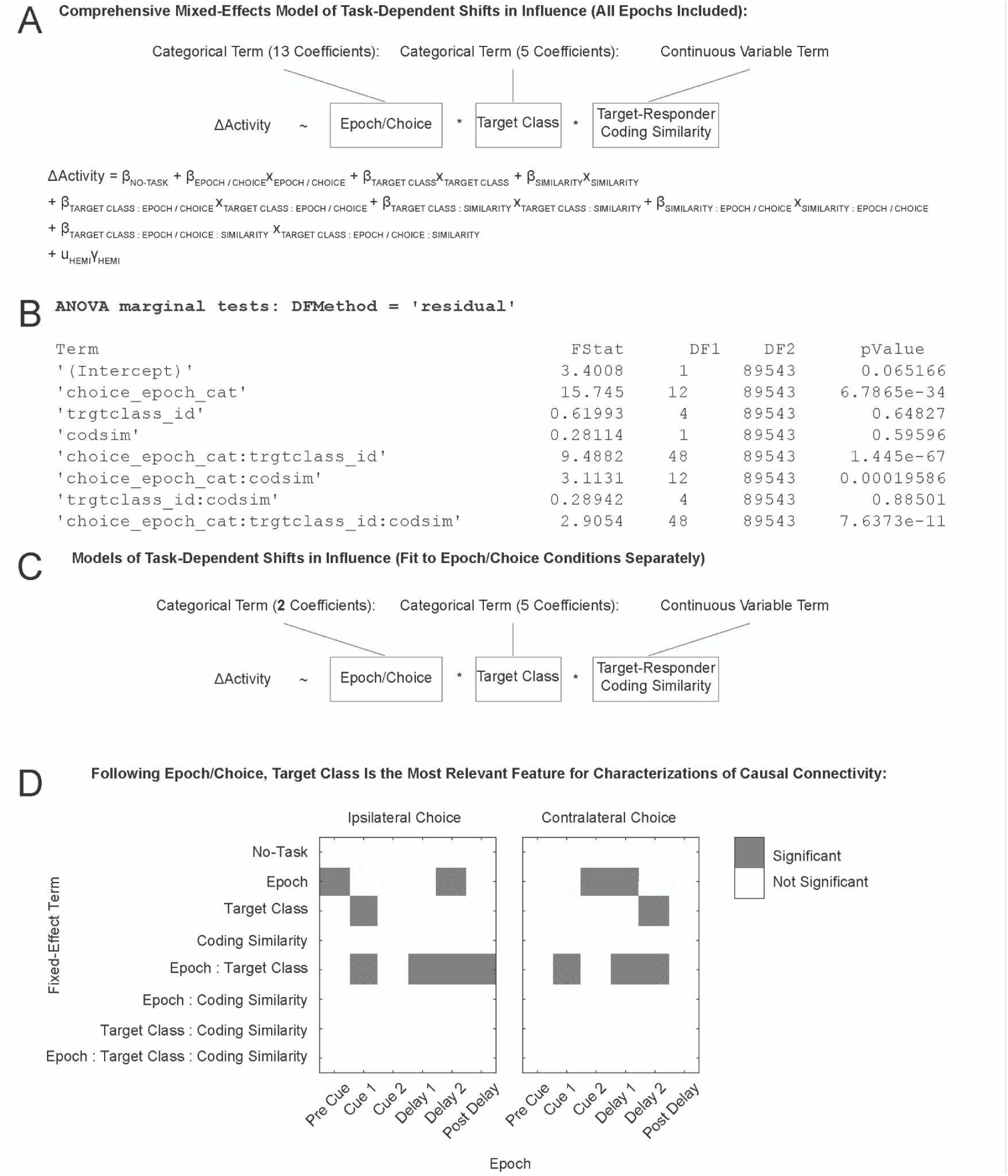
Feature Selection in Mixed-Effects Models of Influence, Related to Figure 4. **A**. Description of the comprehensive mixed-effects model that incorporated all terms up to the joint three-way interaction term between epoch/choice, target class, and coding similarity (130 coefficients). Epoch/choice was parameterized with 13 coefficients (one no-task baseline, and twelve offsets for the 6 epochs * 2 choices). Target class was parameterized with 5 coefficients (one Control target stimulation baseline and four target class specific offsets). Coding similarity, a continuous variable, was parameterized with 1 coefficient. **B.** Results of the marginal tests (*F*-tests) on the fixed-effects terms (Matlab’s **glme.anova**), which test whether the coefficients within each fixed-effect term significantly deviate from zero. **C**. Description of the mixed-effects models estimated per epoch/choice condition. In contrast to panel A, two coefficients were necessary for epoch/choice, one to estimate the no-task baseline, and the other to estimate the epoch/choice-dependent shift. **D**. The significance of terms which modulated the influence, calculated within epoch/choice conditions separately (panel C), evaluated with marginal tests (as in panel B). These were Bonferroni corrected (12 X), and reported in this panel if *P*_CORR_ < 0.05. During the Early Cue, on ipsilateral trials, coding similarity was not significant (*F*-statistic = 0.01, *n.s.*) while the joint target class : epoch term was (*F*-statistic = 13.4, \*\*\*\**P*_CORR_ < 0.0001). Similarly for contralateral trials (coding similarity *F*-statistic = 0.04, *n.s.*; target class : epoch *F*-statistic = 6.4, \*\*\**P*_CORR_ < 0.001). During the Late Cue causal connectivity was not modulated by target class (Epoch : Target Class term, ipsilateral choice trials, *F*-statistic = 2.3, *n.s.*; contralateral choice trials, *F*-statistic = 2.4, *n.s.*; Bonferroni 12X). Thus ongoing epoch/choice and target class modulated influence most consistently, whereas coding similarity did not modulate influence significantly in any of the epochs.

**Figure S9.**
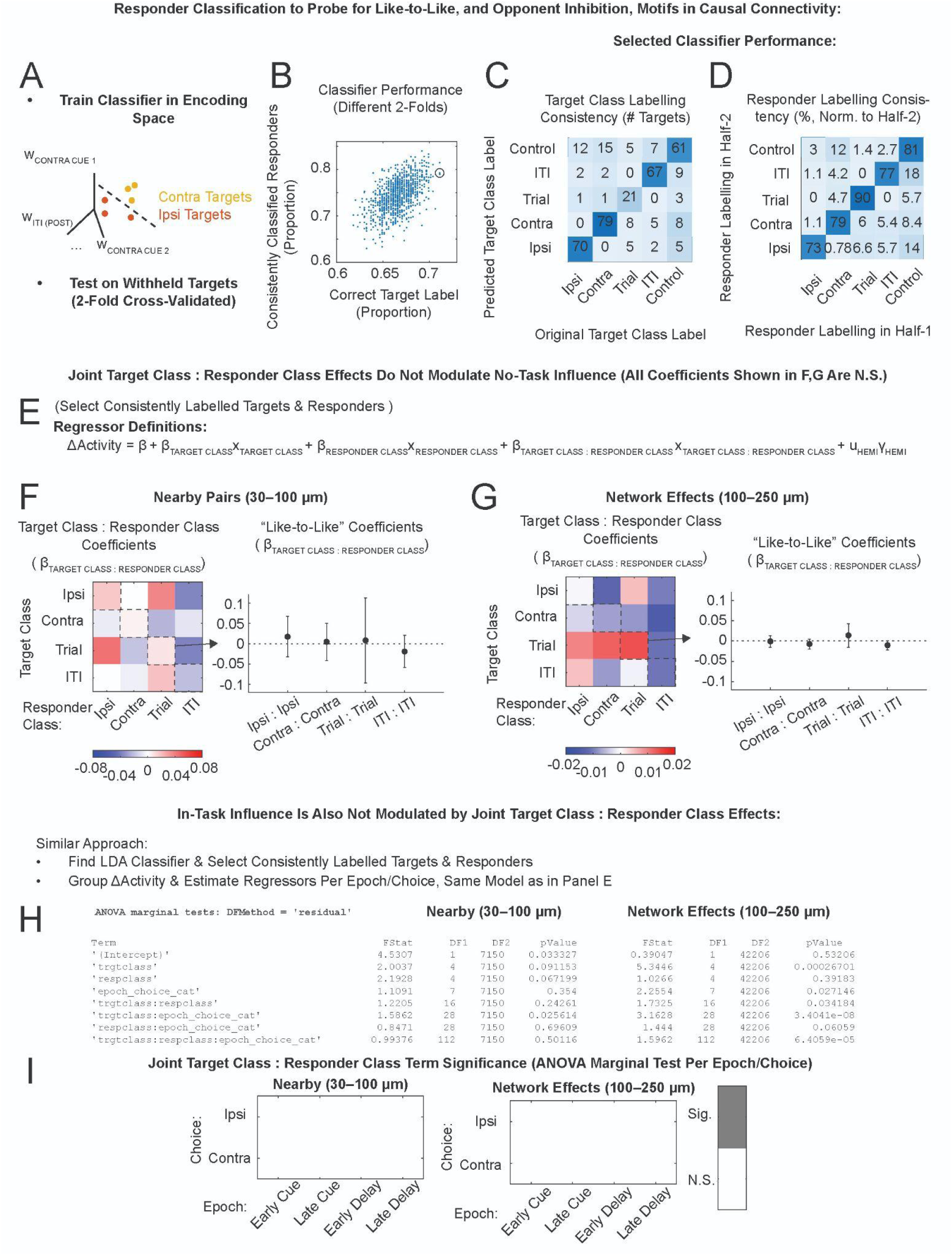
No Evidence of “Like-to-Like” or “Opponent Inhibition” Motifs in the Causal Connectivity, Related to Figures 4 and 5. We classified responder neurons into the same encoding classes as the targets (Ipsi, Contra, Trial, ITI and Control), and then probed whether terms describing joint effects from target class and responder class modulated influence. Joint effect terms did not statistically deviate from zero. Additionally, the inferred coefficients were not consistent with like-to-like (positive terms between the same target and responder class) or opponent inhibition (negative terms between different classes, such as Ipsi targets : Contra responders) motifs. The approach and results are illustrated in greater detail for the no-task data (11 sessions, panels A–G), and then summarized for the in-task data (20 sessions, panels H–J). **A–D**. Approach to responder categorization. **A.** We estimated linear discriminant classifiers that separated the original target classes in the 21-dimensional regressor space of the single neuron GLMS (Fig. S2), using a 2-fold crossvalidated approach. In each fold, we tested labelling predictions on the withheld half of targets. We also used both folds to label responders and measure the consistency of responder labelling. **B.** We repeated the data split and classification 1000X and selected the data split (indicated with black circle) that maximized the proportion of correctly labelled targets (x-axis). The two classifiers estimated from separate halves of this data split classified responders with ∼80% similarity (y-axis). **C.** Confusion matrix of target class labels in the selected classifier. **D.** Similarity of responder labelling in the selected classifier, across halves, normalized by row. Total number of diagonal elements (from Ipsi to Control): 7080, 8566, 2417, 11122, 35284. For subsequent analyses, we only included targets that were correctly classified, and responders that were consistently classified across the two halves of the data (i.e. diagonal elements in panels C and D; 46 228 pairs, or 45%, passed this criteria). These were then grouped by separation distance below. **E.** Definitions of regressors. **F,G**. Influence shifts due to the combination of target class and responder class. All of the coefficients shown in panels F and G were not significant; they are shown here to illustrate the lack of strong positive diagonal elements (like-to-like), as well as the lack of strong negative off-diagonal elements (opponent inhibition, e.g. between Ipsi and Contra). **F.** Influence shifts estimated using target-responder pairs nearby (30–100 μm, ANOVA: *F*-statistic = 0.85, *P* = 0.6). This data from 8 hemispheres, 295 targets, 3410 pairs. Left panel: All joint target class : responder class coefficients estimated by the model in panel E, shown as a heat map (row: target class; column: responder class). Right panel: Coefficients describing the interaction between like-to-like elements (diagonal of the left panel). Error bars are 95% C.I. **G.** Influence shifts estimated using target-responder pairs at network effects separations (100–250 μm, ANOVA: *F*-statistic = 1.4, *P* = 0.15). This data from 8 hemispheres, 298 targets, 19 496 pairs. Left and right panels as in panel F. **H–J**. To similarly probe in-task influence for evidence of like-to-like motifs, we repeated the classification steps in panels A–D using the full 20-session dataset. This yielded a classification with [13072, 11666, 5748, 29518, 33608] consistently identified responders in the five categories as above; a total of 62 749 target-responder pairs (43%) passed correct-target and consistent-responder classification criteria. We then subselected the four best sampled epochs, Early Cue to Late Delay, to perform the analysis within epoch/choice conditions. This seemed the most appropriate approach given our primary finding in this paper, which is that ongoing computation is the most relevant feature in characterizations of causal connectivity. We next searched for the 3-way joint effect between epoch, target class and responder class (panel H) and then subdivided the data by epoch/choice and searched for 2-way joint effects in target class and responder class (panel I). The data was sampled from 11 hemispheres, 274 targets, 3 657 pairs (30–100 μm), and 11 hemispheres, 279 targets, 21 203 pairs (100–250 μm). **H.** Statistics of the terms in the comprehensive mixed-effects model (ANOVA marginal tests), for nearby (left columns) and network effects (right columns) pairs. The joint 3-way term between target class, responder class, and epoch/choice was found to be statistically significant. However, individual coefficients (not shown) were not significant following multiple comparisons correction (30X). Also it is noteworthy that of the three 2-way interaction terms, target class : epoch was highly significant, while the other two terms were borderline, or not at all, significant. This suggests caution in interpreting the significance of the 3-way term naively, and that the significance of the joint 3-way term may be inherited from the significance of the 2-way target class : epoch term. For this reason we subsequently performed similar analyses within separate epoch/choice conditions (panel I). **I.** Significance of the joint target class : responder class term (ANOVA marginal tests), evaluated per epoch/choice condition separately. The joint term did not significantly deviate from zero in any of the conditions (all *P*_CORR_ > 0.05, Bonferroni, 12X). This was true for both nearby (left panel) and network effects (right panel) pairs.

**Figure S10.**
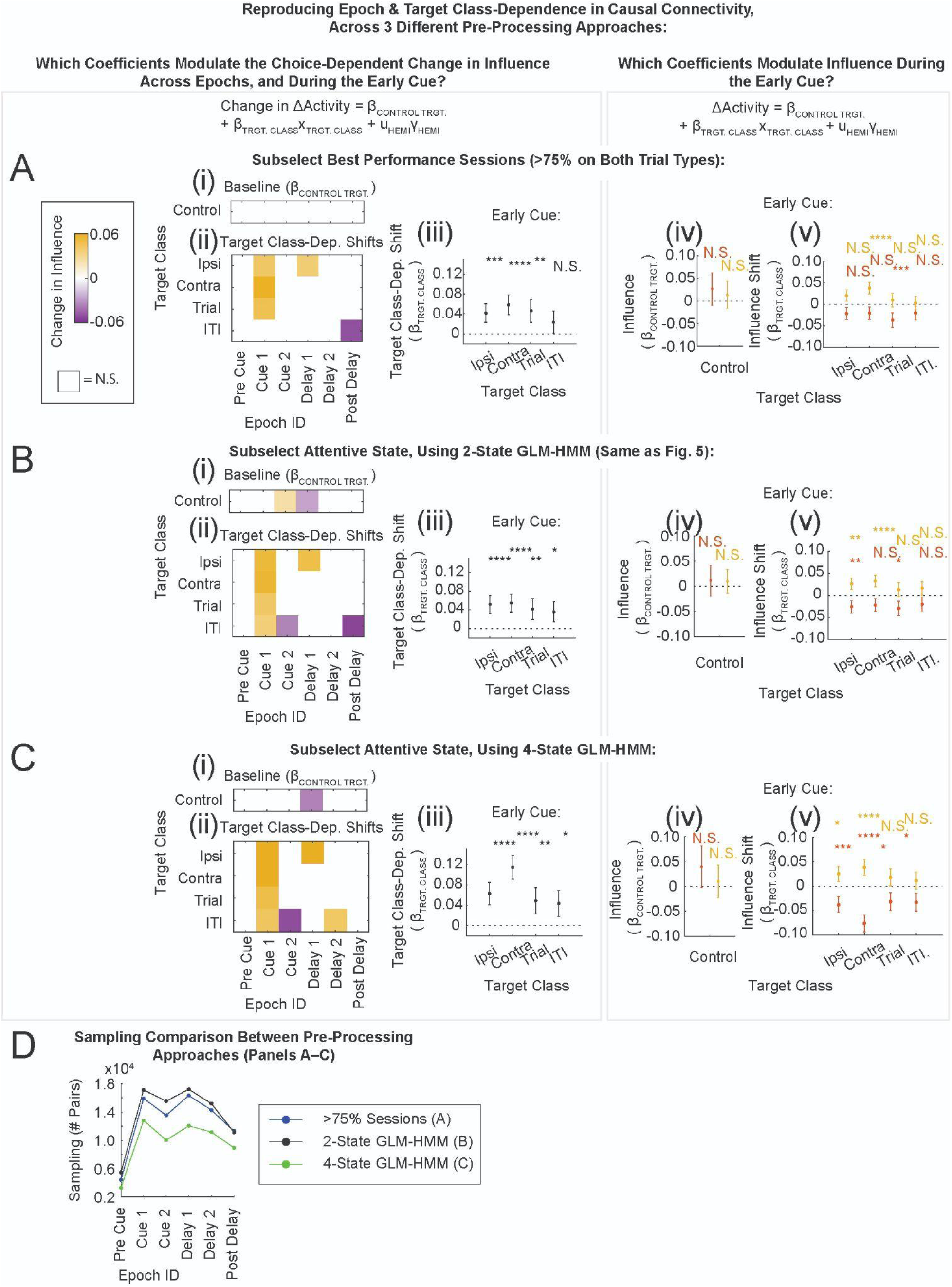
Target Class & Epoch-Dependent Shifts in Influence Are Consistent Across Attentive State Definitions, Related to Figure 5. To get a sense for how strongly our results depended on choices of preprocessing, we repeated the analyses of epoch & target class dependencies in influence following two other preprocessing approaches. For a particular set of selection criteria (panels A–C, described below), we evaluated the dependence of the choice-dependent change in influence on target class, per epoch (left column, compare to Fig. 5D–F). We also reevaluated the dependence of influence on target class, per choice condition during the Early Cue (right column, compare to Fig. S11B). All significance indicators were corrected for multiple comparisons as in the main text. **A.** We selected high performance sessions (mice ran >75% on both right- and left-trial types), and kept all non-aberrant post warm-up trials as described in Methods: Behavioral Trial Selection. **B.** We used the two-state GLM-HMM (Fig. S4) to select the attentive state (reproduced from Fig. 5 and the main text). **C.** We used a four-state GLM-HMM to select a higher-performing attentive state than in the two-state GLM-HMM model. In all three approaches (panels A–C), Control target stimulation did not significantly modulate the choice-dependent change in influence during the Early Cue (i, left column). Target class modulated the choice-dependent change in influence primarily during the Early Cue, and not in other epochs (ii, left column). The effects from Ipsi, Contra and Trial target stimulation were most consistent, while the effect from ITI target stimulation was borderline (iii, left column). Examining the influence by choice condition separately, Control target stimulation was not significant on ipsilateral or contralateral choice trials (iv, right column). Target class-dependent shifts in the influence were positive on contralateral trials (v, right column, gold) and negative on ipsilateral trials (v, right column, red). **D.** Sampling across epochs for the three different pre-processing approaches shown in panels A–C.

**Figure S11.**
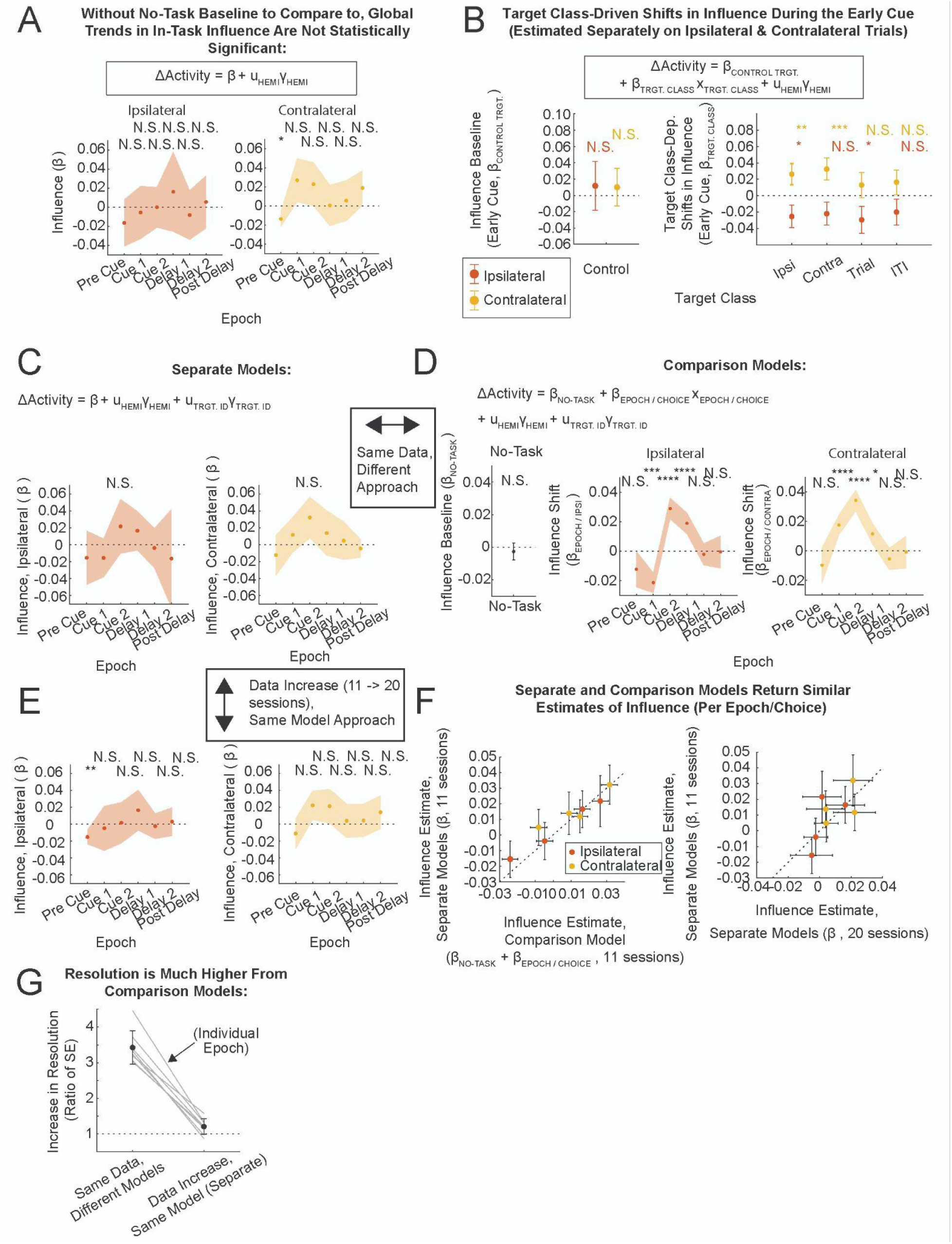
Supplemental Information Related To Figure 5. **A.** Influence per epoch/choice, calculated with separate models per condition (Model 4.1, Table 1: asterisks indicate 12X Bonferroni corrected significance values). **B.** Target class-dependent shifts in the influence, relative to Control target stimulation, during the Early Cue (Model 4.2, Table 1: asterisks indicate 60X Bonferroni corrected significance values). Note that some of these target class-dependent shifts were significant even though averaged trends in influence were not significant without the no-task comparison (panel A). These shifts were estimated separately within ipsilateral (red) and contralateral (gold) choice conditions. Differences between conditions (e.g. within Ipsi targets in the right panel) were similar to the absolute choice-dependent changes in influence reported in Fig. 5F; the number of random-effects coefficients was doubled in this analysis. **C–G**. Models that estimated shifts in influence (e.g. Fig. 3I–K, Fig. 4C–E) had a much lower uncertainty on the estimate of the shift, than the uncertainty that we obtained in estimating absolute values (e.g. panel A here, or Fig. S5C–E). The drawback is that comparison models can not probe whether the absolute value of influence is net-excitatory or net-inhibitory. Here we quantified the reduction in uncertainty across different approaches, to clear up any confusion about the data. **C.** In-task influence vs. epoch, estimated per epoch/choice (same data as Model 3.0, Table 1). **D.** Influence baseline (no-task, left panel) and task-dependent influence shifts (ipsilateral choice, middle panel; contralateral choice, right panel), reproduced from Fig. 4C–E. These terms were estimated using the same data as in panel C. Note the similar epoch-dependent trends, and the significant reduction in uncertainty, compared to panel C. **E.** In-task influence vs. epoch, estimated identically to panel C, using all the in-task data (same data as panel A; Model 4.1, Table 1; note the additional random-effects term compared to panel A). **F.** Estimates of the in-task influence, comparing the different approaches in panels C–E. Each point is one of the four Cue / Delay epochs. Dashed line is unity: the different approaches yielded similar estimates. Left panel: influence estimates from panels C (y-axis) and D (x-axis). Right panel: influence estimates from panels C (y-axis) and E (x-axis). **G.** Comparison of reduction in uncertainty between switching models with the same data (left side of panel; comparing panels D to C; median, 3.3x; range, 3.0–4.5x) and keeping the same model while increasing data sampling (right side of panel; comparing panels E to C; median, 1.2x; range, 0.9–1.6x; data increase from 8 to 11 hemispheres, 300 to 590 targets, 43 883 to 81 734 pairs). Each point is a choice-dependent Cue or Delay epoch, black is average, error bars are standard deviation over epochs.

**Figure S12.**
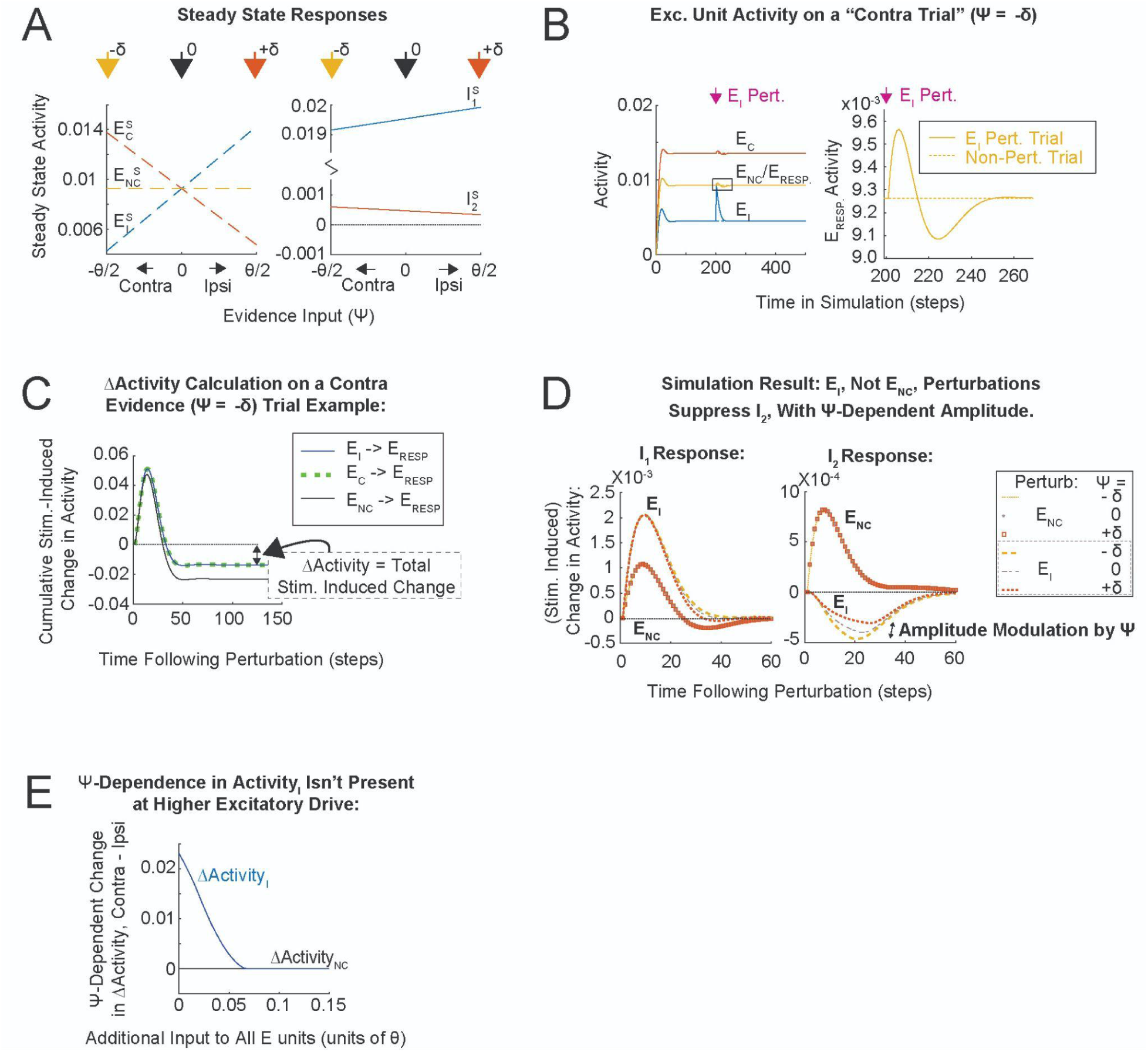
Simulation of the Conceptual Model, Related To Figure 6. **A.** Steady state responses of excitatory (left panel) and inhibitory (right panel) units, in response to varying the evidence input (*Ѱ*). In this model, the low activity levels in I_2_ were key for *Ѱ-*dependence in ΔActivity. **B.** Excitatory unit activity during contra evidence (*Ѱ*<0) trials. Full simulation (left panel), and zoom in on the response of E_RESP_ following stimulation (right panel) on an E_I_-perturbation trial (solid line) and non-perturbation trial (dashed line). The difference between these two is the stimulation-induced change in Activity. The time-dependent decay following a response to perturbation reflects the underdamped regime: pathways involving higher numbers of synapses contribute progressively less to ΔActivity. **C.** Cumulative stimulation-induced change in the activity of E_RESP_. In the simulation, ΔActivity is defined as the total difference between stimulation and non-stimulation trials (summed over time), which can be visualized on this plot as the value of the cumulative as t -> infinity. By construction, ΔActivity following E_I_ (solid blue) and E_C_ (dashed green) perturbations is the same and different to ΔActivity following E_NC_ perturbations (solid black). **D.** Stimulation-induced change in the activity of I_1_ and I_2_, in response to the two different perturbations in the three different *Ѱ* conditions. Left panel: I_1_ was excited by both types of perturbations (E_I_ and E_NC_). Right panel: I_2_ was excited by E_NC_ perturbations and suppressed by E_I_ perturbations. As suggested by the schematic in Fig. 6F, the magnitude of the I_2_ response to E_I_ perturbations was *Ѱ*-dependent. **E.** *Ѱ*-dependent change in ΔActivity, evaluated at *Ѱ* = +/- δ, vs. additional input to all excitatory units. Additional input of 0 is the default setting (in panels A–D). Providing additional input to all excitatory units moved I_2_ away from threshold, and removed the *Ѱ*-dependence in ΔActivity_I_ (blue line). ΔActivity_NC_ (black line) shown for comparison.

**Figure S13.**
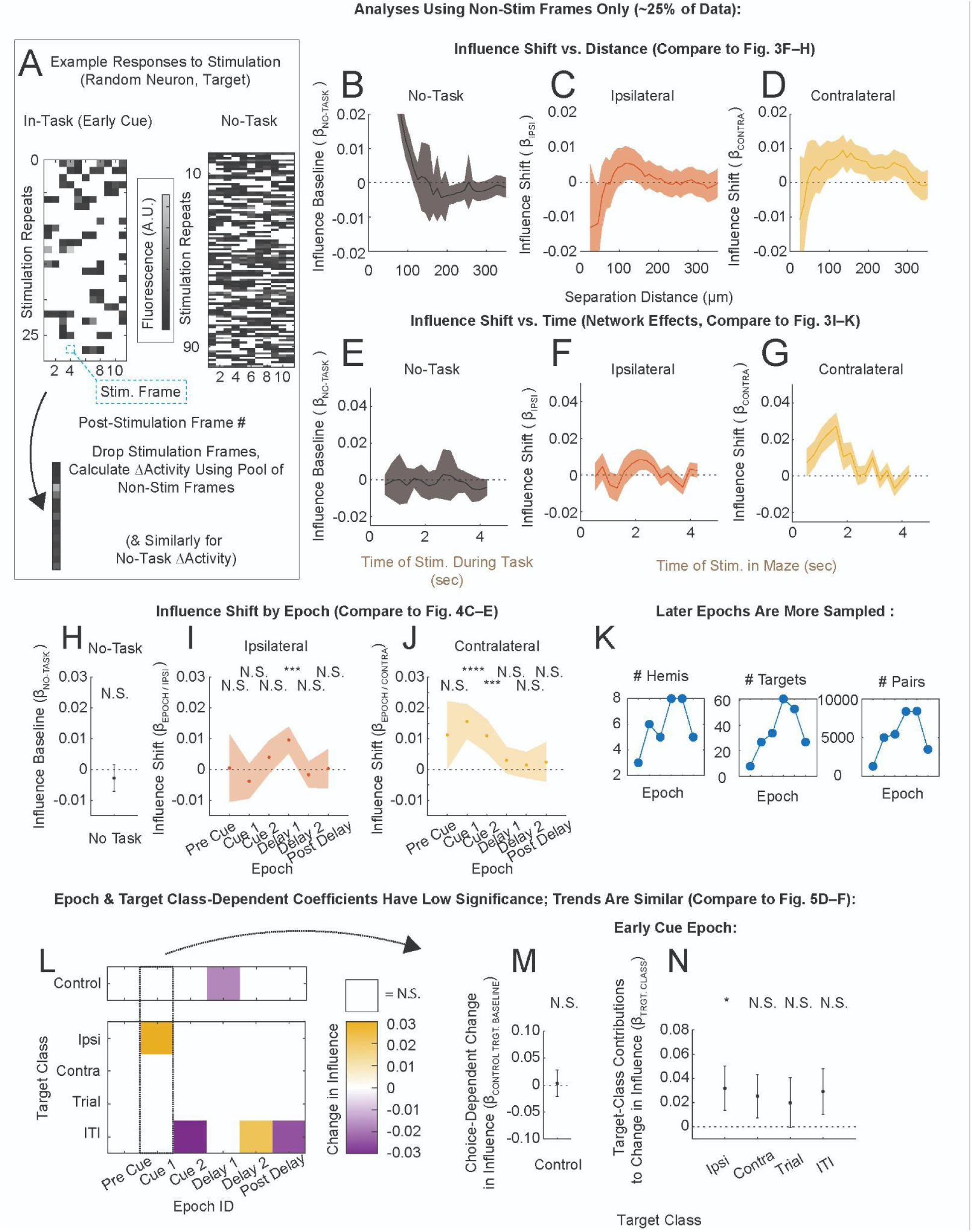
Properties of Causal Connectivity Estimated Using Only Non-Stimulation Imaging Frames Are Consistent With the Rest of the Paper, Related to Figures 3–5. In these analyses we explicitly removed imaging frames with simultaneous stimulation (∼75% of the recorded data in some cases), and recalculated the main features of influence presented in Fig. 3–5 (Methods : Stimulation-Induced Artefacts During Imaging). **A.** Fluorescence of a responder neuron in response to stimulation of a particular target (in-task example, left panel; no-task, right panel). Here, frames with simultaneous stimulation were set to white. We subselected imaging frames without simultaneous stimulation in the 1–3 seconds following stimulation (the same time limits as in other analyses in this paper). These were concatenated into a pool of post-stim frames. In this approach the stimulation and comparison responses, as well as the normalization coefficient, were calculated with respect to one frame. In all other analyses in this paper we calculated these quantities within the 2-second post-stimulation average (11 frames). These changes scaled the magnitude of the normalization coefficient, and subsequently the influence. Additionally, we replaced the sufficient sampling criterion based on trials (at least 10 stim and 10 comparison trials) with a criterion on the number of non-stimulation frames (at least 40 following stim and 40 in the comparison condition). Otherwise, the analyses here were identical to those in the rest of the paper. Because the stimulation rate on the timescale of a trial was different between in-task and no-task experiments (there was no stimulation during the ITI in-task, Methods : In-Task Stimulation), there were more non-stimulation imaging frames following stimulation in the no-task experiment. In-task, the response to stimulation during the early epochs overlapped with stimulation on 75% of trials (see also panel K). **B–D**. Influence baseline (**B**) and influence shifts (**C,D**) vs. distance, compare to Fig. 3F–H. **E–G**. (Network Effects) Influence baseline (**E**) and influence shifts (**F,G**) vs. time of stimulation, compare to Fig. 3I–K. **H–J**. (Network Effects). Influence baseline (**H**) and influence shifts (**I,J**) vs. epoch, compare to Fig. 4C–E. **K**. Sampling of hemispheres, targets and pairs used in panels H–J. In these analyses, the Pre Cue, Early and Late Cue epochs had ∼25% of the data available to estimate ΔActivity. Because ΔActivity is evaluated in the 1–3 second bin following stimulation, later epochs have an increased proportion of non-stimulation imaging frames. Estimates of ΔActivity in the Post-Delay were calculated using imaging frames following completion of the stimulation sequences and were essentially unaffected by the subselection in this analysis. **L–M**. Epoch & target class-dependent analysis of the choice-dependent change in influence, compare to Fig. 5D–F. **L**. Shifts in the choice-dependent change in influence due to target class-specific stimulation, estimated per epoch, shown as a heat map (row: target class; column: epoch). Coefficients which were not significant (*P*_CORR_>0.05) are shown in white. **M**. Baseline choice-dependent change in influence, during the Early Cue, measured by Control target class stimulation (compare to Early Cue in Fig. 5D) **N**. Target class-dependent contributions to the choice-dependent shift in influence, during the Early Cue (compare to Fig. 5F). As in Fig. 5, error bars in panels M,N are the uncorrected 95% C.I.; asterisks indicate multiple comparisons corrected significance values (Bonferroni, 30X). To summarize, these results were consistent with the analyses that included imaging frames with stimulation and which were reported throughout this paper. On their own, these results become less conclusive as the data is increasingly subdivided (especially by both epoch and target class, panels L–N), because of the greatly reduced sampling.

**Figure S14.**
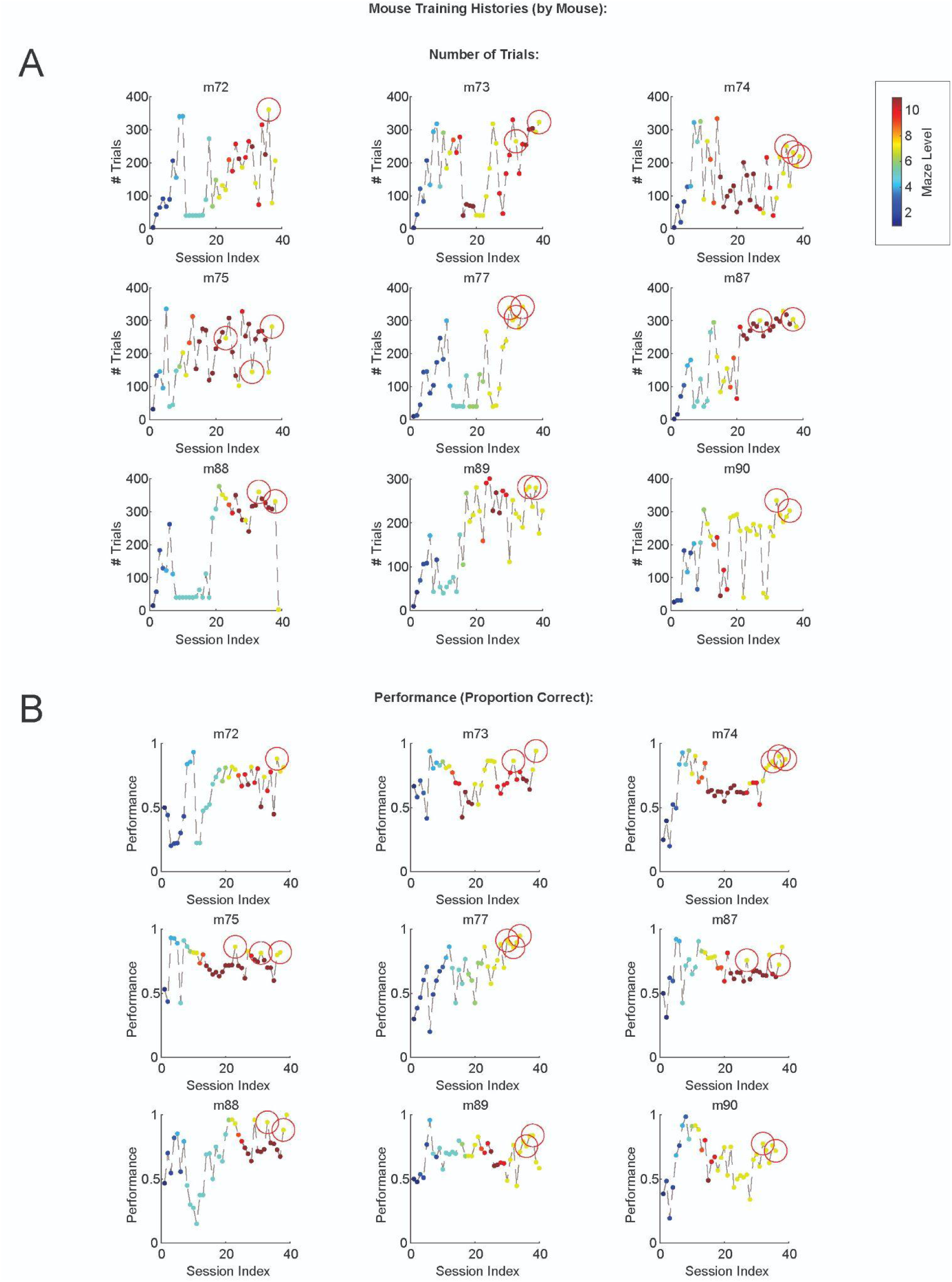
Training Histories of the Mice Involved in Single-Cell Perturbation Experiments, Related to Figure 1. Single-cell stimulation sessions are indicated with red circles. Mice were trained up to T11 (full difficulty ATT maze with distractors), though not all mice reached T11. All animals were trained primarily for a series of multi-neuron perturbation experiments, that have not yet been fully analyzed and which we plan to publish separately (not indicated on these panels). Single-cell stimulation experiments occurred after the multi-neuron perturbation experiments, at the end of the experimental round with the mice. These were performed in T7 (no-distractors version of the ATT maze). **B.** Number of trials run per session, maze level indicated by color. **C.** Performance (% Correct, over all trials, including warm up and aberrant trials) per session, maze level indicated by color.

---

**Table S1.**
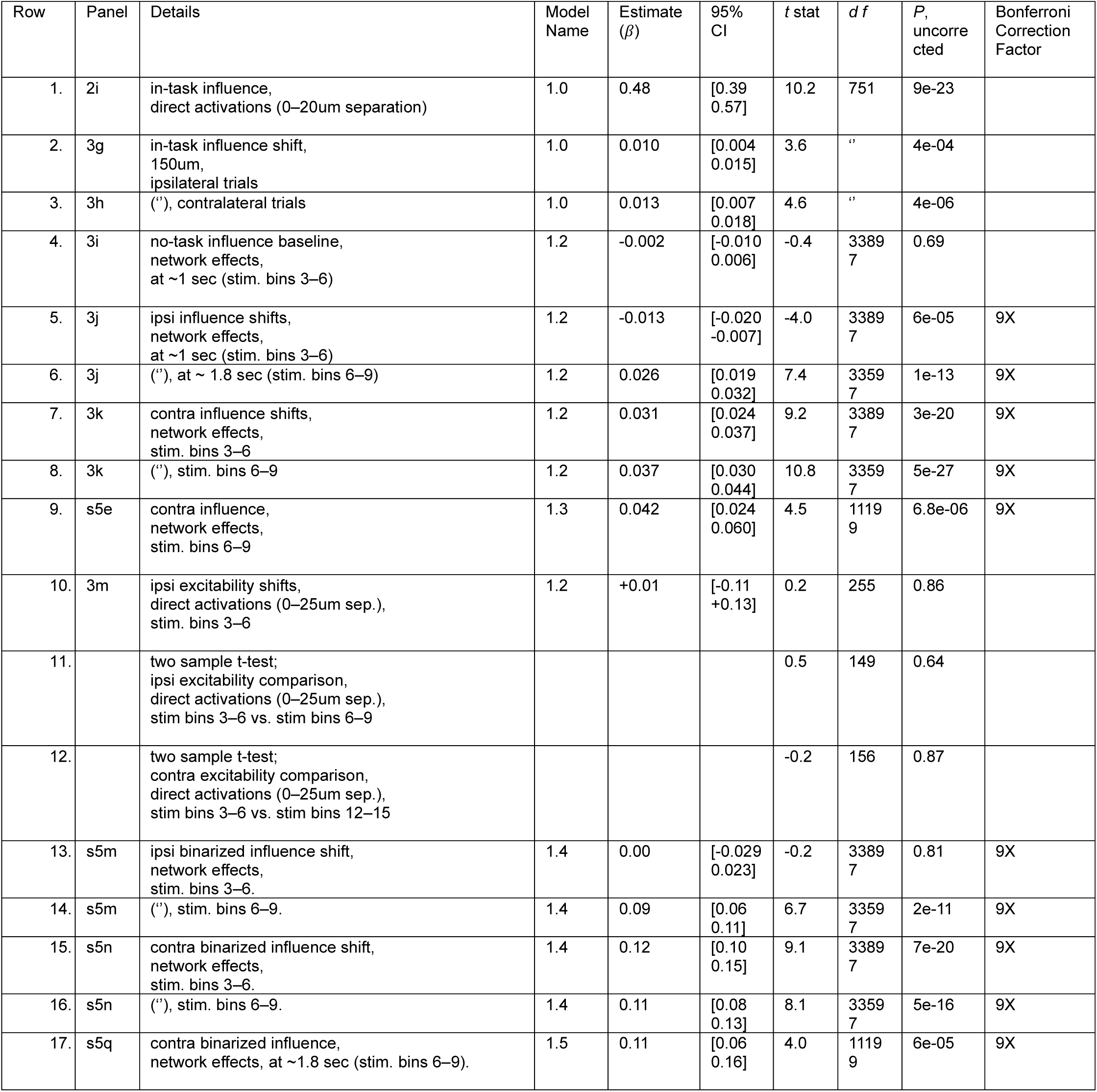
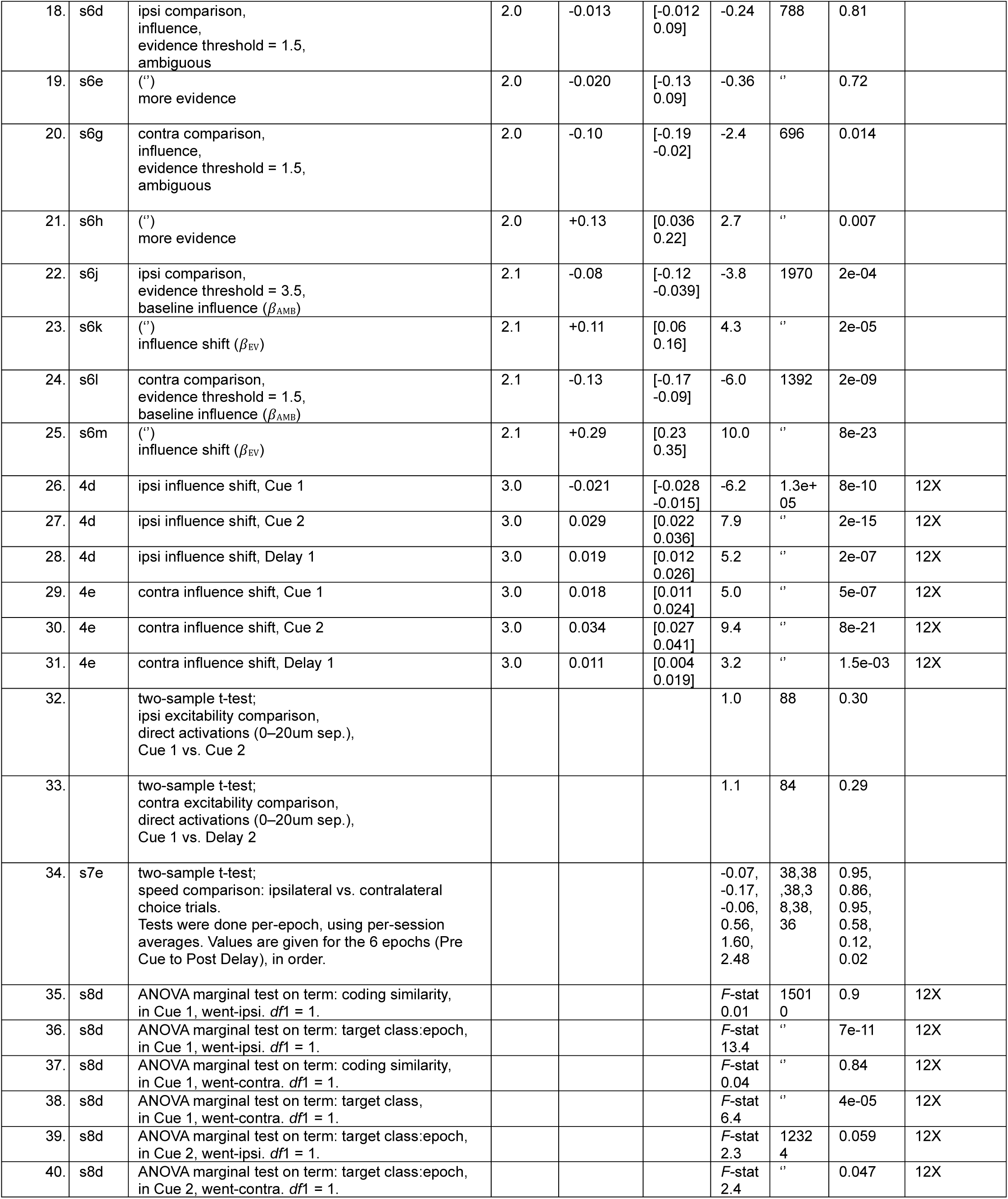
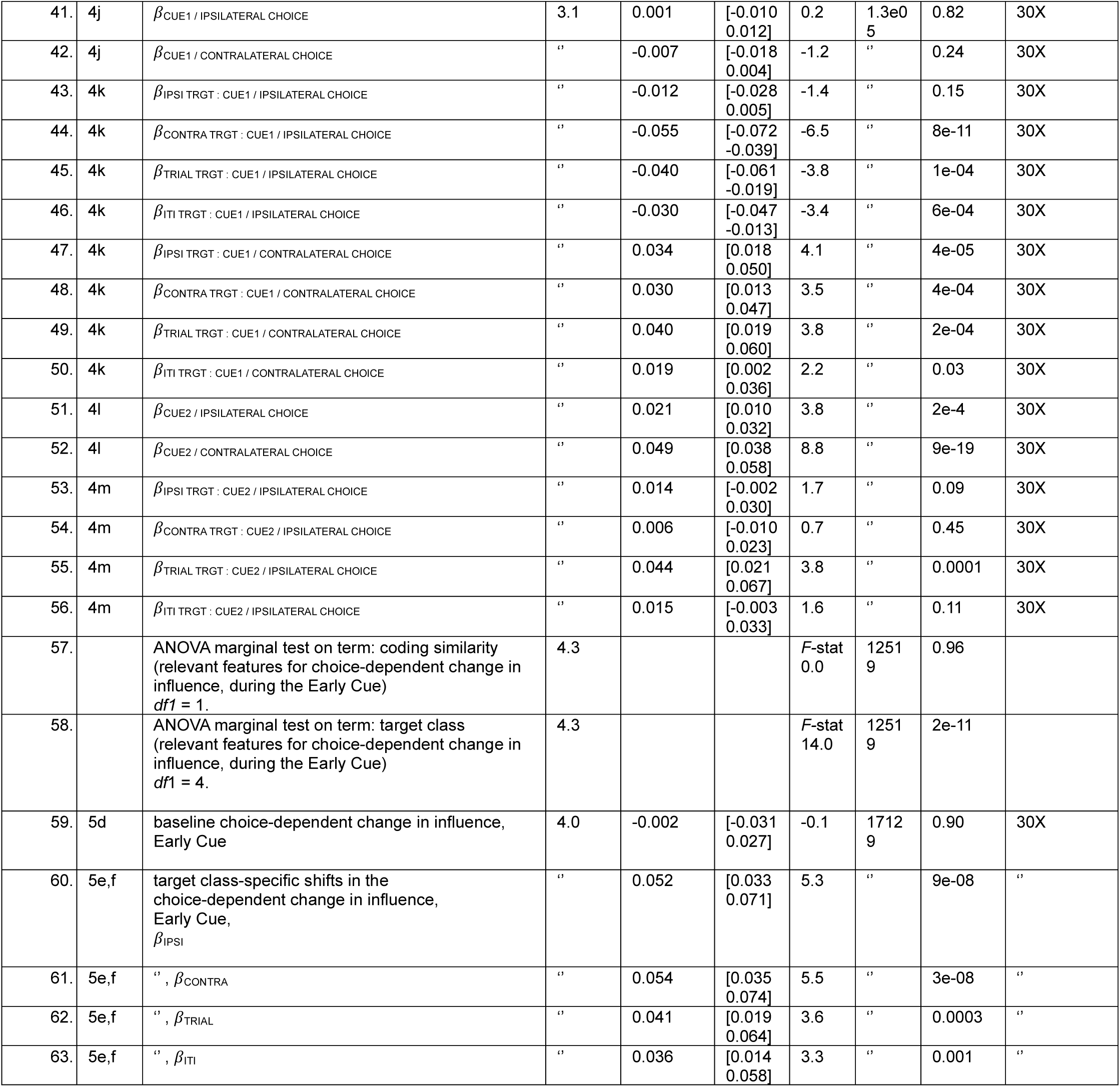
Mixed-effects models are indicated with model name (formulas, sample sizes and other details can be found in Table 1). All statistical comparisons in mixed-effects models are against a zero baseline. ANOVA marginal tests tested the significance of fixed-effects terms (across multiple coefficients), against a zero baseline. *P*-values reported in this table were not adjusted for multiple comparisons; correction factors are given here for the corrected significance values reported in the main text (indicated as P_CORR_).

## REFERENCES

1. Rickgauer, J. P., Deisseroth, K. & Tank, D. W. Simultaneous cellular-resolution optical perturbation and imaging of place cell firing fields. Nat. Neurosci. 17, 1816–1824 (2014).

2. Packer, A. M., Russell, L. E., Dalgleish, H. W. P. & Häusser, M. Simultaneous all-optical manipulation and recording of neural circuit activity with cellular resolution in vivo. Nat. Methods 12, 140–146 (2015).

3. Pégard, N. C. et al. Three-dimensional scanless holographic optogenetics with temporal focusing (3D-SHOT). Nat. Commun. 8, 1228 (2017).

4. Yang, W., Carrillo-Reid, L., Bando, Y., Peterka, D. S. & Yuste, R. Simultaneous two-photon imaging and two-photon optogenetics of cortical circuits in three dimensions. eLife 7, e32671 (2018).

5. Marshel, J. H. et al. Cortical layer–specific critical dynamics triggering perception. Science 365, eaaw5202 (2019).

6. Carrillo-Reid, L., Han, S., Yang, W., Akrouh, A. & Yuste, R. Controlling Visually Guided Behavior by Holographic Recalling of Cortical Ensembles. Cell 178, 447–457.e5 (2019).

7. Gill, J. V. et al. Precise Holographic Manipulation of Olfactory Circuits Reveals Coding Features Determining Perceptual Detection. Neuron 108, 382–393.e5 (2020).

8. Dalgleish, H. W. et al. How many neurons are sufficient for perception of cortical activity? eLife 9, e58889 (2020).

9. Daie, K., Svoboda, K. & Druckmann, S. Targeted photostimulation uncovers circuit motifs supporting short-term memory. Nat. Neurosci. 24, 259–265 (2021).

10. Oldenburg, I. A. et al. The logic of recurrent circuits in the primary visual cortex. Nat. Neurosci. 27, 137–147 (2024).

11. Chettih, S. N. & Harvey, C. D. Single-neuron perturbations reveal feature-specific competition in V1. Nature 567, 334–340 (2019).

12. Randi, F., Sharma, A. K., Dvali, S. & Leifer, A. M. Neural signal propagation atlas of Caenorhabditis elegans. Nature 623, 406–414 (2023).

13. Finkelstein, A., Daie, K., Rózsa, M., Darshan, R. & Svoboda, K. Connectivity underlying motor cortex activity during goal-directed behaviour. Nature 1–7 (2025) doi:10.1038/s41586-025-09758-6.

14. Pinto, L. et al. An Accumulation-of-Evidence Task Using Visual Pulses for Mice Navigating in Virtual Reality. Front. Behav. Neurosci. 12, (2018).

15. Koay, S. A., Charles, A. S., Thiberge, S. Y., Brody, C. D. & Tank, D. W. Sequential and efficient neural-population coding of complex task information. Neuron 110, 328–349.e11 (2022).

16. Brown, L. S. et al. Neural circuit models for evidence accumulation through choice-selective sequences. Preprint at 10.1101/2023.09.01.555612 (2023).

17. Diamanti, E. M. et al. Working memory expands shared task representations in cortex. 2025.09.29.679345 Preprint at 10.1101/2025.09.29.679345 (2025).

18. Pinto, L. et al. Task-Dependent Changes in the Large-Scale Dynamics and Necessity of Cortical Regions. Neuron 104, 810–824.e9 (2019).

19. LaFosse, P. K. et al. Bicistronic Expression of a High-Performance Calcium Indicator and Opsin for All-Optical Stimulation and Imaging at Cellular Resolution. eNeuro 10, (2023).

20. Pachitariu, M. et al. Suite2p: beyond 10,000 neurons with standard two-photon microscopy. 061507 Preprint at 10.1101/061507 (2017).

21. Bolkan, S. S. et al. Opponent control of behavior by dorsomedial striatal pathways depends on task demands and internal state. Nat. Neurosci. 25, 345–357 (2022).

22. Fahrmeir, L., Kneib, T., Lang, S. & Marx, B. D. Regression: Models, Methods and Applications. (Springer, Berlin, Heidelberg, 2021). doi:10.1007/978-3-662-63882-8.

23. Gauthier, J. L. et al. Detecting and correcting false transients in calcium imaging. Nat. Methods 19, 470–478 (2022).

24. Wong, K.-F. & Wang, X.-J. A Recurrent Network Mechanism of Time Integration in Perceptual Decisions. J. Neurosci. 26, 1314–1328 (2006).

25. Ko, H. et al. Functional specificity of local synaptic connections in neocortical networks. Nature 473, 87–91 (2011).

26. Sadeh, S. & Clopath, C. Theory of neuronal perturbome in cortical networks. Proc. Natl. Acad. Sci. 117, 26966–26976 (2020).

27. Yang, G. R., Joglekar, M. R., Song, H. F., Newsome, W. T. & Wang, X.-J. Task representations in neural networks trained to perform many cognitive tasks. Nat. Neurosci. 22, 297–306 (2019).

28. Sussillo, D. & Barak, O. Opening the Black Box: Low-Dimensional Dynamics in High-Dimensional Recurrent Neural Networks. Neural Comput. 25, 626–649 (2013).

29. Fino, E. & Yuste, R. Dense Inhibitory Connectivity in Neocortex. Neuron 69, 1188–1203 (2011).

30. Drinnenberg, A. et al. Large-scale cellular-resolution read/write of activity enables discovery of cell types defined by complex circuit properties. 2025.10.21.683734 Preprint at 10.1101/2025.10.21.683734 (2025).

31. Pfeffer, C. K., Xue, M., He, M., Huang, Z. J. & Scanziani, M. Inhibition of inhibition in visual cortex: the logic of connections between molecularly distinct interneurons. Nat. Neurosci. 16, 1068–1076 (2013).

32. Dipoppa, M. et al. Vision and Locomotion Shape the Interactions between Neuron Types in Mouse Visual Cortex. Neuron 98, 602–615.e8 (2018).

33. Pakan, J. M. et al. Behavioral-state modulation of inhibition is context-dependent and cell type specific in mouse visual cortex. eLife 5, e14985 (2016).

34. Ding, Z. et al. Functional connectomics reveals general wiring rule in mouse visual cortex. Nature 640, 459–469 (2025).

35. Kuan, A. T. et al. Synaptic wiring motifs in posterior parietal cortex support decision-making.

36. Cossell, L. et al. Functional organization of excitatory synaptic strength in primary visual cortex. Nature 518, 399–403 (2015).

37. Chen, I.-W. et al. High-throughput synaptic connectivity mapping using in vivo two-photon holographic optogenetics and compressive sensing. Nat. Neurosci. 1–13 (2025) doi:10.1038/s41593-025-02024-y.

38. Triplett, M. A. et al. Rapid learning of neural circuitry from holographic ensemble stimulation enabled by model-based compressed sensing. Nat. Neurosci. 1–12 (2025) doi:10.1038/s41593-025-02053-7.

39. Luo, T. Z. et al. Transitions in dynamical regime and neural mode during perceptual decisions. Nature 646, 1156–1166 (2025).

40. Bondy, A. G. et al. Brain-wide coordination of internal signals during decision-making. 2024.08.21.609044 Preprint at 10.1101/2024.08.21.609044 (2025).

41. Dombeck, D. A., Harvey, C. D., Tian, L., Looger, L. L. & Tank, D. W. Functional imaging of hippocampal place cells at cellular resolution during virtual navigation. Nat. Neurosci. 13, 1433–1440 (2010).

42. Gu, Y. et al. A Map-like Micro-Organization of Grid Cells in the Medial Entorhinal Cortex. Cell 175, 736–750.e30 (2018).

43. Botcherby, E. J., Juškaitis, R., Booth, M. J. & Wilson, T. An optical technique for remote focusing in microscopy. Opt. Commun. 281, 880–887 (2008).

44. Sofroniew, N. J., Flickinger, D., King, J. & Svoboda, K. A large field of view two-photon mesoscope with subcellular resolution for in vivo imaging. eLife 5, e14472 (2016).

45. Nikolenko, V., et al. SLM microscopy: scanless two-photon imaging and photostimulation using spatial light modulators. Front. Neural Circuits 2, (2008).

46. Grewe, B. F., Voigt, F. F., Hoff, M. van ’t & Helmchen, F. Fast two-layer two-photon imaging of neuronal cell populations using an electrically tunable lens. Biomed. Opt. Express 2, 2035–2046 (2011).

47. Friedrich, J., Zhou, P. & Paninski, L. Fast online deconvolution of calcium imaging data. PLOS Comput. Biol. 13, e1005423 (2017).

## REFERENCES

1. Chen, T.-W. et al. Ultrasensitive fluorescent proteins for imaging neuronal activity. Nature 499, 295–300 (2013).

2. Klapoetke, N. C. et al. Independent optical excitation of distinct neural populations. Nat. Methods 11, 338–346 (2014).

3. Baker, C. A., Elyada, Y. M., Parra, A. & Bolton, M. M. Cellular resolution circuit mapping with temporal-focused excitation of soma-targeted channelrhodopsin. eLife 5, e14193 (2016).

4. Marshel, J. H. et al. Cortical layer–specific critical dynamics triggering perception. Science 365, eaaw5202 (2019).

5. LaFosse, P. K. et al. Bicistronic Expression of a High-Performance Calcium Indicator and Opsin for All-Optical Stimulation and Imaging at Cellular Resolution. eNeuro 10, (2023).

6. Pachitariu, M. et al. Suite2p: beyond 10,000 neurons with standard two-photon microscopy. 061507 Preprint at 10.1101/061507 (2017).

7. Botcherby, E. J., Juškaitis, R., Booth, M. J. & Wilson, T. An optical technique for remote focusing in microscopy. Opt. Commun. 281, 880–887 (2008).

8. Friedrich, J., Zhou, P. & Paninski, L. Fast online deconvolution of calcium imaging data. PLOS Comput. Biol. 13, e1005423 (2017).

9. Bolkan, S. S. et al. Opponent control of behavior by dorsomedial striatal pathways depends on task demands and internal state. Nat. Neurosci. 25, 345–357 (2022).

